# A dynamical mechanism for opposing cortical gradients in resting-state functional and stimulation-derived effective connectomes

**DOI:** 10.64898/2026.11.24.768811

**Authors:** Xiaoyu Chen, Yixiao Feng, Boran Yang, Songting Li, Douglas Zhou

**Affiliations:** School of Mathematical Sciences, Shanghai Jiao Tong University, Shanghai, China.; Institute of Natural Sciences, Shanghai Jiao Tong University, Shanghai, China.; Ministry of Education Key Laboratory of Scientific and Engineering Computing, Shanghai Jiao Tong University, Shanghai, China.; Zhiyuan College, Shanghai Jiao Tong University, Shanghai, China.; State Key Laboratory of Synergistic Chem-Bio Synthesis, Shanghai Jiao Tong University, Shanghai, China.; Courant Institute School of Mathematics, Computing, and Data Science, New York University, New York, USA.

**Keywords:** functional connectome, intracranial electrical stimulation, effective connectome, wholecortex modelling, cortical hierarchy

## Abstract

Resting-state functional connectivity (FC) and stimulation-derived effective connectivity (EC) offer complementary views of cortical interactions, yet how cortical dynamics shape their relationship within a shared anatomical network remains unclear. Using group-level resting-state fMRI and intracranial electrical stimulation data from large-scale human cohorts, we find that regional mean response to electrical stimulation is negatively correlated with its mean FC (*r* = −0.568), with the two measures exhibiting opposing trends along the cortical hi-erarchy. To account for this divergence, we develop a differentiable whole-cortex mean-field model with independently trainable regional parameters, fitted to EC and FC through two-stage EC–FC training using gradient-based optimization. The optimized model reproduces EC (*r* = 0.733) and FC (*r* = 0.726) connectomes and their opposing regional signatures. Without prescribing a hierarchical parameter map, training yields a consistent organization in which recurrent strength and intrinsic noise increase from sensory to association cortex, whereas background input decreases. Linearization based analytical argument and parameter perturbation experiments further suggest how this organization can enhance evoked responsiveness while weakening resting-state correlations along the cortical hierarchy. Together, these findings link opposing connectome signatures, cortical hierarchy and emergent regional parameter organization, while providing a deep-learning-enabled, multimodal modelling framework for investigating cortical dynamics.

## Introduction

Resting-state functional connectivity (FC), measured as correlations between regional fMRI signals, is widely used to characterize spontaneous interactions across the cortex. Its spatial organization is constrained by structural connectivity (SC) (*1*), while also reflecting local dynamics and communication through indirect anatomical pathways (*2–4*). Intracranial electrical stimulation provides a complementary approach by probing how activity propagates in response to an externally imposed input. Cortico-cortical evoked potentials (CCEPs) elicited by single-pulse stimulation can be summarized as response probabilities, providing a directed measure of stimulationderived effective connectivity (EC) (*5–7*). Evoked responses between regions without a detected direct SC connection can be partially explained by multi-hop communication through the structural network, highlighting the contribution of network dynamics to stimulus propagation (*8, 9*). FC and stimulation-derived EC thus offer complementary views of cortical interactions arising from the same anatomical substrate and underlying dynamics, capturing spontaneous coordination and responsiveness to perturbations, respectively. However, how these two measurements relate to one another, and how cortical anatomy and dynamics jointly shape them, remain largely unclear.

An important biological coordinate for examining this relationship is provided by cortical hierarchy (*10, 11*). Along the axis from lower-order sensory-motor to higher-order association cortex, regions exhibit systematic variation in both structural and functional properties (*12–15*). At the structural and molecular levels, the myelin-sensitive T1w/T2w map is identified as a noninvasive neuroimaging index of hierarchy, in line with architectonic features (e.g., neuron size and spine count) and the principal component of genes and neurotransmitter receptors expression, particularly the interneuron markers parvalbumin (PV) and somatostatin (SST) for shaping microcircuit as well as excitation-inhibition (E-I) balance (*16–18*). At the functional level, intrinsic neural timescale generally increases toward higher-order cortex, linking hierarchical organization to differences in temporal integration (*12, 19–21*). This temporal hierarchy suggests a role for stronger recurrent dynamics in higher-order cortex (*21, 22*), consistent with the greater evoked excitability reported in higher-order networks (*23*). Together, these observations motivate examining FC and EC along the cortical hierarchy to clarify the role of regional differences in shaping spontaneous and evoked interactions.

Whole-cortex biophysical modelling offers a mechanistic approach to understanding how anatomical connectivity and neural dynamics jointly shape cortical interactions (*24, 25*). A whole-cortex dynamic mean-field model (MFM) was introduced in (*26*) that captures slow, NMDA-mediated population dynamics and reproduces key features of resting-state FC by coupling cortical regions through empirical SC. However, fitting resting-state FC alone may not fully constrain cortical dynamics, and combining FC with stimulation-derived EC provides a stronger basis for evaluating whether a common model can account for both spontaneous and evoked interactions. Integrating multimodal constraints within a single whole-cortex model nevertheless remains a frontier challenge in computational neuroscience (*27–32*). Advances in deep learning offer tools for addressing the challenge, particularly through automatic differentiation, that support the flexible design and integration of fitting objectives across data modalities (*33, 34*). Brain simulation platforms such as BrainPy (*35*) and The Virtual Brain (*36–38*) make the application of deep learning to whole-cortex model training increasingly practical, providing a foundation for multimodal training frameworks that combine FC and stimulation-derived EC to investigate the dynamical basis of cortical interactions.

In this study, we investigate how cortical dynamics jointly shape human FC and stimulation-derived EC. We compared group-level connectomes from the Human Connectome Project (HCP) resting-state fMRI data (*39, 40*) and the F-TRACT intracranial stimulation data (*5–7*). Despite their edge-level similarity as to SC, the node-level metrics show opposing EC and FC node-strength patterns: EC node strength (i.e., mean response to stimulus) increases along the hierarchy, whereas FC node strength decreases. To explain these patterns, we implemented the whole-cortex MFM as a recurrent neural network (RNN) in BrainPy, and developed a connectome-based, cross-modality, endto-end optimization framework using back-propagation through time (BPTT), allowing regional parameters to be optimized independently under EC and FC constraints. The fitted models reproduce both connectomes and their opposing regional patterns. Without a prescribed hierarchical parameter map, hierarchy emerges in the model parameters, with recurrent strength and intrinsic noise increasing from sensory to association cortex and background input decreasing. Furthermore, through theoretical analyses and computational experiments, we demonstrated how EC and FC node strength relate to regional parameters of local recurrence and intrinsic noise. These findings provide a candidate dynamical mechanism for the opposing EC and FC patterns, together with a computational framework that opens an avenue for deep-learning-enabled multimodal whole-cortex modelling.

## Results

### Node strength divergence between empirical EC and FC along the axis of cortical hierarchy

We first investigated EC and FC at edge level, particularly their relationship to the underlying SC. Both EC (*r* = 0.515, p < 0.05, Fig. 1A right panel) and FC (*r* = 0.394, p < 0.05, Fig. 1B right panel) showed positive correlations with SC, particularly between anatomically connected regions (EC: *r* = 0.420, p < 0.05, Fig. 1A right panel; FC: *r* = 0.523, p < 0.05, Fig. 1B right panel). Anatomically connected pairs of regions also exhibited significantly higher EC (p < 0.05, Fig. 1A left panel) and FC (p < 0.05, Fig. 1B left panel) compared to anatomically unconnected pairs. The result reveals that both EC and FC are constrained by, and converge toward SC as the underlying backbone. However, despite reaching significance, the correlation between EC and FC remained modest (*r* = 0.196; Fig. 1C), suggesting an underlying divergence between the two modalities. We then investigated the connectome data at node level using node strength (NS), a standard graphtheoretic metric that summarizes the weights of connections of each node (*41*). This metric has been widely used to characterize brain network organization (*42–44*), development and aging (*45, 46*) and disease-related functional alterations (*47, 48*). SC-NS and FC-NS correspond to the mean SC and FC weights for each node, respectively. Note that EC-NS included both afferent (i.e., mean response of the node when stimulating other nodes) and efferent (i.e., mean impact of the node on other nodes when stimulating the node) versions due to the directed nature of EC (see Fig. 2A for detailed definitions and illustration). Pairwise correlations were computed among the abovedefined node strengths and between these node strengths and cortical gradients of T1w/T2w ratio, PV expression and SST expression (Fig. 2B,C). SC-NS was positively correlated with FC-NS but showed no significant correlation with EC-NS (Fig. 2C). Surprisingly, a significant negative correlation was found between EC-NS (afferent) and FC-NS (*r* = −0.568, FWE-corrected p < 0.05, Fig. 2C,D). This negative relationship results in a paradoxical pattern: EC and FC edge weights are generally positively correlated within groups defined by their target (afferent for EC) or source (efferent for EC) nodes (Supplementary Fig. S1A,B), yet pooling all edges yields a weaker EC-FC correlation than the mean within-node correlation (Supplementary Fig. S1C,D). Along the axis of cortical hierarchy, SC-NS showed no significant correlation with cortical gradients, whereas FCNS and EC-NS exhibited significant but opposite trends (Fig. 2C). Taking SST as an instance for cortical hierarchy, SC-NS showed no significant correlation (*r* = −0.04, Fig. 2E top panel), while FC-NS showed a significant negative correlation (*r* = −0.572, FWE-corrected p < 0.05, Fig. 2E middle panel) and EC-NS (afferent) showed a significant positive correlation (*r* = 0.681, FWEcorrected p < 0.05, Fig. 2E bottom panel). These results suggest that hierarchy-related information is embedded in the node strength of EC and FC, with opposing patterns: EC-NS increases while FC-NS decreases along the cortical hierarchy.

**Fig. 1:**
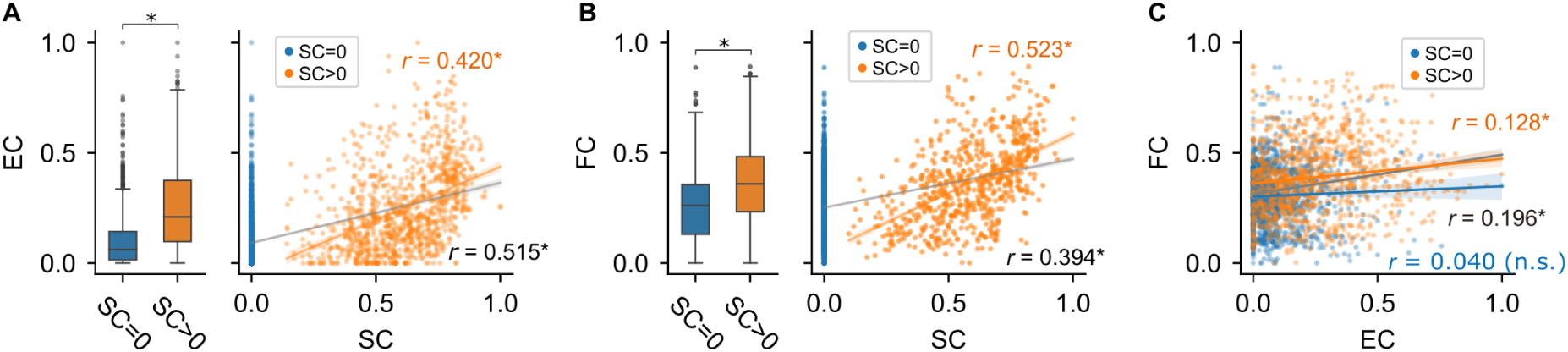
Edge-level relationships among empirical structural, effective and functional connectomes. **A** Left: effective connectivity (EC) between structurally unconnected (SC = 0, blue) and connected (SC > 0, orange) pairs of cortical regions. Right: EC versus structural connectivity (SC), with regression lines for structurally connected pairs (orange) and all measured pairs (grey). **B** The same comparisons for functional connectivity (FC). **C** FC versus EC, with regression lines for structurally connected pairs (orange), structurally unconnected pairs (blue) and all measured pairs (grey). Each point in the scatter plots represents a connection. Boxplots in **A,B** compare the two structural-connectivity groups (*, *p* < 0.05, two-sample *t*-test). Correlation coefficients in **A–C** are Pearson’s *r* (*, *p* < 0.05; n.s., not significant).

**Fig. 2:**
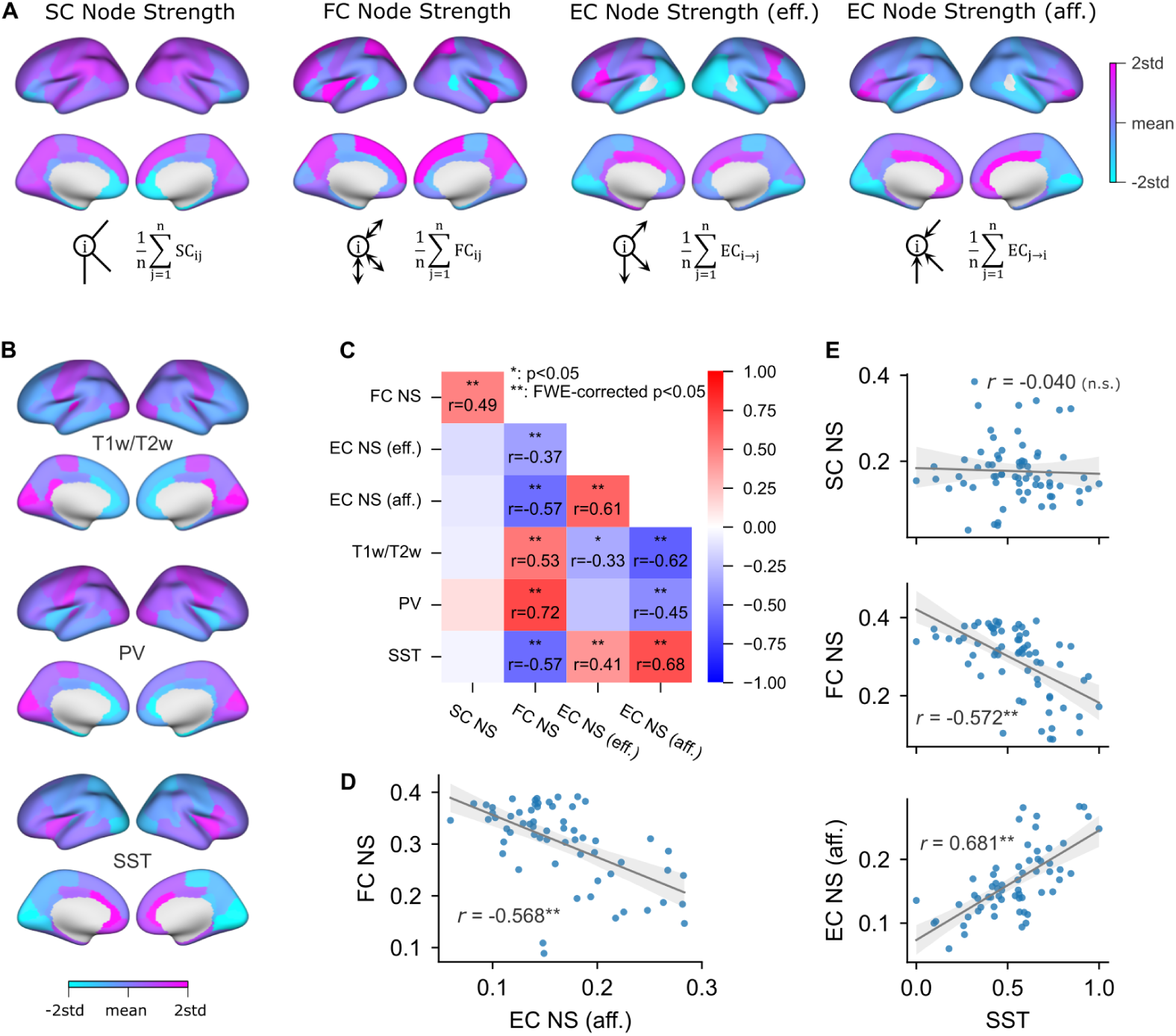
Opposing relationships of empirical EC and FC node strengths with cortical hierarchy. **A** Cortical maps and definitions of node strength (NS) for SC, FC, efferent EC and afferent EC. NS is the mean weight of all available connections of each node; efferent and afferent EC quantify its mean influence on other nodes and its mean response to stimulation of other nodes, respectively. Self-connections and unmeasured EC entries are excluded (see Materials and Methods for details). **B** Cortical maps of T1w/T2w ratio and parvalbumin (PV) and somatostatin (SST) gene expression. The colour scales in **A,B** span two standard deviations below and above each map’s mean. **C** Pairwise Pearson correlations among node strengths and cortical gradients. Asterisks indicate significance (*, *p* < 0.05; **, family-wise error (FWE)-corrected *p* < 0.05). **D** Negative correlation between afferent EC NS and FC NS. **E** Relationships of SST expression with SC NS (top), FC NS (middle) and afferent EC NS (bottom). Points in **D,E** denote cortical regions; grey lines show linear fits. Significance notation follows **C**; n.s., not significant.

We further validated the opposing EC and FC NS patterns by following control analyses: the findings are robust to an alternative FC from (*49*) with independent fMRI preprocessing and subject selection procedures (Supplementary Fig. S2A–C). Using the HCP-MMP parcellation (*50*) (Supplementary Fig. S2D), we replicated both the negative correlation between EC-NS (afferent) and FC-NS and their opposing relationships with cortical hierarchy (all p < 0.001; Supplementary Fig. S2E,F). Given that Euclidean distance (ED) was significantly embedded in both EC and FC

(Supplementary Fig. S2G,H), NS was recomputed using ED-regressed EC and FC (Supplementary Fig. S2I), with a replicated negative EC–FC NS correlation (*r* = −0.411, p < 0.001; Supplementary Fig. S2J) and their correlations as to cortical hierarchy, e.g., *r* = 0.710 for EC-NS (afferent) vs. SST and *r* = −0.602 for FC-NS vs. SST (Supplementary Fig. S2K). We further validated the robustness of the opposing EC and FC NS patterns by recomputing node strength separately for SC-connected and SC-unconnected edges, showing highly consistent NS estimates across different computational definitions (Supplementary Fig. S3A). The negative relationship between ECand FC-NS persisted regardless of whether NS was computed using SC-connected or SC-unconnected edges (Supplementary Fig. S3B).

### A cross-modality framework for end-to-end gradient-based training of a wholecortex MFM

To explain how the opposing EC and FC NS patterns could arise from a shared structural network, we used a whole-cortex MFM, which comprises 68 interconnected regions (nodes), each corresponding to a cortical parcel in the Desikan-Killiany (DK) parcellation (*26, 51*). Regions interact through long-range excitatory projections whose weights are defined by a diffusion MRI derived, group-averaged SC from the Human Connectome Project (HCP) (*39*), and globally scaled by parameter *G* (Fig. 3A). At region with index *i* (*i* = 1, 2,…, 68), excitatory neuronal population forms a local microcircuit whose dynamics are governed by recurrent connection (*w_i_*), background input (*I_i_*) and intrinsic noise level (*σ_i_*). Note that *w* is relaxed to allow negative values to represent net feedback inhibition (see Supplementary Fig. S4 and Supplementary Note 1). Accordingly, the model parameter space spans a sufficiently broad range to encompass a wide variety of dynamical regimes that facilitate signal propagation, analogous to the global balanced amplification (GBA) regime (*52*). Such a regime of brain-wide dynamics enables signal propagation while maintaining system stability, with strong inter-regional excitation balanced by intra-regional recurrent inhibition (*52*). This extension of parameter space is validated as necessary to enable network signal propagation and achieve optimal model-empirical fit, particularly for indirect edges in EC (Supplementary Fig. S5). Throughout this and following paragraphs, we use the plain notation (e.g., *w*) to represent the regionally homogeneous setting, where all regional parameters share a single common value, while a subscript or vector symbol (e.g., *w_i_* or *w⃗*) denotes the regionally heterogeneous setting, where region’s parameters are independent of each other. For illustration, the parameter space can be marked by a bifurcation line which delineates a necessary condition for bistability in the regionally homogeneous, noise-free MFM dynamics (see Supplementary Fig. S6 and Supplementary Note 2).

**Fig. 3:**
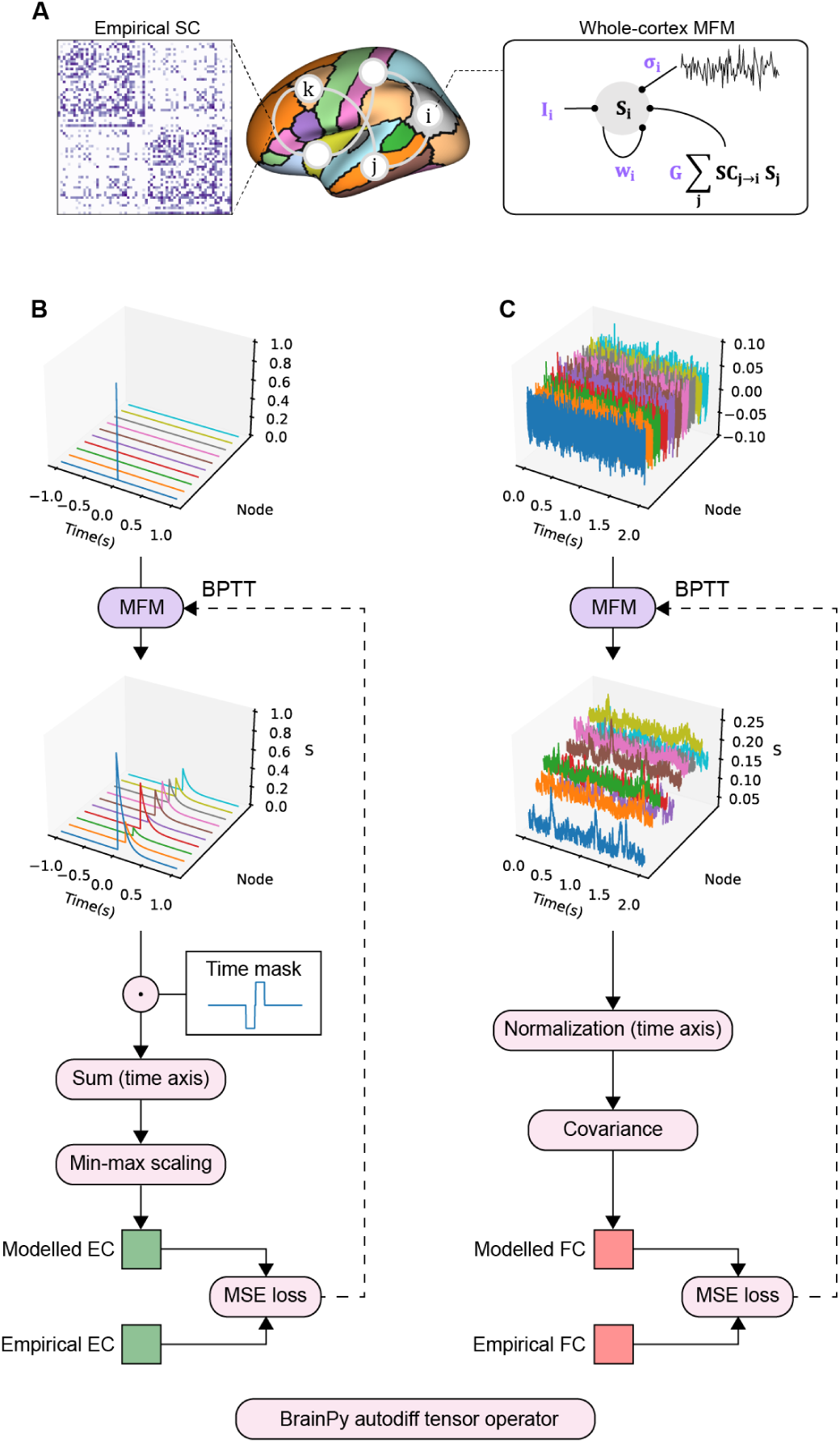
Connectome-based, end-to-end gradient-based training of a whole-cortex MFM. **A** The whole-cortex mean-field model represents each of 68 Desikan-Killiany cortical regions by a synaptic gating variable *S_i_*. Inter-regional interactions follow empirical SC and are scaled by global coupling *G*. Regional recurrent strength *w_i_*, background input *I_i_* and intrinsic noise amplitude *σ_i_* are independently trainable. **B** EC fitting. A single-pulse input is delivered to one node of the model, implemented as an RNN, and responses are recorded across nodes. A temporal mask combines pre-stimulus baseline subtraction and post-stimulus integration; summation over time and min-max scaling produce modelled EC. Back-propagation through time (BPTT) minimizes the mean squared error (MSE) between modelled and empirical EC to optimize *G*, *w_i_* and *I_i_*. **C** FC fitting. Noise inputs drive a virtual resting-state simulation. Normalization of each regional time series followed by covariance calculation yields Pearson-correlation-based FC. BPTT minimizes the FC MSE and additionally enables optimization of *σ_i_*. All operations are implemented with differentiable BrainPy tensor operators.

To train this model, we developed a cross-modality framework that optimizes regional parameters independently by fitting fMRI-derived, group-averaged FC from HCP (*39, 40*) and iEEGderived, group-aggregated EC from F-TRACT (*5–7*). The framework implements the MFM as an RNN in BrainPy, applies autodiff tensor operators to obtain modelled EC and FC, and performs optimization in an end-to-end fashion. Virtual stimulus-response trial is conducted by injecting a single-pulse stimulus into the targeted node, recording the RNN response, and calculating modelled EC as the baseline corrected area under the evoked response curve (see Fig. 3B and Materials and Methods for details). Virtual resting-state trial is conducted by injecting Gaussian noises into the RNN and calculating modelled FC as Pearson’s correlation between time series of the responses (see Fig. 3C and Materials and Methods for details). BPTT with Adam optimizer minimizes the mean squared error (MSE) between modelled and empirical EC to train the parameters G, *w_i_* and *I_i_*, while FC fitting additionally optimizes the noise parameter *σ_i_*. As a validation of trainability, we tested EC fitting and FC fitting on synthetic EC and FC matrices derived using the same MFM with predefined ground-truth parameters. In regionally homogeneous cases (a single shared *w*, *I* and *σ* across cortical regions), the fitting procedures could bring parameter estimates closer to the ground-truth values from their initial settings (Supplementary Fig. S7). In regionally heterogeneous cases, we introduced regional heterogeneity by adding a random vector to *w_i_* or *σ_i_* when generating synthetic EC or FC matrices, respectively, and examined whether fitting could recover the imposed regional patterns (Supplementary Fig. S8). Together, these tests support the trainability of the framework by showing its ability to recover model parameters and imposed regional heterogeneity across a range of parameter initializations and settings. We selected the hyperparameters for EC and FC fitting using five-fold cross-validation, which supported generalization to held-out connections and showed consistent modelled connectomes and training trajectories across folds (Supplementary Fig. S9). These results also suggest that the model may help address the incomplete coverage of EC measurements by providing estimates for empirically unobserved connections (Supplementary Fig. S10).

Based on this framework, we implemented three training strategies to assess how EC and FC constrain regional dynamics. In two-stage EC–FC training, EC fitting first determined global coupling, regional recurrent strength and background input; subsequent FC fitting estimated regional intrinsic noise while keeping the EC-trained parameters fixed. This strategy therefore relied primarily on EC to constrain network dynamics, with FC providing complementary information about spontaneous fluctuations. For comparison, FC-only training consisted of a single stage of FC fitting, in which all model parameters were optimized against FC alone, testing whether resting-state information was sufficient to recover both connectomes and their regional organization. Hybrid EC–FC training consisted of a single stage of joint EC and FC fitting with the weighted objective ℒ_hybrid_ = *α*ℒ_EC_ + (1 −*α*)ℒ_FC_, where ℒ_EC_ and ℒ_FC_ denote the EC and FC MSE losses, respectively, and the EC–FC loss ratio *α* controls their relative contributions. This strategy allowed us to test how the balance of EC and FC constraints influenced model fits and parameter organization. Detailed training settings are provided in Materials and Methods and Supplementary Table S2.

### High-fidelity EC and FC simulations with emergent model parameter regime and hierarchical heterogeneity

We first examine the two-stage training process by projecting the training trajectories onto the (*G*, *w̅*, *I̅*) axes, where *w̅* and *I̅* denote the mean values of *w_i_* and *I_i_* across all regions, respectively. Two-stage training was initialized using a broad range of regionally homogeneous seeds, which span the axis of *w̅* (Fig. 4A, two-dimensional projection; Supplementary Fig. S11, full threedimensional trajectories). The training trajectories with optimized losses converged onto a narrow regime of positive *G* and negative *w̅* under the bifurcation line (Fig. 4A), similar to the GBA regime in which a combination of strong inter-regional excitation coupling and intra-regional recurrent inhibition supports signal propagation while maintaining stable dynamics (*52*). We quantified model performance using the goodness of fit, i.e., Pearson’s correlation r-value between modelled and empirical entries of EC or FC matrix. For FC, only upper triangular entries were considered due to its symmetry. The 26 retained models from two-stage training were defined by jointly achieving high model-empirical fits (r > 0.6) for both EC and FC (Fig. 4B).

**Fig. 4:**
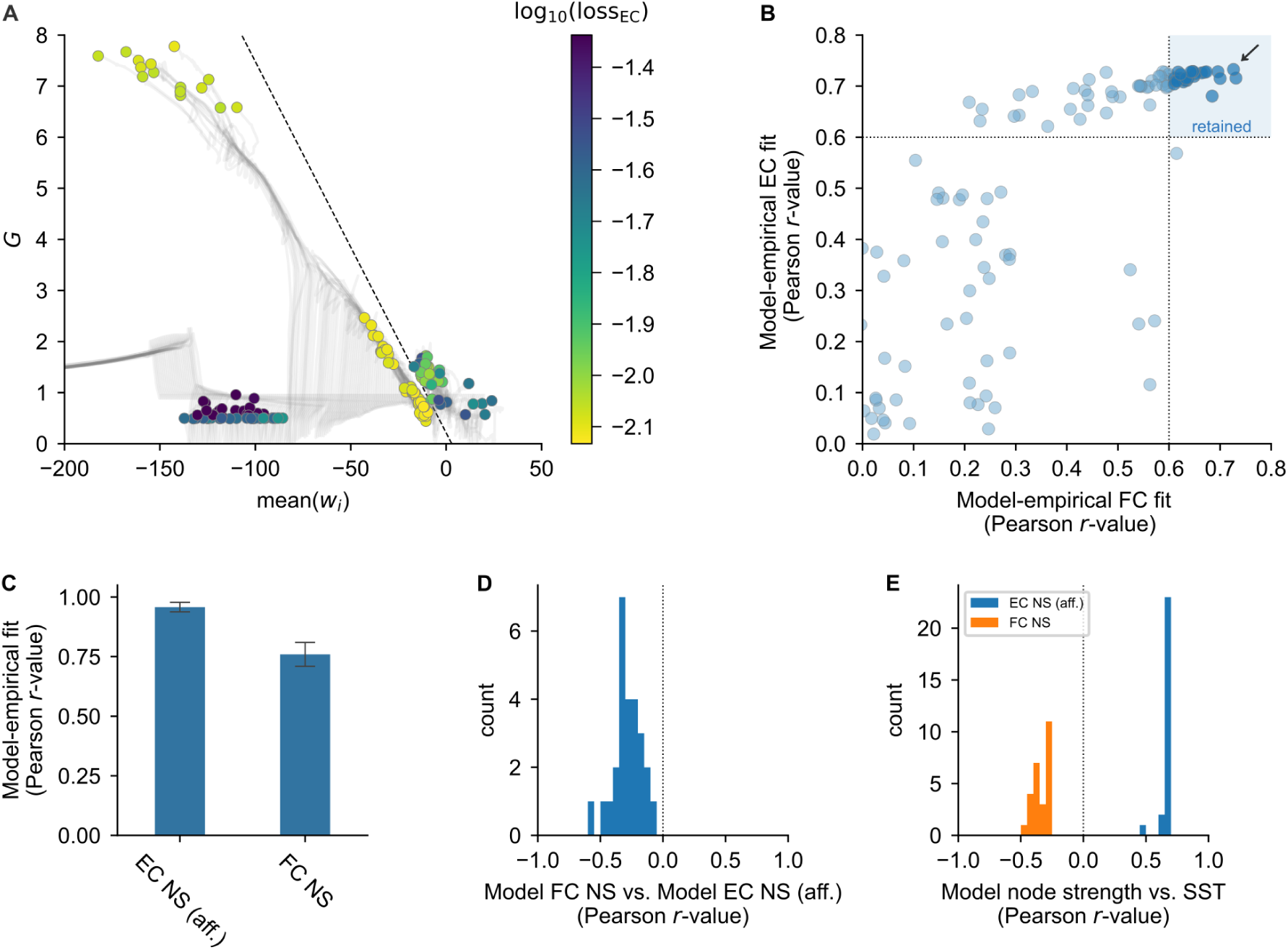
Two-stage training reproduces edge-level fits and opposing node-strength patterns. **A** EC-fitting trajectories projected onto global coupling *G* and mean(*w_i_*). Grey curves denote training trajectories; coloured circles mark their endpoints and encode log_10_(loss_EC_). The dashed line denotes the homogeneous-model bifurcation reference, shown for illustrative purpose in the projected parameter space (see Supplementary Note 2 for details). **B** Model-empirical FC and EC fits after two-stage training, quantified by Pearson’s *r*. The shaded region retains 26 models with *r* > 0.6 for both modalities; the arrow identifies the optimal model. **C** Mean model-empirical fits for afferent EC NS and FC NS across the 26 retained models from two-stage training; error bars indicate standard deviations. **D** Distribution of Pearson correlations between modelled FC NS and afferent EC NS across the same 26 retained models. **E** Distributions of Pearson correlations between SST expression and modelled afferent EC NS (blue) or FC NS (orange) across the same 26 retained models.

As the 26 retained models achieve edge-level fit to both EC and FC (Fig. 4B), we verified that they can straightforwardly reproduce the above empirical findings at the node level, especially the opposing EC and FC NS patterns. These 26 retained models captured the node strength patterns with significant model-empirical FC-NS fit (*r* = 0.76 ± 0.05) and EC-NS (afferent) fit (*r* = 0.96 ± 0.02, Fig. 4C), and replicated the negative correlation between EC-NS (afferent) and FC-NS (Fig. 4D). The modelled afferent EC NS and FC NS also showed opposite relationships with SST expression across these 26 retained models (Fig. 4E).

We validated whether regional heterogeneity of model parameters is vital for reproducing the opposing EC and FC NS patterns, by assessing regionally homogeneous model with parameters *w*, *I* and *σ* set uniformly across cortical regions. Across the full grid search, homogeneous models failed to simultaneously reproduce afferent EC-NS and FC-NS patterns (Supplementary Fig. S12B). We selected a subset with good edge-level fits to both EC and FC (see Materials and Methods; Supplementary Fig. S12A), and these retained homogeneous models failed to reproduce the negative correlation between afferent EC-NS and FC-NS (Supplementary Fig. S12C). These results provide simulation-based support for the necessity of regional heterogeneity in reproducing the opposing EC and FC NS patterns.

We then examined the optimally-trained case, which yielded high-fidelity simulations of both EC (*r* = 0.733, p < 0.05, Fig. 5A,B) and FC (*r* = 0.726, p < 0.05, Fig. 5C,D). See Supplementary Fig. S13 for modelled time courses of resting-state BOLD and the single-pulse stimulation evoked responses. Beyond the model-empirical fits of EC (*r* = 0.733, p < 0.05, Fig. 5B) and FC (*r* = 0.727, p < 0.05, Fig. 5D) between anatomically connected regions, the model dynamics captured brainwide network indirect effects arising from multi-hop propagation. Significant correlations were observed between modelled and empirical EC (*r* = 0.487, p < 0.05, Fig. 5B) or FC (*r* = 0.698, p < 0.05, Fig. 5D) as to anatomically unconnected pairs of regions. At the node level, model-empirical fits for EC-NS (afferent) and FC-NS reached *r* = 0.97 and *r* = 0.86, respectively (Fig. 5E,F), with a significant negative correlation between modelled EC-NS (afferent) and FC-NS (*r* = −0.435, p < 0.05, Fig. 5G). This negative relationship was consistent with the empirical finding (Fig. 2D). In this model, afferent EC NS and FC NS were positively and negatively correlated with SST expression, respectively (Fig. 5H).

**Fig. 5:**
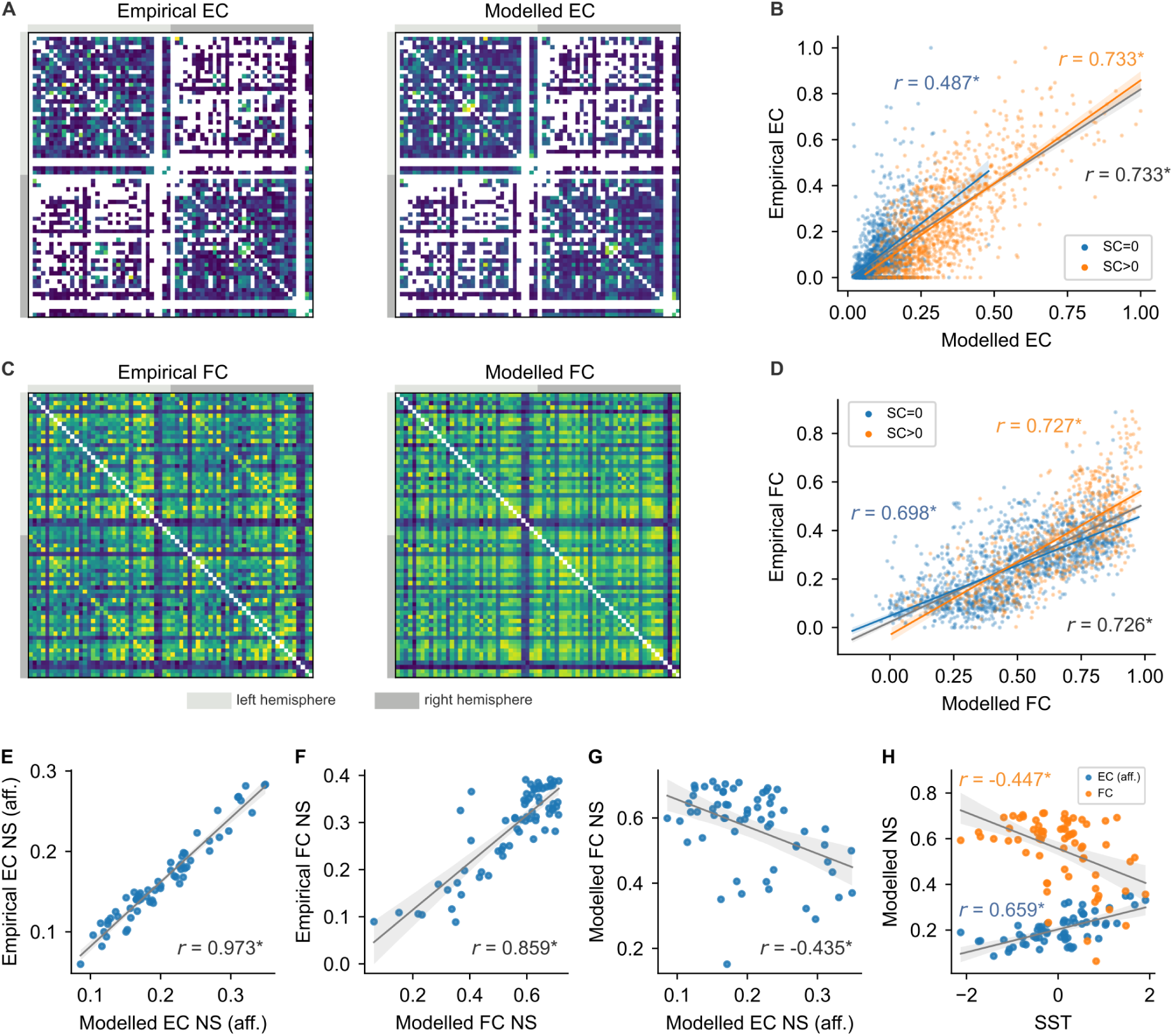
Optimal model captures EC and FC at both edge and node levels. **A** Empirical and modelled EC matrices for the optimal model obtained through two-stage training. White entries indicate masked connections; the empirical measurement mask is also applied to the displayed modelled EC. **B** Empirical versus modelled EC for structurally connected (orange), structurally unconnected (blue) and all measured (grey regression line) pairs of regions. **C,D** Corresponding FC matrices and edge-level comparisons. Light and dark grey strips beside the matrices identify left- and right-hemisphere regions. **E,F** Empirical versus modelled afferent EC NS (**E**) and FC NS (**F**). **G** Negative relationship between modelled afferent EC NS and FC NS. **H** Relationships of SST expression with modelled afferent EC NS (blue) and FC NS (orange). Points in **E–H** represent cortical regions. Lines show linear fits; annotations report Pearson’s *r* (*, *p* < 0.05).

Given the strong model-empirical fits and the failure of homogeneous models to reproduce the opposing NS patterns, we next tracked regional parameter heterogeneity throughout training of the optimal model. Across the EC-fitting and FC-fitting stages (Supplementary Fig. S14A,B), the loss rapidly decreased and stabilized onto a plateau, while regional differences in parameters developed alongside systematic correlations with cortical hierarchy (i.e., *w_i_* and *σ_i_* positively, while *I_i_* negatively, correlated with hierarchy). See Supplementary Fig. S14C and Fig. 2B for an illustration of the trained regional parameters as well as the cortical gradients.

To assess whether this emergent hierarchical pattern was consistent across the 26 retained models, we examined how regional parameters (*w_i_*, *I_i_* and *σ_i_*) correlated with myelin and gene expression gradients (T1w/T2w ratio, PV and SST expression). Robust patterns were observed: both *w_i_* and *σ_i_* were positively correlated with SST and negatively correlated with T1w/T2w and PV, aligning positively along the axis of cortical hierarchy; while *I_i_* exhibited inverse patterns of correlation (FWE-corrected p < 0.05, Fig. 6A,B), consistent with previous findings (*49*).

**Fig. 6:**
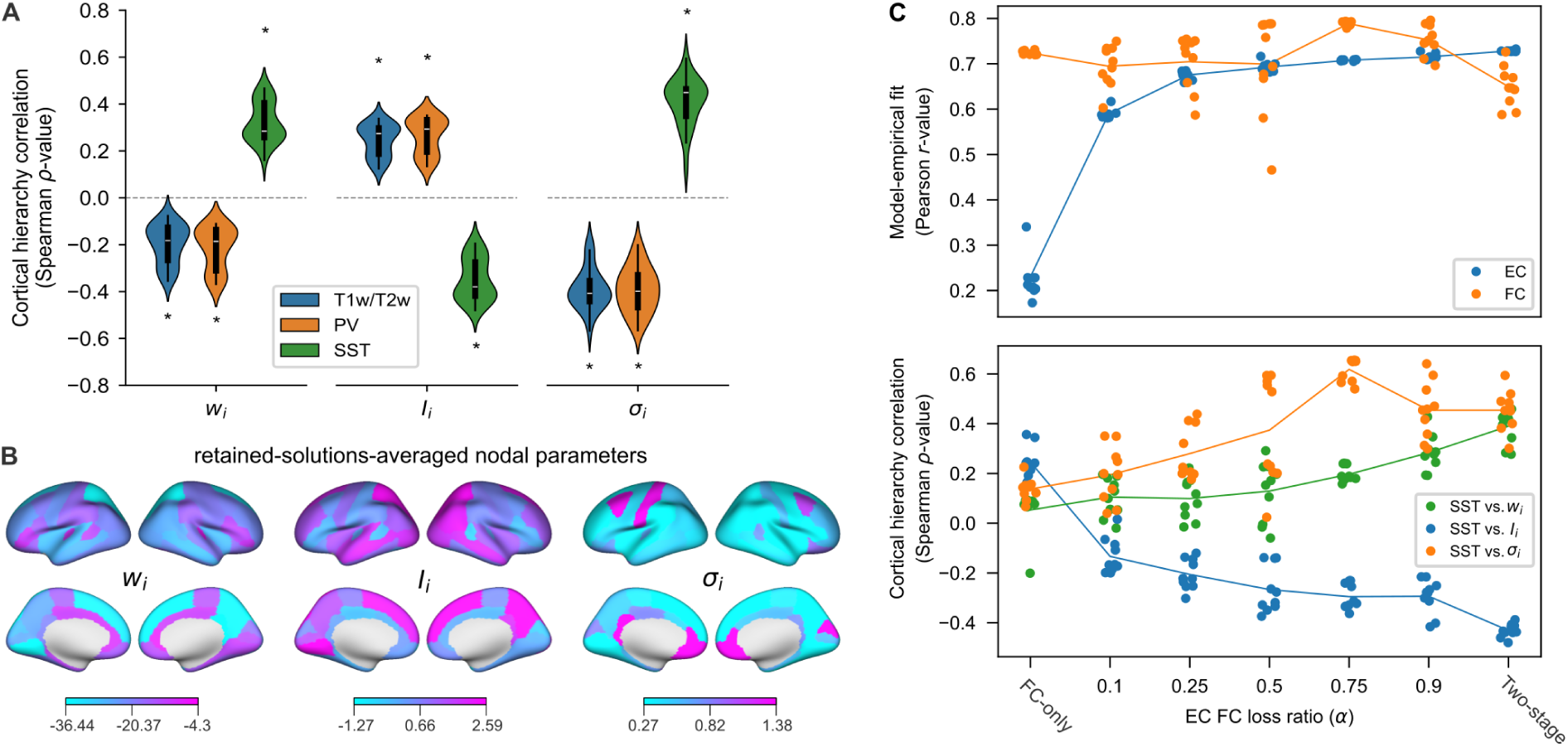
Hierarchical organization of trained parameters and its dependence on EC and FC constraints. **A** Distributions of Spearman correlations between regional parameters (*w_i_*, *I_i_*, *σ_i_*) and T1w/T2w (blue), PV (orange) and SST (green) across the 26 retained models from two-stage training. Asterisks denote FWE-corrected *p* < 0.05 in one-sample *t*-tests against zero. **B** Cortical maps of regional parameters averaged across the same 26 retained models; colour scales show parameter values. **C** Comparison of FC-only training, hybrid EC–FC training and two-stage training. Hybrid training minimizes ℒ_hybrid_ = *α*ℒ_EC_ + (1 − *α*)ℒ_FC_, with loss ratio *α* ∈ {0.1, 0.25, 0.5, 0.75, 0.9}. Top: model-empirical EC (blue) and FC (orange) fits, measured by Pearson’s *r*. Bottom: Spearman correlations of SST expression with *w_i_* (green), *I_i_* (blue) and *σ_i_* (orange). Dots denote the ten models with the lowest final training losses for each pipeline; lines connect their means. FC-only training and two-stage training are reference conditions, and *α* applies only to hybrid training.

To determine how including EC as a training constraint contributed to this parameter organization, we compared these results with FC-only training, which jointly optimized *G*, *w_i_*, *I_i_* and *σ_i_* against FC alone in a single stage. FC-only training also converged toward the bifurcation line (Supplementary Fig. S15A). However, it only achieved high model-empirical fits (r > 0.6) for FC in a subset of well-trained models (n = 89), and failed to achieve considerable model-empirical fits for EC (Supplementary Fig. S15B). In addition, regional parameters of these well-trained models exhibited robust but distinct relationships with cortical hierarchy. While *σ_i_* remained positively correlated with cortical hierarchy, same as in two-stage training, the patterns of *w_i_* and *I_i_* were inversed showing negative and positive correlations with cortical hierarchy, respectively (FWE-corrected p < 0.05, Supplementary Fig. S15C), replicating the inverted gradient finding as reported by (*53*).

To further test dependence of trained regional parameters on EC constraints, we used hybrid EC–FC training to vary the EC–FC loss ratio *α* in the joint objective (see Materials and Methods). This comparison tested whether the contrasting outcomes of FC-only and two-stage training varied gradually with the balance of EC and FC constraints. Training followed similar trajectories across different *α* values, consistently converging toward the regime near the bifurcation line (Supplementary Fig. S16). Increasing the contribution of EC progressively improved model-empirical EC fit, while relatively high FC fit was still retained across a broad range of *α* values (Fig. 6C top panel). The parameter–hierarchy relationships also varied with the relative EC and FC constraints: as *α* increased, *w_i_* tended to become more positively correlated with SST expression and *I_i_* more negatively correlated, whereas *σ_i_* showed positive correlations throughout (Fig. 6C bottom panel).

To distinguish the contributions of recurrence and noise heterogeneity to these training outcomes, we next tested mixed homogeneous/heterogeneous parameterizations (Supplementary Fig. S17A). Under two-stage training, which fitted both EC and FC (Fig. 4B), constraining *σ* to be homogeneous significantly reduced model-empirical FC fit (Supplementary Fig. S17B,C). Under FC-only training, heterogeneous *w_i_* and *I_i_* with homogeneous *σ*, as in (*53*), consistently yielded a decrease in *w_i_* along cortical hierarchy (Supplementary Fig. S17D–F). Conversely, the inverse strategy with only heterogeneous *σ_i_* consistently yielded a positive correlation between *σ_i_* and cortical hierarchy (Supplementary Fig. S17G–I). Together with the FC-only and hybrid-loss comparisons, these control analyses indicate that EC constraints shape the hierarchical organization of *w_i_* and *I_i_*, whereas heterogeneity in *σ_i_* is vital for matching FC.

### Link between node strength and regional heterogeneity of model parameters through local dynamics

The preceding analyses showed that reproducing the opposing empirical EC and FC NS patterns required regional heterogeneity within the tested model framework, and that multimodal training yielded a consistent hierarchical organization of regional parameters. Comparisons across training constraints and parameterizations further suggested a link between EC constraints and the hierarchy of recurrent strength and background input, and highlighted the contribution of noise heterogeneity to FC fitting. We therefore directly clarify these relationships using analytical solutions of linearization of MFM (Supplementary Note 5).

Following previous theoretical derivation (*26,54*), we linearized the MFM around a stable equilibrium ***S***^∗^ (Supplementary Note 5.1). For the Jacobian *J* of the linearized system, the analytical solution of integrated pulse response is EC = −*J*^−1^, while the stationary neural covariance satisfies 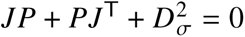, where *D_σ_* = diag(*σ*_1_,…, *σ_N_*) contains the intrinsic-noise amplitudes. Pearson normalization gives 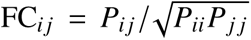 (Supplementary Note 5.2). We next asked whether the models retained after training operated in a regime where linearization around a stable equilibrium could explain major connectome patterns generated by the nonlinear MFM. In the optimallytrained model, analytical and simulated EC were significantly correlated (*r* = 0.897, *p* < 0.001; Fig. 7A). Analytical FC also significantly correlated with simulated FC, whether calculated from BOLD signals with Balloon–Windkessel hemodynamics (*r* = 0.807, *p* < 0.001) or directly from synaptic-gating activity (*r* = 0.858, *p* < 0.001; Fig. 7B). Across the 26 retained models, the mean similarities between analytical and simulated connectomes were 0.594±0.148 for EC, 0.762±0.102 for BOLD FC and 0.702 ± 0.080 for neural FC (Fig. 7C). Noise-free trajectories initialized at low and high activity approached the same region-specific steady states in most cases, although some models retained initial-state dependence (Supplementary Fig. S18). These results support the use of linearization to approximate the dynamics of the trained MFMs, providing an analytical basis for investigating how model parameters shape EC and FC.

**Fig. 7:**
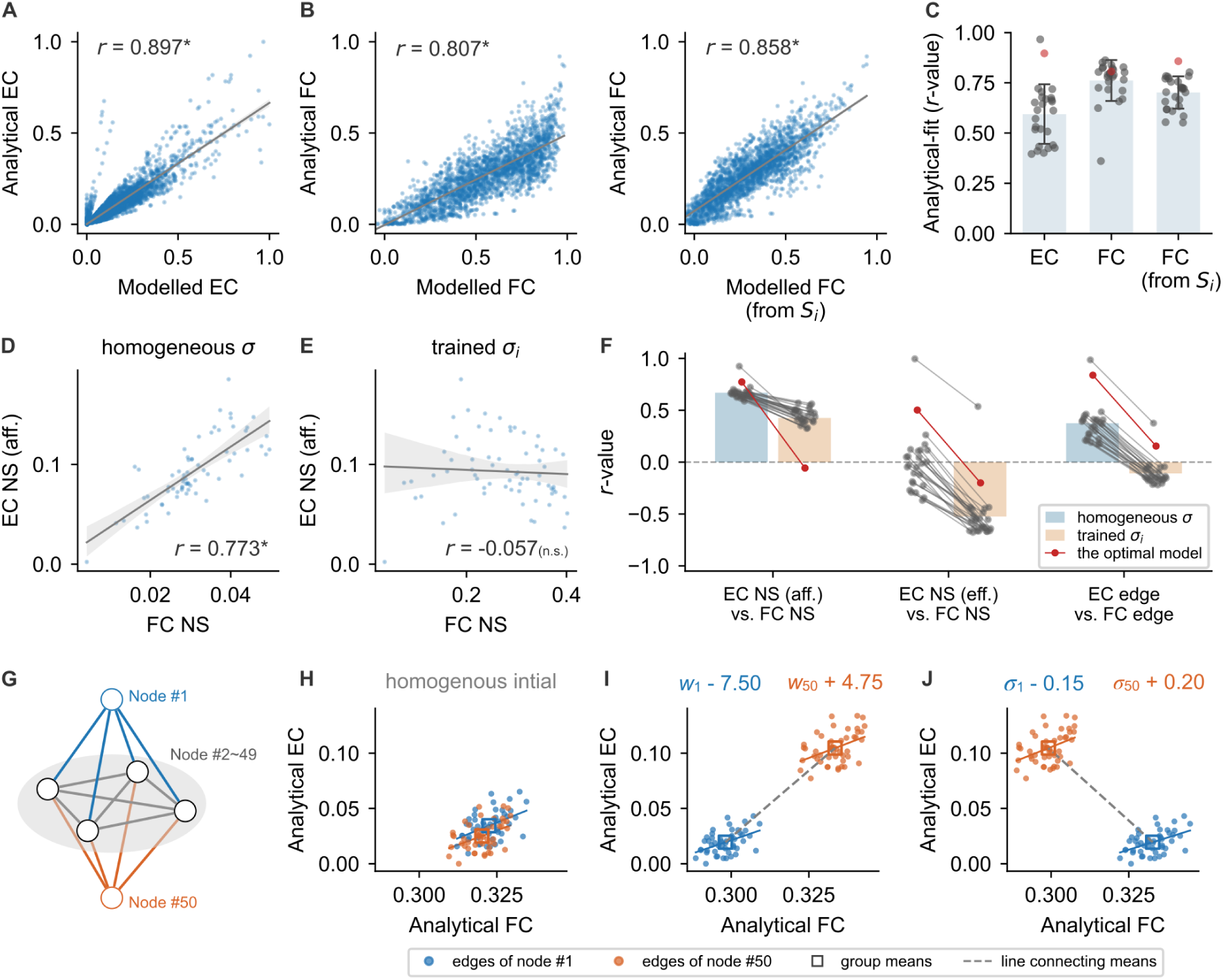
Analytical solutions reveal the roles of recurrence and intrinsic noise in shaping EC– FC node strength divergence. **A–B** Agreement between linearization-based analytical solutions and forward simulations in the optimally trained MFM. **A**, analytical EC versus simulated EC. **B**, analytical FC versus simulated FC calculated from BOLD signals (left) or directly from synapticgating activity *S_i_* (right). **C** Agreement between analytical solutions and simulations, quantified by Pearson’s *r*, across the 26 retained models for EC, BOLD FC and neural FC. Error bars show the mean and standard deviation; dots denote individual models, with the optimally trained model highlighted in red. **D–E** Relationship between analytical afferent EC and FC node strengths in the optimally trained model under homogeneous noise (**D**) and trained heterogeneous noise (**E**). In **A– F**, solid gray lines show linear fits; ∗ denotes *p* < 0.001, and n.s. denotes a nonsignificant correlation (*p* ≥ 0.05). Both simulated and analytical EC provide complete connectivity matrices without the sampling issue present in empirical measurements. Edge-wise comparisons involving EC thus use all off-diagonal entries, whereas FC–FC comparisons use upper-triangular entries. Node strengths are calculated as off-diagonal row means for afferent EC and FC, and column means for efferent EC. **F** Paired comparison across the 26 retained models of Pearson correlations between afferent EC and FC node strengths (left), efferent EC and FC node strengths (middle), and EC and FC connection weights (right). Blue and orange bars show means under homogeneous and trained heterogeneous noise, respectively. Connected dots represent the same model in the two conditions, with the optimally trained model highlighted in red. All three correlations are significantly suppressed under trained heterogeneous noise (all *p* < 0.001, paired *t*-tests). **G–J** Two-step analytical illustration of how recurrence and intrinsic noise reshape the relationships between EC and FC in a 50-node toy model. **G** Schematic of the symmetric structural network. Nodes 1 (blue) and 50 (orange) are each connected to a 48-node core subnetwork (nodes 2–49; gray). Connection weights are independently sampled from a uniform distribution over [0.75, 1.25] for edges within the core and those connecting each of the two nodes to the core. Nodes 1 and 50 have no direct structural connection; only representative core nodes and connections are drawn. To represent nodes 50 and 1 as higherand lower-order hierarchical nodes, respectively, we introduce parameter heterogeneity in two steps starting from a homogeneous initialization, illustrating how opposing EC and FC node-strength patterns can emerge along the hierarchy (**H–J**). **H** Analytical EC versus FC under homogeneous initialization (*w_i_* = −31.80, *I_i_* = 0.40, *σ_i_* = 1.00, *G* = 0.66). **I** Step 1: corresponding relationships after decreasing *w*_1_ by 7.50 and increasing *w*_50_ by 4.75. Compensating background input changes increase *I*_1_ by 1.306 and reduce *I*_50_ by 0.796, preserving each node’s equilibrium. Noise remains homogeneous. Node 50 now has higher mean incoming EC (afferent node strength) than node 1, while the two nodes’ mean EC and FC remain positively associated. **J** Step 2: analytical EC versus FC after decreasing *σ*_1_ by 0.15 and increasing *σ*_50_ by 0.20, while holding recurrent strengths and background inputs fixed after step 1. The ordering of mean EC remains unchanged, whereas that of mean FC is reversed, yielding opposing EC and FC means despite positive within-node EC–FC relationships. In **G-J**, blue and orange represent the 48 incoming connections from the core to nodes 1 and 50, respectively; colored solid lines show within-group linear fits, open squares indicate group means, and gray dashed lines connect the two means.

The analytical solutions further clarify how intrinsic noise shapes the relationship between EC and FC. Under homogeneous noise, *P* ∝ EC(**I** − Ξ), with the common covariance scale omitted because it cancels in Pearson normalization. Here, **I** is the identity matrix and Ξ accounts for asymmetry in *J*; for symmetric *J*, Ξ vanishes and *P* ∝ EC (Supplementary Note 5.5). Our theoretical arguments suggest that when the asymmetry correction Ξ has a limited effect on covariance (||Ξ||_2_ = 0.374 ± 0.111 across the 26 retained models), covariance largely follows EC, leaving FC strongly constrained by EC under homogeneous noise. With positive heterogeneous noise amplitudes, 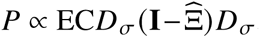, where *D_σ_* = diag(*σ*_1_,…, *σ_N_*) and 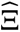 is the corresponding asymmetry correction in coordinates normalized by these amplitudes (Supplementary Note 5.6). Regional noise amplitudes thus provide additional degrees of freedom to reshape covariance and FC while leaving EC unchanged. This contrast motivated us to investigate whether regional noise heterogeneity accounts for EC–FC divergence in the trained models. To isolate its contribution, we compared analytically derived EC and FC under homogeneous and trained heterogeneous noise conditions while holding the deterministic dynamics fixed. In the optimally trained model, replacing the trained heterogeneous noise amplitudes with a common amplitude yielded a strong positive correlation between afferent EC and FC node strengths (*r* = 0.773, *p* < 0.001; Fig. 7D). Restoring the trained heterogeneous noise amplitudes markedly abolished this positive correlation (*r* = −0.057, n.s.; Fig. 7E). Note that the linearized model did not fully reproduce the negative EC–FC correlation, suggesting a contribution from nonlinear dynamics. Across the 26 retained models, correlations between FC node strength and both afferent and efferent EC node strengths, as well as edge-level EC–FC correlations, were significantly lower under trained heterogeneous noise than under homogeneous noise (all *p* < 0.001, paired *t*-tests; Fig. 7F). These comparisons show that trained noise heterogeneity suppresses EC-FC node strength coupling under homogeneous noise, thereby promoting EC–FC divergence.

We also examined how recurrence and background input shape EC. A path expansion of −*J*^−1^ separates structural propagation from local amplification: with homogeneous recurrence and a homogeneous equilibrium, afferent EC node strength increases with SC node strength at first order in structural coupling. Regional differences in *w_i_* reweight responses along structural paths, allowing EC node-strength patterns to depart from those of SC, where increasing *w_i_* reduces the net local decay and enhances EC node strength (Supplementary Note 5.3). Adjusting background input by 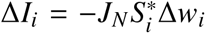 balances greater recurrence with lower background input, allowing regional EC to change without affecting equilibrium (Supplementary Note 5.4). Finally, we used a 50-node toy model with analytical solutions to illustrate the full sequence linking recurrence with background input compensation, and intrinsic noise to EC–FC divergence (Fig. 7G–J). We constructed a symmetric 50-node network with uniformly sampled connection weights, comprising a 48-node core subnetwork and two additional nodes (nodes 1 and 50) designated as lowerand higher-order hierarchical nodes, respectively, through subsequent parameter tuning (Fig. 7G). Starting from homogeneous regional parameters (Fig. 7H), we increased recurrence at node 50 and decreased it at node 1, while adjusting background inputs to preserve each node’s equilibrium. These changes produced higher mean incoming EC at node 50 than at node 1 (Fig. 7I). FC changed in the same direction: node 50 now had both higher EC and higher FC than node 1, so the line connecting the two group means retained a positive slope in the EC–FC plane (Fig. 7I). Introducing noise heterogeneity at fixed recurrent strengths and background inputs altered FC while leaving EC unchanged. Increasing *σ*_50_ reduced node 50’s mean FC, whereas decreasing *σ*_1_ increased node 1’s mean FC. Node 50 therefore has higher EC with lower FC, while node 1 has lower EC with higher FC (Fig. 7J). Incoming connections retained positive EC–FC relationships within each node, but the line connecting the two group means acquired a negative slope. Together, heterogeneity in recurrence and noise reproduced the paradoxical pattern of positive within-node EC–FC relationships coexisting with opposing EC and FC node-strength patterns on the same structural network, as suggested by our earlier empirical findings (Fig. 1, Fig. 2 and Supplementary Fig. S1).

We next examined whether the analytical findings hold in simulations of the original nonlinear MFM, by progressively introducing heterogeneity of regional parameters into homogeneous models to assess its impact on modelled NS: the retained homogeneous models were selected as the initial seeds; a vector 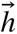 (normalized to zero mean and unit standard deviation) was generated to serve as a synthetic cortical gradient, consisting of 68 elements each sampled from Gaussian distribution; the synthetic cortical gradient was introduced into the homogeneous parameters in a linear form, with a scaling factor applied over 10 incremental steps (Fig. 8A). Note that the scaling factor defined the direction of parameter perturbation, being positive, negative and positive for 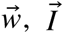 and 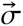, respectively, resulting in an increase of *w_i_* and *σ_i_* but decrease of *I_i_* in high-hierarchy nodes (Fig. 8A middle panel). At each step, the majority of modelled EC-NS exhibited a gradual increase in positive correlation with the introduced synthetic cortical gradient (Fig. 8B top and middle panels), accompanied by an inverse trend for FC-NS (Fig. 8B bottom panel). This phenomenon was consistent across 20 trials of random vectors for synthetic cortical gradient (Supplementary Fig. S19). Taken together, these results provide simulation-based evidence for the sufficiency of regional heterogeneity in tuning the NS patterns of modelled connectomes, through a process in which local recurrence increases the mean response or impact of the node, while local intrinsic noise suppresses FC.

**Fig. 8:**
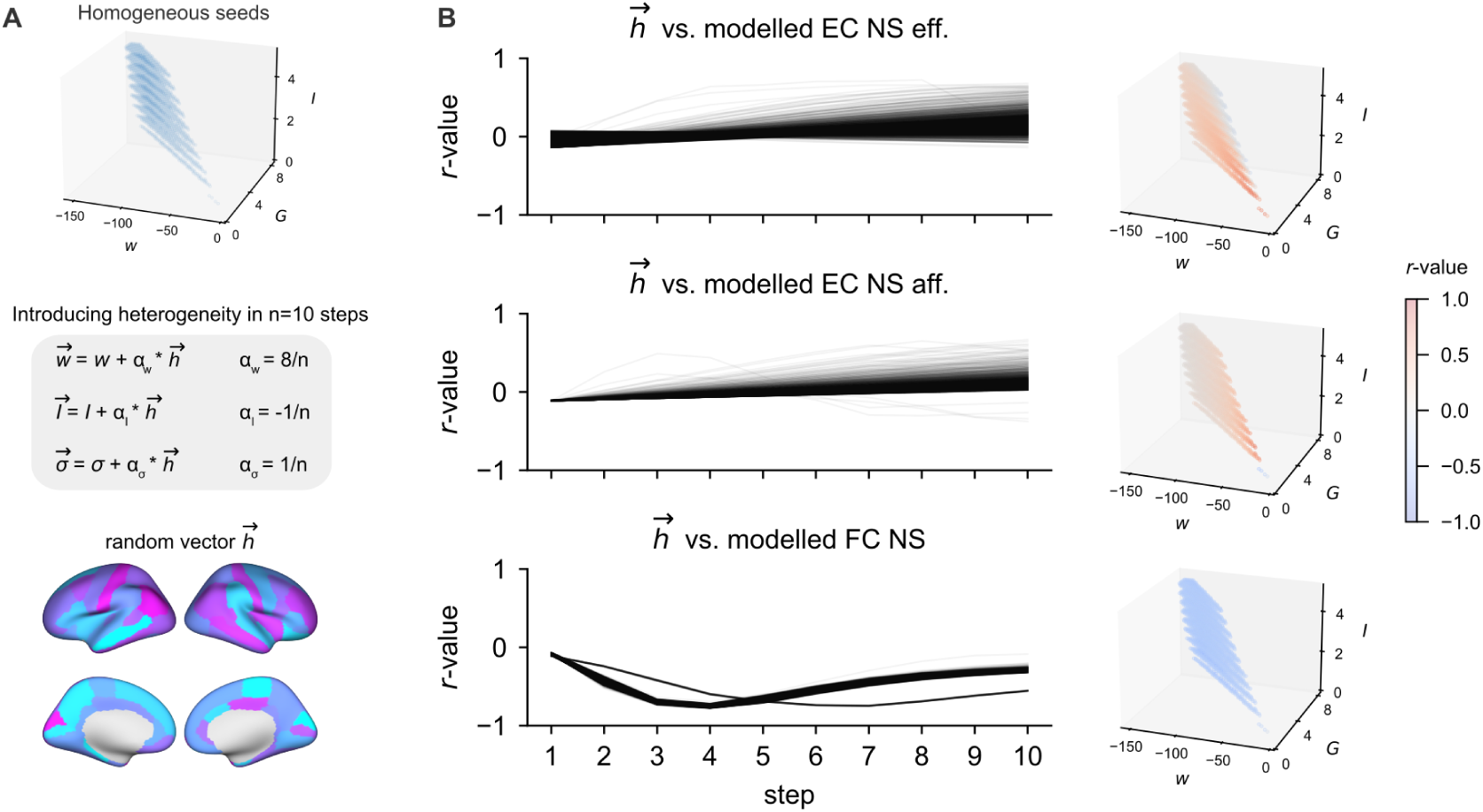
Regional heterogeneity tunes EC and FC through local dynamics. **A** Introduction of heterogeneity into the retained homogeneous 68-node models. A random vector 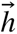, normalized to zero mean and unit standard deviation, is added incrementally over *n* = 10 steps, with per-step coefficients 8/*n*, −1/*n* and 1/*n* for 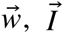 and 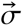, respectively. The initial parameter regime, update scheme and cortical map of 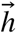 are shown from top to bottom. **B** Pearson correlations of 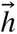 with modelled efferent EC NS (top), afferent EC NS (middle) and FC NS (bottom). Left: trajectories over the ten steps for individual model seeds. Right: final correlations, colour-coded in the parameter space of the initial homogeneous models.

## Discussion

Stimulation-derived EC and resting-state FC reveal opposing node-strength patterns along the cortical hierarchy, despite their shared relationship with the underlying structural network. A differentiable whole-cortex model constrained by both modalities accounts for this divergence, with hierarchical organization emerging from independently optimized regional parameters without a prescribed spatial map. Theoretical arguments based on MFM linearization, together with parameter perturbation experiments, link this organization to the complementary effects of local recurrence and intrinsic noise on evoked responsiveness and spontaneous correlations. Together, these findings connect regional connectome organization to local circuit dynamics and highlight the value of multimodal constraints for understanding the dynamical basis of cortical specialization.

### Opposing EC and FC node-strength patterns along the cortical hierarchy

FC node strength is significantly correlated with the FC principal gradient (Supplementary Fig. S20), a dominant axis of FC variation and an established marker of cortical hierarchy (*55*). Previous study has reported a negative correlation between FC node strength and cortical hierarchy, whereby higher-order association regions exhibit weaker overall FC than lower-order sensory areas (*54*). Interestingly, the trend was found to be refined during brain development, as FC node strength increased in sensory-motor regions and declined in association regions from childhood through adolescence (*45*). Here, we complemented this result of FC by our findings on EC, that both afferent and efferent EC node strengths exhibited positive correlation with the cortical hierarchy, i.e., greater responsiveness and influence of higher-order regions in stimulus–response paradigms. This hierarchy-related increase in EC node strength receives additional support from intracranial stimulation studies using different response measures: Momi et al. (*23*) reported stronger global evoked responses when stimulating higher-order networks, particularly in late response components; Terlau et al. (*56*) found larger late CCEP N2-amplitudes and a higher proportion of reciprocal connections toward association cortex. Our systematic comparison of EC and FC connectomes brings together these lines of evidence and directly reveals this opposing pattern along the cortical hierarchy, highlighting that FC alone does not fully characterize cortical dynamics.

### Bridging connectomic metrics and cortical dynamics

Graph-theoretical metrics have been widely used to characterize brain connectomes (*41–43,45–48*). However, bridging these descriptive measures with the underlying neural dynamics remains a fundamental challenge in the field (*2, 57–59*). Previous studies have begun to address this challenge through network diffusion dynamics (*2, 8*) and biophysical modelling approaches (*44, 54, 60*). Our work further advances this direction by establishing mechanistic links between connectomic metrics and cortical dynamics at both the global network and local node levels. At the global network level, the regime that facilitates signal propagation, rather than being artificially imposed (*52*), was first shown to emerge naturally from data-driven training, demonstrating its necessity to reproduce the empirical EC with identified indirect effects of multi-hop communication (*8, 9*). At the local node level, previous studies supported that a heterogeneous model significantly improves node strength fitting (*44, 54, 60*), with its modelled FC exhibiting increased similarity to the first eigenvector of Jacobian of linearized dynamics and decreased similarity to SC (*54*). Going beyond these observations, we developed a theoretical argument based on analytical solutions of the MFM to directly link model parameters to EC and FC node strengths (Supplementary Note 5). Validated through parameter perturbation analyses in both linearized toy model and the nonlinear MFM, this argument revealed distinct dynamical mechanisms underlying EC and FC node strengths (Figs. 7–8). Specifically, stronger local recurrent interactions enhanced regional responsiveness to external stimulation, thereby increasing EC node strength, whereas elevated intrinsic noise weakened spontaneous interregional correlations, leading to reduced FC node strength. These distinct parameter-dependent effects provide a mechanistic explanation for the opposing regional patterns of EC and FC node strength along the cortical hierarchy, highlighting how heterogeneous local circuit dynamics shape graph-theoretical measures of brain connectivity.

### Cortical hierarchy: functional constraints and biological interpretation of regional heterogeneity

Early mean-field models assumed homogeneous local circuit properties, meaning that regional parameters such as recurrent connection, background input and intrinsic noise are identical across all cortical regions (*26*). Subsequent modelling studies extended these models to a heterogeneous version by linearly introducing hierarchical gradients into regional parameters with significantly increased performance of model-empirical FC fit, thereby highlighting the sufficiency of cortical heterogeneity in shaping FC from SC (*44, 49, 54*). As an examination of necessity, Wang et al. (*53*) have allowed the parameters to vary across cortical regions using a stochastic optimization approach, but this resulted in an inverse gradient of recurrent connection strength that decreased rather than increased along the cortical hierarchy, in contrast to other modelling studies where increased recurrence also fits in the findings of increased timescale and synaptic excitation along hierarchy (*17, 49, 54, 61, 62*). This discrepancy calls for closer examination of whether a unique, hierarchy-specific cortical heterogeneity is necessary for cortical dynamics simulation (*17*).

Our results directly address this issue: across the 26 retained models, a consistent hierarchical pattern emerged, with positive, negative and positive correlations between cortical hierarchy and *w⃗*, *I⃗* and *σ⃗*, respectively. By taking the empirical findings on node strength together with the established relationship between graph-theoretical metrics and dynamics, we provided a analytical explanation for the hierarchical pattern that emerged in the parameters: an increase in *w_i_* combined with a decrease in *I_i_* leads to an overall enhancement of recurrent dynamics to promote EC node strength along the cortical hierarchy, whereas an increase in *σ_i_* suppresses FC node strength along the same axis. This mechanism reconciles the contradictory increase and decrease in recurrent strength along the cortical hierarchy previously reported by (*49*) and (*53*), respectively. It provides a mechanistic basis for the increasing recurrent strength reported by (*49*), which was originally imposed through a hierarchical prior. It also explains the decrease reported by (*53*) (also replicated under our training framework; Supplementary Fig. S17): under FC-only training, where *w⃗* and *I⃗* were trained heterogeneously but *σ* remained homogeneous, the suppression of FC node strength along the cortical hierarchy was primarily achieved through a decrease in recurrent dynamics. This increase in recurrent dynamics along cortical hierarchy is supported by the known structure of cortical excitation/inhibition microcircuits, wherein PV interneurons preferentially target the perisomatic region of excitatory pyramidal neurons, SST interneurons inhibit PV interneurons and target the dendrites of pyramidal neurons, and the SST-to-PV ratio, thought to regulate recurrent excitation, increases along the cortical hierarchy (*17, 63, 64*). Additionally, increased timescales in neural dynamics along the cortical hierarchy (*12, 19, 20*) have also been attributed to this increased recurrent excitation (*22, 61*). Taken together, it is suggested that higher-order regions serve as hubs for information integration and distribution through recurrent dynamics, in line with the ignition phenomenon and global neuronal workspace hypothesis of consciousness (*17, 65*). Notably, the increase in intrinsic noise along cortical hierarchy is consistent with experimental evidence showing increased resting-state fMRI signal variability along the same hierarchical axis (*64*). In particular, a recent study found that neural variability in iEEG signals, both following electrical stimulation and during cognitive tasks, is hierarchically structured, such that reduced variability in sensory cortex enhances sensory representation, whereas increased variability in association cortex supports working memory (*56*). Beside attributing this hierarchy of neural variability to increased reciprocal connectivity in higher-order cortex (*56*), we instead suggest a more straightforward interpretation, that it may be explained by intrinsic noise as a local property of the cortical nodes themselves. Considering that noise plays an important role in computation (*66*), e.g., neural sampling (*67*), it is speculated that higher-order regions not only integrate and maintain information through increased recurrence, but also participate in a hierarchical generative process with elevated neural variability (*68, 69*).

### Methodological implications

Training large-scale model in high-dimensional parameter space is a cutting-edge problem fraught with both software and hardware challenges (*37, 70*). We advanced the training and simulation of whole-cortex model with multi-fold contributions. First, we proposed a novel paradigm for studying the perturbational properties of the model by introducing EC, further emphasizing the indispensable role of multimodal approaches in studying brain connectomes (*1, 71, 72*). Second, together with emerging studies (*36, 38, 73*), our work demonstrates how deep learning can expand the scope of whole-cortex model training. Previous training strategies have mainly employed statistical inference (*54*), expectation-maximization (*53*) or evolutionary algorithm (*49*), all of which rely on extensive simulation-based sampling and are vulnerable to the curse of dimensionality as number of trainable parameters increases. As such, a widely used technique is to reduce the degrees of freedom of regional parameters by constraining them as a linear combination of preselected gradients (*49, 54*), which enables a feasible training process while substantially narrowing the explored solution space. By using gradients to guide parameter updates, the framework makes it feasible to optimize parameters without imposing such low-dimensional spatial constraints, allowing full-degree-of-freedom training of regional model parameters. The framework also offers both scalability and flexibility, with great potential to extend to larger-scaled parallel simulation and complex architectures, and interoperability from JAX to PyTorch across computational platforms (*35,36,38*). An important future direction is to extend the framework by augmenting connectome-based training with training based on resting-state (*74*), task-related (*75*) or stimulus-evoked time series (*23, 76*), with potential implications for advancing diagnosis and treatment of brain disorders (*77, 78*). Finally, we verified the utility of training techniques such as gradient clipping and parameter-specific learning-rate multipliers for training the MFM. Interestingly, we observed that the training trajectories converge to a critical regime, suggesting that criticality, beyond being vital for fitting performance, is also compatible with BPTT and may facilitate the training process. Given that criticality is a key property of large-scale brain dynamics (*79*), it is worthwhile to further develop training techniques that leverage criticality (*80*), alongside well-established methods (e.g., truncated BPTT (*81*) and e-prop (*82*)), for training large-scale models under diverse optimization objectives and resource-limited conditions, such as future individualized modelling and clinical applications.

### Limitations

Several limitations of this study should be discussed. First, electrical stimulus-response experiments can only be conducted on epilepsy patients with implanted iEEG. Although F-TRACT project has applied conservative procedures to exclude pathological conditions (*5–7*), the gap between its epilepsy patients and healthy subjects from HCP remains an open issue. This gap could be narrowed through future integration of functional and diffusion MRI into F-TRACT protocol or through initiative of the Epilepsy Connectome Project (*83*). Second, due to the practical challenges of neurostimulation experiments, only a single population-level EC was available, as it was derived by averaging across a large number of subjects and trials (*5–7*). Although model generalizability can be examined following a cross validation scheme by randomly splitting the connectome into train and test sets (*8*), this group-averaging nature in EC limited our ability to perform external validation across individuals and medical sites, which calls for larger sample size and better utilization of multi-center potential of F-TRACT. Third, we opted to use the MFM with a coarse parcellation, considering computational cost, the sampling issue in EC, and the need to enable direct comparison with previous studies (*49, 53*). Future work should move toward higher-resolution parcellation such as HCP-MMP with 360 parcels (*50*), and the model incorporating nonlinear dynamics of inhibitory populations (*54, 84*). Fourth, although our linearization-based theoretical analysis provides an explanation for our findings, the proposed suppressive effect of intrinsic noise on FC node strength is not universal but depends on the underlying parameter regime. Whether a more general principle governs how increased intrinsic neural noise constrains a node’s communication capacity to the rest of the network remains an open theoretical question. Fifth, the Balloon-Windkessel model for fMRI signal simulation substantially increases the number of simulation time steps as well as the difficulty of BPTT training. We bypassed this issue by excluding the Balloon–Windkessel model during training but conducting it during testing simulations, with validation of limited impact on FC simulation (Supplementary Fig. S21). To ensure the feasibility of training directly based on fMRI signals, future techniques are required to either simplify the hemodynamics or replace with temporal kernel convolution (*85*). Lastly, only static information from brain connectomes was involved in the current pipeline, though there is potential to incorporate dynamic properties, such as functional connectivity dynamics (FCD) (*49*), or to directly utilize time series through next-step prediction (*74*) in future investigations.

## Materials and methods

### Datasets

We used the preprocessed and group-averaged functional connectome (FC) and structural connectome (SC) matrices openly available in ENIGMA toolbox (*40*), based on a cohort of 207 unrelated healthy subjects from the Human Connectome Project (HCP) (*39*). Detailed preprocessing procedures are described in (*40*) and are available through the ENIGMA Toolbox documentation (http://enigma-toolbox.readthedocs.io). Notably, subject-specific FC matrices were thresholded by setting negative connections to zero, and then z-transformed and averaged across subjects. The averaged matrix was tanh-transformed back to Pearson’s r-values (FC_ij_ ∈ [−1, 1]). SC was defined as the log-transformed number of reconstructed streamlines between pairs of cortical regions inferred by diffusion MRI, where group averaging was performed by a distance-dependent thresholding procedure accounting for the edge length distribution of individual subjects (*86*). Max-scaling, i.e., dividing by the maximum value, was performed as to SC (SC_ij_ ∈ [0, 1]).

The response probability matrix from the Functional Tractography Project (F-TRACT) (*5–7*) was obtained as the effective connectome (EC; see Supplementary Note 3 for this terminology). We adopt the version F-TRACT_P_10_v2210 (*87*) with following specifications: 583 epileptic patients after applying stringent criteria to exclude data with epileptic-like interictal spikes, and poststimulus period up to 200 ms to account for potential effects of indirect signal propagation. Data are openly accessible via the F-TRACT database (https://f-tract.eu/), with methodological details described in (*5–8*). In brief, F-TRACT is an ongoing project dedicated to measure, collect and aggregate cortico-cortical evoked potentials (CCEPs) from patients with drug-resistant epilepsy. These patients were implanted with intracranial electroencephalography (iEEG) electrodes as part of their clinical evaluation for candidacy of resective surgery. Time series of CCEPs were measured by single-pulse electrical stimulation (inter pulse interval is equal to 1 s) of a single stimulation site (two contiguous electrode contacts) and simultaneous recording with all remaining electrode contacts. Time series were preprocessed to filter bad channels, correct stimulation artifact, re-reference with bipolar montage (i.e., adjacent electrode contact as reference) and band-pass filter between 1 and 45 Hz. Time series were then z-scored based on the pre-stimulus baseline and thresholded as significant response if z > 5 during post-stimulus period. Each electrode contact was localized based on individual patient’s T1 MRI and CT scans, and was assigned to a specific regional label of an anatomical parcellation by taking the most frequent voxel label in a sphere of 3 mm radius around the contact center. A brain-wide, group-level response probability matrix (denoted EC) was constructed based on all stimulus-response pairs of CCEPs trials across patients. Entry in the matrix (EC_ij_ ∈ [0, 1]) indicates the probability of observing significant response in region *i* (row index) following stimulation of region *j* (column index), thereby quantifying the strength of effectiveness from region *j* to *i* (also denoted as EC_j→i_). This naturally results in an asymmetric EC matrix, in contrast to the symmetric SC and FC matrices. The original matrix, with row indexing stimulus and column indexing response, was transposed in this study to conform to the matrix multiplication convention (*v⃗*_response_ = EC *v⃗*_stimulus_) for column vectors (*v⃗*) of input stimulus and output response of cortical regions. Note that a minimum sample size of stimulus-response pairs (N ≥ 10) was required. If this condition was not met, the entry was masked with an NaN (Not a Number) value and excluded from any subsequent computations (Supplementary Fig. S10). Diagonal entries were additionally masked as NaN values, as we only focused on the information of inter-regional signal propagation.

In line with previous modelling studies (*49, 53*), and taking into account the incomplete measurement issue in EC (Supplementary Fig. S10), the Desikan-Killiany parcellation (*51*) with 68 cortical regions (Supplementary Table S1) was used for all connectomes. The cortical gradients were from (*49*), including T1w/T2w ratio for myelin and gene expression of parvalbumin (PV) and somatostatin (SST).

### Node metrics

We used node strength (NS), a standard graph-theoretic metric also referred to as weighted degree (*41*) or functional connectivity strength (*45,46*), to assemble the results of edges (connections) into nodes (regions) level properties. Following previous studies (*42,44,48*), we adopt a normalized form calculated as the mean weight of each node’s available connections. The metrics were defined as follows:

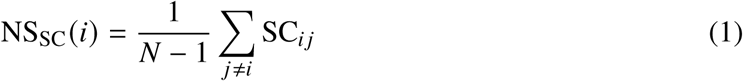

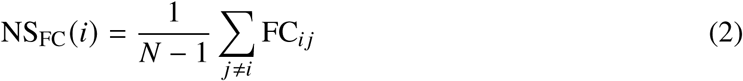

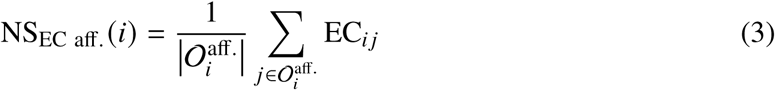

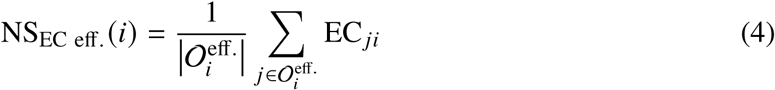

where NS denotes the mean weight of edges to/from node *i*, and *N* is the number of cortical regions. The sets 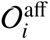. and 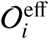 contain, respectively, source regions *j* ≠ *i* with observed EC*_i j_* and receiver regions *j* ≠ *i* with observed EC*_ji_*. Vertical bars denote set cardinality, and the EC means are defined for nonempty sets. Thus, self-connections and unobserved (NaN) EC entries are excluded (Supplementary Fig. S10). As EC is asymmetric (directed), we calculated the afferent (aff.) and efferent (eff.) NS for in-degree and out-degree connections, respectively. The afferent NS is interpreted as mean response of the node across all stimulus conditions, whereas the efferent as the mean impact.

### Whole-cortex dynamic mean-field model

We adopt the whole-cortex MFM, a mean-field reduction of a leaky integrate-and-fire spiking neural population, derived by simplifying fast glutamatergic (AMPA) and GABAergic activity and tracking the slow glutamatergic (NMDA) activity (*26*). The dynamics of each cortical region *i* (*i* = 1, 2,…, 68) are described by the following nonlinear stochastic differential equations:

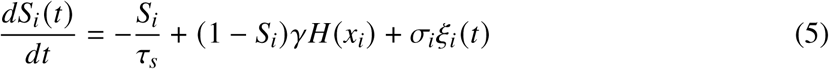

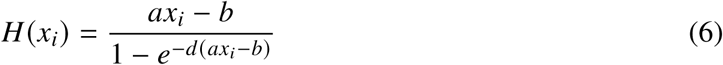

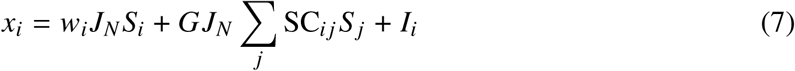

where *S_i_* denotes the synaptic gating variable (i.e., fraction of open channels for NMDA receptor) of cortical region *i*. The term *σ_i_ξ_i_* (*t*) represents Gaussian white noise scaled by *σ_i_* during restingstate simulations, or a prescribed external input during evoked-response simulations. The total input current *x_i_* encompasses three factors: (1) intra-regional input tuned by recurrent connection weight *w_i_*; (2) inter-regional input controlled by the SC matrix (SC*_i j_* is the SC between regions *i* and *j*) and global coupling factor *G*; (3) background input current *I_i_* including background activity and inputs from subcortical relays. *H*(*x_i_*) is the Abbott-Chance activation function (*88*), a softReLU shaped input-output nonlinearity that converts total input current into population firing rate. Note that a piecewise surrogate was applied to this activation function to address its numerical issue during automatic differentiation (see Supplementary Fig. S22 and Supplementary Note 4 for details). Following previous studies (*26, 49*), parameters for input–output function were set to *a* = 270 nC^−1^, *b* = 108 Hz and *d* = 0.154 s. Synaptic coupling *J_N_* was set to 0.2609 nA. Synaptic kinetics were set to *γ* = 0.641 and *τ_s_* = 0.1 s. Model simulation was numerically applied in a forward EulerMaruyama form:

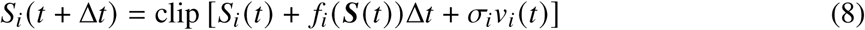

where 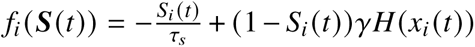 denotes the deterministic component of the dynamics. A clipping function clip(x) = min{1, max{0, x}} was applied to ensure synaptic gating variable within its valid range (*S_i_* ∈ [0, 1]), particularly under perturbation of pulsed stimulus. Time step was set to Δ*t* = 0.5 ms. Note that previous studies typically used coarser time steps of 10 or 100 ms (*26, 49, 53*), primarily to model slow fluctuation for fMRI signal while reducing computational cost. Here, a more precise time step was required for simulation of CCEPs, with a finer temporal scale and sampling rates up to 2 kHz (*89*). In resting-state simulation, *v_i_* (*t*) for region *i* was independently sampled random noise from standard Gaussian scaled by 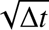, with its amplitude tuned by *σ_i_*. BOLD signals were derived from synaptic gating variables using the Balloon-Windkessel hemodynamic model (*85, 90*), with parameters set according to previous studies (*49, 53*). As EC is an aggregated connectome with diminished noise, we simulated single-pulse electrical stimulus for CCEPs in a noise free fashion, where *v_i_* (*t*) is the injected perturbation signal with *σ_i_* set to 1 for the targeted region and 0 for all remaining regions.

### Model training

The MFM was implemented as an RNN using BrainPy, a JAX and XLA embedded platform that enables parallel simulation and GPU-accelerated optimization of dynamic system (*35*). EC fitting and FC fitting were designed in an end-to-end fashion as follows.

### EC fitting

Virtual stimulus trial was performed by delivering the input time series of single-pulse stimulus to a target region and recording the response time series of neural activity (*S_i_*) of remaining regions. Each trial was run for 0.5 s, spanning a window of [-250, 250] ms with pulse timing set at 0 ms and pulse amplitude set at 1 (i.e., *S _j_* (*t* = 0) → 1, where *j* denotes the targeted region). 1.5 s intertrial intervals were run before and after each trial, spanning the periods of [–1000, –250] ms and [250, 1000] ms, and resulting in a total simulation duration of 2s. Note that inter-trial intervals were to enable the model to re-stabilize to its equilibrium state, and not involved in the following loss calculation. In line with previous studies (*6, 7, 89*) which focused on the [-200, 200] ms window for CCEPs while discarded the [-10, 10] ms interval for artifact removal, we set the pre-stimulus baseline of [-200, −10] ms and post-stimulus period of [10, 200] ms. A time mask was then defined by assigning −1, 1 and 0 to pre-stimulus, post-stimulus and all other time points, respectively. The response time series of region *i* after stimulating region *j* was dot-multiplied with the time mask and then summed along time axis, yielding the baseline corrected area under the stimulus evoked curve, which served as a modelled measurement of strength of effectiveness (i.e., modelled EC_ij_). Batch-wise training was adopted with a batch size equal to the number of cortical regions (N = 68), so that a batch comprised 68 trials with each trial stimulating a unique region in a randomly shuffled order. As a result, every training iteration produced a complete modelled EC matrix. Subsequently, a correct scale of modelled EC_ij_ ∈ [0, 1] was ensured using a min-max scaling function, defined as min-max(*x*) = (*x* − *x_min_*)/(*x_max_* − *x_min_*), where *x_min_* and *x_max_* represent the minimum and maximum values of the input *x*, respectively. The mean squared error (MSE) loss between modelled and empirical EC was computed with masked NaN entries of empirical EC. As all procedures were implemented with tensor operators enabling automatic differentiation (autodiff), the MSE loss was minimized using a deep learning optimization scheme back-propagation through time (BPTT) (*33*), which unfolds the RNN across time into a deep forward computational graph for error back-propagation. By leveraging computational advantages of deep learning, in addition to the parameter *G*, we were able to relax intra-regional parameters to be independently trainable across regions, with each region *i* having its own trainable *w_i_* and *I_i_*. Gradients were clipped to an absolute threshold of 10^10^ to prevent instability of exploding gradients during BPTT training. Adam optimizer was adopted with default hyperparameters first moment decay *β*_1_ = 0.9 and second moment decay *β*_2_ = 0.999 (*91*).

### FC fitting

Virtual resting-state simulation was run for 2 s (same as in EC fitting) without applying BalloonWindkessel model. Model FC was obtained by computing Pearson’s correlation between time series of neural activity *S_i_* (*t*). The MSE loss between modelled and empirical FC with masked diagonal entries was minimized with BPTT. With embedded noise information in FC, the training enables optimization of all parameters *G*, *w_i_*, *I_i_* and *σ_i_*, with batch size of 68, gradient clipping at 10^5^ and Adam optimizer. Note that the choice of a short window was to mitigate the long-sequence BPTT training issues (*92*) (e.g., vanishing or exploding gradients, high computational cost), whereas modelling fMRI signals would require simulation sequences several orders of magnitude longer. During testing, after a 5 min warm-up period for signal stability, we adopt 10 min fMRI signals to generate FC with Balloon-Windkessel model applied. Supplementary Fig. S21 illustrates the validation for modelled FCs obtained with/without the Balloon-Windkessel model.

### Hyperparameter selection and cross-validation

To select hyperparameters and evaluate generalization to held-out connections, we performed fivefold cross-validation following (*8*) (Supplementary Fig. S9A). The candidate Adam learning rates were {0.01, 0.05, 0.1, 0.5, 1.0}. We also evaluated a parameter-specific learning-rate multiplier for recurrent strength *w_i_*, setting its learning rate to either one or ten times the base learning rate (lr*_w_* ×1 or lr*_w_* × 10, respectively), while retaining the base learning rate for the other trainable parameters. This approach follows the general practice in deep learning of assigning different learning rates to different subsets of model parameters (*93, 94*). EC entries (excluding NaN) were randomly partitioned into five folds. As FC is symmetric, its partition was applied only to the upper-triangular entries (excluding the diagonal) and then mirrored to the lower triangle to prevent data leakage between training and test sets. Each fold served as the test set in turn, while the remaining four folds were used for training. Predictions for held-out region pairs were concatenated across folds and compared with their empirical values using Pearson’s *r*. For computational efficiency, crossvalidation used initial recurrent strengths from −150 to 20 in steps of 5.

The highest pooled out-of-sample EC fit was obtained with a learning rate of 0.05 and lr*_w_* × 10 (*r* = 0.66; Supplementary Fig. S9B,F), whereas the highest pooled FC fit was obtained with a learning rate of 0.05 and lr*_w_* × 1 (*r* = 0.69; Supplementary Fig. S9C,I). These settings were used for the first stage of two-stage training and for FC-only training, respectively. Modelled connectomes and training trajectories were consistent across folds (Supplementary Fig. S9D–F,G–I).

### Training procedures

We trained the model in which intra-regional parameters were optimized independently across regions, such that the total degree of freedom of all trainable parameters (*G*, *w_i_*, *I_i_*, *σ_i_*) is 1 + 3 × 68 = 205. Note that, to span a sufficiently broad parameter space encompassing the signal propagation regime (similar to the GBA regime (*52*)), we decided to relax the range of *w_i_* and *I_i_* to include negative values, which were interpreted as representing net self-recurrent or background inhibition mediated by interneurons (see Supplementary Fig. S4 and Supplementary Note 1). Parameter *G* was still constrained to positive values as it exclusively represents excitatory inter-regional connections, and *σ_i_* was kept positive as it denotes an amplitude measure.

Model initialization was homogeneous across regions, with *w_i_* = *w*_init_, *I_i_* = *I*_init_ and *σ_i_* = *σ*_init_ for all *i*. Initial seeds were defined by *G*_init_ = 1, *I*_init_ = 0 and *σ*_init_ = 10^−5^, with *w*_init_ varied from −150 to 20 in unit increments. This strategy balanced broad coverage of parameter space against computational cost by varying the initial recurrent strength. EC fitting used the same initialization for *G*, *w_i_* and *I_i_*, but simulated evoked responses without intrinsic noise. Full-batch training (batch size = 68) was adopted for each epoch.

Two-stage EC–FC training first optimized *G*, *w_i_* and *I_i_* by EC fitting for 10,000 epochs with an Adam learning rate of 0.05 and a learning-rate multiplier of 10 for *w_i_* (lr*_w_* × 10). The subsequent FC-fitting stage optimized only *σ_i_*, keeping *G*, *w_i_* and *I_i_* fixed, for 1,000 epochs with a learning rate of 0.1 to limit computational cost. In comparison, single-stage FC-only training optimized *G*, *w_i_*, *I_i_* and *σ_i_* for 10,000 epochs with a learning rate of 0.05 and a learning-rate multiplier of 1 for *w_i_* (lr*_w_* × 1).

Single-stage hybrid EC–FC training jointly optimized *G*, *w_i_*, *I_i_* and *σ_i_* by minimizing the objective,

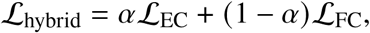

where ℒ_EC_ and ℒ_FC_ are the EC and FC MSE losses defined above, and the EC–FC loss ratio *α* ∈ {0.1, 0.25, 0.5, 0.75, 0.9} sets the EC and FC loss coefficients to *α* and 1 − *α*, respectively. For each *α*, training used Adam for 10,000 epochs with a learning rate of 0.05, a batch size of 68 and a learning-rate multiplier of 1 for *w_i_* (lr*_w_* × 1). Initialization and settings for all three training strategies are summarized in Supplementary Table S2. Training was performed on an A100 GPU (NVIDIA Inc.) with 80 GB memory and CUDA version 11.8.

### Grid search of regionally homogeneous models

To evaluate whether homogeneous regional parameters could account for the opposing EC and FC NS patterns, we held *w_i_* = *w*, *I_i_* = *I* and *σ_i_* = *σ* across all cortical regions and performed a four-dimensional grid search over *G*, *w*, *I* and *σ*. The sampled ranges were *G* ∈ {0.2 : 0.2 : 8}, *w* ∈ {−150 : 2 : 20}, *I* ∈ {0.5 : 0.5 : 5} and *σ* ∈ {0.5 : 0.5 : 5}, where the notation specifies the start, step size and end of each range. This yielded 40 × 86 × 10 × 10 = 344,000 parameter combinations. The sampled combinations are projected onto the (*w*, *G*, *I*) axes in Supplementary Fig. S12A (left).

Homogeneous models with Pearson model–empirical correlations of *r* > 0.4 for both EC and FC at the edge level were designated as retained homogeneous models (Supplementary Fig. S12A, right). Across all grid-searched homogeneous models, we quantified the model–empirical correlations for afferent EC-NS and FC-NS, using *r* = 0.5 as the threshold for each NS fit (Supplementary Fig. S12B). For the retained homogeneous models, we also computed the Pearson correlation between modelled afferent EC-NS and FC-NS to assess whether the empirical negative relationship was reproduced (Supplementary Fig. S12C).

### Analytical solutions of EC and FC

Given model parameters *G*, *w_i_*, and *I_i_*, we first obtained a stable deterministic equilibrium ***S***^∗^ by forward simulation of the noise-free MFM and evaluated the Jacobian *J* at this equilibrium. The integrated response to a unit pulse gives analytical EC as EC = −*J*^−1^. For resting-state fluctuations, we obtained the stationary covariance *P* by solving 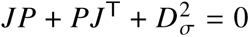, given the regional intrinsic-noise amplitudes *σ_i_* with *D_σ_* = diag(*σ*_1_,…, *σ_N_*). Analytical FC was computed by Pearson normalization, 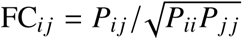. See Supplementary Note 5 for the full derivations.

### 50-node toy model

We constructed a 50-node toy model with a symmetric SC matrix, comprising a 48-node core subnetwork (nodes 2–49) and two additional nodes (1 and 50). Connection weights within the core and between each of the two additional nodes and the core were independently sampled from the same uniform distribution on [0.75, 1.25]. Self-connections and the direct connection between nodes 1 and 50 were absent.

We initialized the toy model with homogeneous parameters: *G* = 0.66, *w_i_* = −31.80, *I_i_* = 0.40, and *σ_i_* = 1.00 for all nodes *i*. We obtained the equilibrium ***S***^∗^ by integrating the deterministic dynamics for 5 s. This equilibrium, which could differ across nodes, was preserved during subsequent recurrence changes by adjusting background input according to 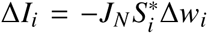. Parameter manipulations proceeded in two steps. First, we decreased *w*_1_ by 7.50 and increased *w*_50_ by 4.75 while keeping all noise amplitudes fixed at 1.00. The corresponding compensation increased *I*_1_ by 1.306 and reduced *I*_50_ by 0.796. Second, we held the resulting recurrent strengths and background inputs fixed and decreased *σ*_1_ by 0.15 while increasing *σ*_50_ by 0.20. Core-node parameters remained at their initial values throughout both steps.

At each parameter setting, we evaluated analytical EC and FC at the fixed equilibrium using the analytical solutions described above. All Jacobian eigenvalues were negative real values throughout both steps, confirming local stability of the equilibrium.

### Statistics

Spearman’s correlation *ρ*-value was applied for correlation analyses involving model parameters, considering the non-normality of distribution of model parameters resulting from the nonlinearity of model dynamics. Pearson’s correlation r-value was used for all other correlation analyses and for quantifying model fit. Bonferroni correction for multiple comparisons was applied to control the family-wise error rate (FWE-corrected).

## Acknowledgments

Data were provided in part by the Human Connectome Project, WU-Minn Consortium (Principal Investigators: David Van Essen and Kamil Ugurbil; 1U54MH091657) funded by the 16 NIH Institutes and Centers that support the NIH Blueprint for Neuroscience Research; and by the McDonnell Center for Systems Neuroscience at Washington University. We thank the ENIGMA Toolbox team for making the HCP-derived connectivity matrices available, and the F-TRACT Consortium for sharing their data through EBRAINS. Part of the computations in this study were run on the Siyuan-1 cluster supported by the Center for High Performance Computing at Shanghai Jiao Tong University.

## Author contributions

Conceptualization: X.C., S.L. and D.Z.; Methodology: X.C. and Y.F.; Data curation: X.C.; Investigation: X.C. and Y.F.; Formal analysis: X.C.; Visualization: X.C.; Validation: X.C.; Project administration: S.L. and D.Z.; Funding acquisition: S.L. and D.Z.; Supervision: S.L. and D.Z.; Writing - original draft: X.C.; Writing - review and editing: X.C., S.L. and D.Z.

## Data, code and materials availability

Only publicly available data were used in this study: functional and structural connectome matrices are openly available in ENIGMA toolbox (https://enigmatoolbox.readthedocs.io/); stimulation-based effective connectome matrix is openly available from F-TRACT database (https://f-tract.eu/). The processed data and code are publicly available in the GitHub repository (https://github.com/Dr-Chen-Xiaoyu/EC-FC-BPTT-DecoModel). No physical materials were generated in this study.

## Supplementary materials

Supplementary Notes 1 to 5

Figs. S1 to S22

Tables S1 to S3

References

