## Supplemental Information for "A dynamical mechanism for opposing cortical gradients in resting-state functional and stimulation-derived effective connectomes"

**This PDF file includes:**

Supplementary Notes 1 to 5

Figures S1 to S22

Tables S1 to S3

References

**Supplementary Tables** 17

**Supplementary Figures** 20

**References** 49

### Supplementary Notes

#### 1. Relaxing recurrent strength to negative domain

We treated the mean-field model (MFM) as a phenomenological model. Because the inhibitory population was linearized and simplified into the recurrent connection (*1*), we relaxed the range of recurrent strength $w$ to include negative values to represent net feedback inhibition. This relaxation allows the model to enter the Global Balanced Amplification (GBA) regime (*2*), thereby facilitating inter-regional signal propagation as demonstrated in a two-node example (Fig. S4A–B) and the homogeneous model (Fig. S4C).

A negative $w$ could be interpreted as an increased weight of I-to-E connection, as illustrated in a two-node example of the E-I MFM with both excitatory (E) and inhibitory (I) populations (*3, 4*) (Fig. S4D–E). To maintain comparability with previous studies, the present study retains the MFM, and leaves a full exploration of the E-I MFM for future studies, as outlined by the "Limitations" section in main text.

#### 2. The bifurcation line

We demonstrated that a bifurcation line, previously referred to as the "first bifurcation" in (*5*), constitutes a necessary condition for bistability in the homogeneous, noise-free MFM. It is claimed that bistability can arise only if

$$w + \lambda G \geq w_0\ (I_0) \tag{S1}$$

where $\lambda$ is the spectral radius of SC and $w_0$ is a constant derived based on $I_0$. The proof of the claim is provided below, starting with the following lemma.

**Lemma**: $\exists\, k \in \{1, \ldots, N\}$ where $N$ is the number of nodes, such that

$$\sum_{j=1}^{N} \mathrm{SC}_{kj} S_j \leq \lambda S_k. \tag{S2}$$

**Proof**: since SC is real and symmetric, SC can be eigen-decomposed into

$$\mathrm{SC} = Q \Lambda Q^{\mathrm{T}} \tag{S3}$$

where $\Lambda$ is the diagonal matrix of eigenvalues and $Q$ is the orthogonal matrix which is unitary.

Using sub-multiplicativity and unitary invariance of matrix 2-norm yield the following inequality:

$$\|\mathrm{SC}\vec{S}\|_2 = \|Q\Lambda Q^{\mathrm{T}}\vec{S}\|_2 = \|\Lambda Q^{\mathrm{T}}\vec{S}\|_2 \leq \|\Lambda\|_2\|Q^{\mathrm{T}}\vec{S}\|_2 = \lambda\|\vec{S}\|_2 \tag{S4}$$

where $\|\Lambda\|_2 = \lambda$ as 2-norm of a diagonal matrix is the largest absolute diagonal element, i.e., spectral radius.

Expanding 2-norm into element-wise form gives:

$$\sqrt{\sum_{k=1}^{N}\left(\sum_{j=1}^{N}\mathrm{SC}_{kj}S_j\right)^2} \leq \lambda\sqrt{\sum_{k=1}^{N}(S_k)^2}. \tag{S5}$$

By assuming the lemma is false, then for $\forall\, k \in \{1, \ldots, N\}$ there is

$$\sum_{j=1}^{N}\mathrm{SC}_{kj}S_j > \lambda S_k. \tag{S6}$$

As all elements of SC and $S$ are non-negative, we have:

$$\sqrt{\sum_{k=1}^{N}\left(\sum_{j=1}^{N}\mathrm{SC}_{kj}S_j\right)^2} > \sqrt{\sum_{k=1}^{N}(\lambda S_k)^2} = \lambda\sqrt{\sum_{k=1}^{N}(S_k)^2} \tag{S7}$$

which results in a contradiction and proof of the lemma.

Next, we consider the model with $I_0 = 0$ and $G = 0$ (i.e., single node condition). The fixed point is given by

$$f(S) = \frac{dS(t)}{dt} = -\frac{S}{\tau_S} + (1-S)\gamma H(wJ_N S) = 0 \tag{S8}$$

where a critical value can be found at $w = w_0 \approx 2.9185$. If $w \leq w_0$, there is only one stable point $S_{\mathrm{low}} \approx 10^{-6}$, where $f(S) \leq 0$ for $S > S_{\mathrm{low}}$. If $w > w_0$, there are three fixed points: two stable points and one saddle point (Fig. S6A–B). Note that $w_0$ is non-linearly derived based on $I_0 = 0$, but follows a similar argument in the case of non-zero $I_0$ (Fig. S6C).

For $G > 0$ (i.e., coupled nodes condition), the existence of the high stable state with solution away from zero requires

$$\frac{dS_i(t)}{dt} = -\frac{S_i}{\tau_S} + (1-S_i)\gamma H\left(wJ_N S_i + GJ_N\sum_{j=1}^{N}\mathrm{SC}_{ij}S_j\right) = 0 \tag{S9}$$

with $S_i > S_{\text{low}}$ for $\forall\, i \in \{1, \ldots, N\}$. Since $H$ is an increasing function, and by the lemma, there is $S_k$:

$$0 = -\frac{S_k}{\tau_S} + (1 - S_k)\gamma H\left(wJ_N S_k + GJ_N \sum_{j=1}^{N} \mathrm{SC}_{kj} S_j\right) \leq -\frac{S_k}{\tau_S} + (1 - S_k)\gamma H((w + \lambda G) J_N S_k). \quad \text{(S10)}$$

If $w + \lambda G < w_0$, under the statement of single node equation with condition of $S_k > S_{\text{low}}$, we have the right-hand side of the above equation negative:

$$-\frac{S_k}{\tau_S} + (1 - S_k)\gamma H((w + \lambda G) J_N S_k) < -\frac{S_k}{\tau_S} + (1 - S_k)\gamma H((w_0) J_N S_k) \leq 0 \quad \text{(S11)}$$

which results in a contradiction and proof of the claim.

For illustration, we plotted the bifurcation line $w + \lambda G = w_0$ with $w_0 = 2.9185$ and the empirical spectral radius $\lambda = 13.7563$, with confirmed validity through grid search of parameter space of the homogeneous, noise-free MFMs (Fig. S6D).

#### 3. Terminology about effective connectome

The connectome derived from stimulus-response paradigm has been variously named in previous literature, such as "effective connectome" (*6, 7*), "perturbome" (*8*), "evoked connectome" (*9*) and even "functional connectome" (*10, 11*). For clarity and convenience of writing, we adopt the term "effective connectome", as this study also addressed resting-state fMRI correlation measurements commonly referred to as "functional connectome".

Despite the conceptual similarity, this "effective connectome" is not to be confused with the other measurements also described as "effective" (*12*), which are computationally inferred from resting-state neural activity using methods such as Granger causality and are not involved in this study.

#### 4. Surrogate of Abbott-Chance activation function

The vanilla nonlinearity of Abbott-Chance activation function (*1, 13*) follows the form:

$$H(x) = \frac{x}{1 - e^{-dx}} \quad \text{(S12)}$$

where parameter $d$ is a noise factor of neural population that determines the curvature of the function. When $d$ is large, the function approximates a linear threshold function ReLU, while a smaller

$d$ results in a softened curvature around the threshold. However, this form would cause numerical instability during automatic differentiation in BrainPy (or JAX), particularly around the threshold or for large input values of $x$ (Fig. S22A–B). For example, $H(x)$ exhibits an indeterminate form of 0/0 as $x$ approaches 0, returning NaN (Fig. S22A).

To address this issue, we customized the non-linearity using a piecewise surrogate approach:

$$H(x) = \begin{cases} \dfrac{d}{12}x^2 + 0.5x + \dfrac{1}{d} & \text{if } |x| < 0.1, \\ \max\{0, x\} & \text{if } |x| > 500, \\ \dfrac{x}{1 - e^{-dx}} & \text{otherwise.} \end{cases} \tag{S13}$$

where the function reverts to a linear form for $|x| > 500$, while second-order Taylor approximation is applied within the threshold region defined by $|x| < 0.1$. This customized function ensures stable automatic differentiation (Fig. S22C–D).

To maintain consistency with previous studies (*5, 14, 15*), this study decided to preserve Abbott-Chance activation function in its original form as closely as possible, while ensuring its compatibility with automatic differentiation. For future convenience of implementation, the Softplus function can be considered as a candidate nonlinearity (*16*), given its widespread use as a soft-ReLU-shaped activation function in BrainPy or PyTorch:

$$\text{Softplus}(x) = \begin{cases} x & \text{if } \beta x > 20, \\ \dfrac{1}{\beta}\ln(1 + e^{\beta x}) & \text{otherwise.} \end{cases} \tag{S14}$$

where only one piecewise surrogate is applied for large input values of $x$ for numerical stability. $\beta$ determines the curvature around the threshold, providing a shape similar to the Abbott-Chance nonlinearity with $\beta = \ln(2) \times d$ (Fig. S22E–H).

#### 5. Analytical relationships between MFM parameters and connectomic node strength

We relate the effective connectome (EC), the functional connectome (FC) and their node strengths (NS) to the parameters ($w_i$, $I_i$ and $\sigma_i$) of the mean-field model (MFM). We start by linearizing the nonlinear dynamics around a stable equilibrium (*4, 5*). Using Green function of the linearized system, we derive EC and resting covariance, relate EC strength to recurrence $w_i$ and identify the compensating background input $I_i$ that preserves the equilibrium. Under homogeneous noise, covariance closely resembles EC, differing only by a bounded correction arising from asymmetry in the system's Jacobian. This close correspondence motivates the introduction of heterogeneous noise ($\sigma_i$), which can substantially reshape covariance and the resulting Pearson-normalized FC, as illustrated analytically by a four-node example demonstrating noise-induced suppression.

##### 5.1 Linearization of MFM dynamics

Let $C_{ij} = G\,\mathrm{SC}_{ij}$ denote the scaled structural connection from node $j$ to node $i$, where $G \geq 0$ is the global coupling. The matrix $C$ is fixed throughout and satisfies $C = C^\top$, $C_{ij} \geq 0$ and $C_{ii} = 0$. We also assume $J_N > 0$. We use the transfer function $H$ defined in main-text Eq. (6). The deterministic drift and input current are

$$f_i(\boldsymbol{S}) = -\frac{S_i}{\tau_s} + \gamma(1 - S_i)H(x_i), \tag{S15}$$

$$x_i = J_N w_i S_i + J_N \sum_j C_{ij} S_j + I_i. \tag{S16}$$

Write $\mathbf{I}$ for the $N \times N$ identity matrix. Resting-state and stimulation experiments use separate equations: resting activity satisfies $\mathrm{d}S_i = f_i(\boldsymbol{S})\mathrm{d}t + \sigma_i \mathrm{d}v_i$, whereas the post-pulse response satisfies $\dot{S}_i = f_i(\boldsymbol{S})$ without noise. The $v_i(t)$ are independent standard Wiener processes, collected in $\boldsymbol{v}(t) = (v_1(t), \ldots, v_N(t))^\top$, whose increments satisfy

$$\mathbb{E}[\mathrm{d}v_i \mathrm{d}v_j] = \mathbf{I}_{ij}\mathrm{d}t. \tag{S17}$$

Thus $\sigma_i$ is a noise amplitude and $\sigma_i^2$ is its variance rate.

At a deterministic equilibrium, $\boldsymbol{f}(\boldsymbol{S}^*) = 0$, so

$$S_i^* = \frac{\tau_s \gamma H(x_i^*)}{1 + \tau_s \gamma H(x_i^*)}, \qquad x_i^* = J_N w_i S_i^* + J_N \sum_j C_{ij} S_j^* + I_i. \tag{S18}$$

Writing $\boldsymbol{s} = \boldsymbol{S} - \boldsymbol{S}^*$ and differentiating Eq. (S15) at the equilibrium gives $\boldsymbol{f}(\boldsymbol{S}^* + \boldsymbol{s}) = J\boldsymbol{s} + O(\|\boldsymbol{s}\|^2)$. Retaining only the first-order terms yields the linearized resting-state dynamics that determine FC:

$$\boxed{\mathrm{d}\boldsymbol{s} = J\boldsymbol{s}\mathrm{d}t + D_\sigma \mathrm{d}\boldsymbol{v}.} \tag{S19}$$

Here $D_a = \mathrm{diag}(a_1, \ldots, a_N)$ denotes the diagonal matrix associated with a vector $\boldsymbol{a}$. The linearized stimulation dynamics contain a deterministic input $\boldsymbol{b}(t)$ and determine EC:

$$\boxed{\frac{\mathrm{d}\boldsymbol{s}}{\mathrm{d}t} = J\boldsymbol{s} + \boldsymbol{b}(t).} \tag{S20}$$

The Jacobian is

$$\begin{aligned} J_{ij} &= -\ell_i \mathbf{I}_{ij} + J_N g_i (w_i \mathbf{I}_{ij} + C_{ij}), \\ J &= -D_\ell + J_N D_g (D_w + C), \end{aligned} \tag{S21}$$

where the gain and decay rate are

$$g_i = \gamma(1 - S_i^*)H'(x_i^*) > 0, \qquad \ell_i = \tau_s^{-1} + \gamma H(x_i^*) = \frac{1}{\tau_s(1 - S_i^*)}. \tag{S22}$$

For a unit-normalized state pulse at node $j$, $\boldsymbol{b}(t) = \boldsymbol{e}_j \delta(t)$, where $\boldsymbol{e}_j$ is the $j$th unit vector and $\delta(t)$ is the unit-area Dirac pulse. Equivalently, $\boldsymbol{s}(0^+) - \boldsymbol{s}(0^-) = \boldsymbol{e}_j$; after the pulse, $\boldsymbol{b}(t) = 0$.

**Properties of the Jacobian** The off-diagonal entries of $J$ are nonnegative, so $J$ is a Metzler matrix and $e^{Jt}$ is entrywise nonnegative for $t \geq 0$. Moreover,

$$\mathcal{J} := D_g^{-1/2} J D_g^{1/2} = -D_\ell + J_N D_g D_w + J_N D_g^{1/2} C D_g^{1/2} \tag{S23}$$

is symmetric. Thus $J$ is diagonalizable with real eigenvalues. We require $J$ to be Hurwitz, meaning every eigenvalue has negative real part, which is equivalent to $\mathcal{J} \prec 0$ and ensures decay of pulse responses and existence of a stationary covariance. The matrix $J$ itself is symmetric exactly when $C_{ij}(g_i - g_j) = 0$ for all $i, j$. A homogeneous-state equilibrium, defined by $S_i^* = S^*$ for all $i$, implies identical $x_i^*$, $g_i$ and $\ell_i$ across nodes and hence a symmetric $J$. Neither a homogeneous-state equilibrium nor symmetry of $J$ is assumed unless stated.

For later use, diagonalize the symmetric representative as

$$\begin{aligned} &\mathcal{J} = U D_\lambda U^\top, \qquad U^\top U = \mathbf{I}, \qquad \lambda_a < 0, \\ &R = D_g^{1/2} U, \qquad L = D_g^{-1/2} U, \qquad J = R D_\lambda L^\top, \qquad L^\top R = \mathbf{I}. \end{aligned} \tag{S24}$$

Thus $R$ and $L$ contain the right and left eigenvectors of $J$.

**Green function** The solution of $\dot{\boldsymbol{s}} = J\boldsymbol{s}$ is $\boldsymbol{s}(t) = e^{Jt}\boldsymbol{s}(0^+)$, giving the causal Green function

$$K(t) = \begin{cases} e^{Jt}, & t \geq 0, \\ 0, & t < 0. \end{cases} \tag{S25}$$

For a unit state pulse at node $j$, the deterministic response is

$$\boxed{\boldsymbol{s}^{(j)}(t) = K(t)\boldsymbol{e}_j, \qquad t \geq 0.} \tag{S26}$$

Hence $K_{ij}(t)$ is the response at node $i$ to stimulation at node $j$. Resting activity initialized at equilibrium instead satisfies

$$\boxed{\boldsymbol{s}(t) = \int_0^t K(t-r)D_\sigma \mathrm{d}\boldsymbol{v}(r).} \tag{S27}$$

Both experiments therefore share the same propagation kernel.

##### 5.2 Analytical solutions for EC, resting covariance and FC

The EC entry $E_{ij} = \int_0^\infty K_{ij}(t)\mathrm{d}t$ represents the time-integrated response at node $i$ to a pulse applied at node $j$. Since $K(t) \to 0$ under stability,

$$J\int_0^\infty K(t)\mathrm{d}t = K(\infty) - K(0^+) = -\mathbf{I}, \qquad \boxed{E = -J^{-1}.} \tag{S28}$$

Simulated EC is measured over a finite post-stimulation window of 10–200 ms, whereas integrating the theoretical response to infinite time yields the compact expression in Eq. (S28) and facilitates analytical treatment. The strong correlation between analytical and simulated EC (main-text Fig. 7A) supports this infinite-time solution as a useful approximation to the finite-window measure.

Independence of the noise increments in Eq. (S27) gives the finite-time covariance and its stationary limit:

$$\begin{aligned} P(t) &= \mathbb{E}[\boldsymbol{s}(t)\boldsymbol{s}(t)^\top] = \int_0^t K(t-r)D_\sigma^2 K(t-r)^\top \mathrm{d}r, \\ P &= \int_0^\infty K(t)D_\sigma^2 K(t)^\top \mathrm{d}t. \end{aligned} \tag{S29}$$

Differentiating the integrand and integrating over time yields the Lyapunov equation, same as in (*4, 5*):

$$\boxed{JP + PJ^\top + D_\sigma^2 = 0.} \tag{S30}$$

For Hurwitz $J$, this solution is unique. The eigenvectors in Eq. (S24) also give an explicit spectral solution. Set $\widetilde{P} = L^\top P L$. The Lyapunov equation becomes

$$\begin{gathered} D_\lambda \widetilde{P} + \widetilde{P} D_\lambda + L^\top D_\sigma^2 L = 0, \\ \widetilde{P}_{ab} = -\frac{(L^\top D_\sigma^2 L)_{ab}}{\lambda_a + \lambda_b}, \qquad P = R \widetilde{P} R^\top. \end{gathered} \tag{S31}$$

Equivalently,

$$P_{ij} = -\sum_{a,b,k} \frac{R_{ia} R_{jb} L_{ka} L_{kb}}{\lambda_a + \lambda_b} \sigma_k^2. \tag{S32}$$

Pearson normalization then gives

$$\mathrm{FC}_{ij} = \frac{P_{ij}}{\sqrt{P_{ii} P_{jj}}}. \tag{S33}$$

For analytical tractability, we derive FC directly from synaptic-gating fluctuations and do not seek an exact analytical expression for BOLD FC under the Balloon–Windkessel hemodynamic model. The strong correlation between analytical FC and FC computed from simulated BOLD signals (main-text Fig. 7B) supports this neural-level solution as a useful approximation to BOLD FC.

With $\mathbf{1} = (1, \ldots, 1)^\top$, afferent node strength is defined throughout this theoretical note as the row sum including the diagonal:

$$\mathbf{NS}_{\mathrm{EC}} = E\mathbf{1}, \qquad \mathbf{NS}_P = P\mathbf{1}, \qquad \mathbf{NS}_{\mathrm{FC}} = \mathrm{FC}\mathbf{1}. \tag{S34}$$

The diagonal-inclusive row sums in Eq. (S34), such as $E\mathbf{1}$, are used only in this theoretical derivation for analytical convenience; all NS values reported in the main text are calculated as means of the available off-diagonal connections.

##### 5.3 Recurrence $w_i$ and EC node strength

**Neumann expansion of EC** To display the contributions of local recurrence and network propagation, collect the net local decay in the diagonal matrix

$$D_h = D_\ell - J_N D_g D_w, \qquad h_i = \ell_i - J_N g_i w_i. \tag{S35}$$

Since $J$ is a Hurwitz Metzler matrix, its diagonal entries satisfy $J_{ii} < 0$. With $C_{ii} = 0$, we have $J_{ii} = -\ell_i + J_N g_i w_i = -h_i$, so $h_i = \ell_i - J_N g_i w_i > 0$ for every node. Thus $D_h$ is positive definite at every stable operating point considered here, including when some recurrence parameters are negative. The exact EC matrix is

$$\begin{aligned} E &= \left[D_\ell - J_N D_g C - J_N D_g D_w\right]^{-1} \\ &= (D_h - J_N D_g C)^{-1} = (\mathbf{I} - T)^{-1} D_h^{-1}, \\ T &= J_N D_h^{-1} D_g C = J_N (D_\ell - J_N D_g D_w)^{-1} D_g C. \end{aligned} \tag{S36}$$

The matrix $T$ represents one network-propagation step, including the receiving gain and local recurrence. It is entrywise nonnegative and is similar to $B = J_N D_h^{-1/2} D_g^{1/2} C D_g^{1/2} D_h^{-1/2}$. Since $\mathbf{I} - B = -D_h^{-1/2} \mathcal{J} D_h^{-1/2} \succ 0$,

$$\rho(T) = \rho(B) = \lambda_{\max}(B) < 1, \tag{S37}$$

where $\rho(\cdot)$ denotes the spectral radius and $\lambda_{\max}(\cdot)$ denotes the largest eigenvalue. The convergent Neumann expansion therefore takes the matrix form

$$\boxed{E = \sum_{m=0}^{\infty} T^m D_h^{-1}.} \tag{S38}$$

The identity term gives the local response and $T, T^2, T^3, \ldots$ add paths traversing one, two, three, or more structural connections. Local recurrence is included to all orders through $D_h^{-1}$.

**EC node strength and recurrence $w_i$** The EC node-strength vector is $\mathbf{NS}_{\text{EC}} = E\mathbf{1}$. Multiplying Eq. (S38) by $\mathbf{1}$ yields the following expressions under three different conditions.

**(i) Homogeneous recurrence and homogeneous-state equilibrium.** If $w_i = w$ and $S_i^* = S^*$, then $g_i = g$, $\ell_i = \ell$ and $D_h = (\ell - J_N g w)\mathbf{I}$. With $\mathbf{NS}_C = C\mathbf{1}$, a first-order truncation with error $O(\|C\|^2)$ gives

$$\boxed{\text{NS}_{\text{EC}}(i) = \frac{1}{\ell - J_N g w} + \frac{J_N g}{(\ell - J_N g w)^2} \text{NS}_C(i) + O(\|C\|^2).} \tag{S39}$$

Thus EC strength is an increasing affine function of structural node strength at first order.

**(ii) Heterogeneous recurrence and homogeneous-state equilibrium.** If $w_i$ varies but $S_i^* = S^*$, the common gain and decay rate remain $g, \ell$, while $D_h = \ell\mathbf{I} - J_N g D_w$ gives different local responses:

$$\boxed{\text{NS}_{\text{EC}}(i) = \frac{1}{\ell - J_N g w_i} + \frac{J_N g}{\ell - J_N g w_i} \sum_j \frac{C_{ij}}{\ell - J_N g w_j} + O(\|C\|^2).} \tag{S40}$$

Recurrence now weights both the receiving node and the nodes supplying its input, so $\mathrm{NS}_C(i)$ alone no longer determines EC strength. At fixed $S^*$, increasing $w_i$ reduces the net local decay $\ell - J_N g w_i$ and increases the weights of both the local response and propagation terms.

**(iii) Heterogeneous recurrence and heterogeneous-state equilibrium.** When both $w_i$ and $S_i^*$ vary, the gains and decay rates also vary:

$$\boxed{\mathrm{NS}_{\mathrm{EC}}(i) = \frac{1}{\ell_i - J_N g_i w_i} + \frac{J_N g_i}{\ell_i - J_N g_i w_i} \sum_j \frac{C_{ij}}{\ell_j - J_N g_j w_j} + O(\|C\|^2).} \tag{S41}$$

Although regional differences in $g_i$ and $\ell_i$ modulate the magnitude and precise form of this relationship through the local response and propagation weights, the positive dependence of EC node strength on local recurrence persists at a fixed equilibrium: increasing $w_i$ reduces $h_i = \ell_i - J_N g_i w_i$ and thereby increases $\mathrm{NS}_{\mathrm{EC}}(i)$, provided that $g_i > 0$ and stability is maintained.

Excluding the diagonal removes the zeroth-order term $h_i^{-1}$ and all higher-order contributions to $E_{ii}$, while the remaining nonzero off-diagonal terms in row $i$ of the Neumann expansion still increase with $w_i$ at a fixed equilibrium, preserving the positive recurrence–NS relationship while stability is maintained.

##### 5.4 Recurrence $w_i$ and compensating background input $I_i$

To vary recurrence $w_i$ while keeping a chosen equilibrium vector $\boldsymbol{S}^*$ fixed, we compensate for the change in recurrence by adjusting the background input $I_i$. Solving the equilibrium input-current relation in Eq. (S18) for $I_i$ gives

$$\boxed{\begin{aligned} I_i &= x_i^* - J_N w_i S_i^* - J_N \sum_j C_{ij} S_j^*, \\ \left.\frac{\partial I_i}{\partial w_i}\right|_{S^*} &= -J_N S_i^* < 0. \end{aligned}} \tag{S42}$$

Thus greater recurrence requires less background input to maintain the same equilibrium. For a finite change at node $i$, $\Delta I_i = -J_N S_i^* \Delta w_i$. This compensation makes it possible to tune the Jacobian $J$ and hence the system dynamics by varying recurrence without shifting the equilibrium.

The negative slope applies to changes within a node. Across nodes, both $S_i^*$ and the intercept $x_i^* - J_N \sum_j C_{ij} S_j^*$ may differ, so a negative cross-node sample correlation does not follow automatically.

If we additionally assume a homogeneous-state equilibrium $S_i^* = S^*$ and equal structural row sums $c_i = \sum_j C_{ij}$, the relation simplifies to

$$I_i = x^* - J_N S^* (w_i + c_i) \tag{S43}$$

which gives an exact negative affine relation between $I_i$ and $w_i$ across nodes.

##### 5.5 EC–covariance relation under homogeneous noise

Let $\sigma_i = \sigma > 0$, so $D_\sigma^2 = \sigma^2 \mathbf{I}$. In this case all noise sources receive the same variance weight. The remaining covariance structure is determined by the same dynamics as EC.

**Symmetric Jacobian: exact covariance–EC proportionality** If additionally $J = J^\top$, the covariance reduces to

$$\boxed{P = \sigma^2 \int_0^\infty e^{2Jt} \mathrm{d}t = -\frac{\sigma^2}{2} J^{-1} = \frac{\sigma^2}{2} E.} \tag{S44}$$

**Nonsymmetric Jacobian: the asymmetry correction** For general $J$, set $\Xi = (JP - PJ^\top)/\sigma^2$, so $\Xi^\top = -\Xi$. The Lyapunov equation gives $JP = (\sigma^2/2)(-\mathbf{I} + \Xi)$ and therefore

$$\boxed{P = -\frac{\sigma^2}{2} J^{-1} + \frac{\sigma^2}{2} J^{-1} \Xi = \frac{\sigma^2}{2} E(\mathbf{I} - \Xi).} \tag{S45}$$

The correction is determined entirely by the dynamics:

$$J\Xi + \Xi J^\top = -(J - J^\top), \quad \Xi = \int_0^\infty e^{Jt} (J - J^\top) e^{J^\top t} \mathrm{d}t. \tag{S46}$$

The diagonal terms cancel from $J - J^\top$, so Jacobian asymmetry is determined by regional gain differences weighted by structural connectivity: $(J - J^\top)_{ij} = J_N (g_i - g_j) C_{ij}$. At a fixed equilibrium, varying $w_i$ leaves $J - J^\top$ unchanged but can still alter $\Xi$ through the propagation factors $e^{Jt}$ in Eq. (S46).

**A Frobenius-norm bound on the asymmetry correction** Let $\|\cdot\|_F$ and $\|\cdot\|_2$ denote the Frobenius and spectral norms, respectively. Define $g_{\min} = \min_i g_i > 0$, $g_{\max} = \max_i g_i$ and $\kappa_g = g_{\max}/g_{\min}$. Using the eigenvectors in Eq. (S24), set $X = L^\top \Xi L$ and $M = L^\top L = U^\top D_g^{-1} U$. The equation for $\Xi$ gives

$$X_{ab} = \frac{\lambda_b - \lambda_a}{\lambda_a + \lambda_b} M_{ab}. \tag{S47}$$

Because $\lambda_a < 0$, the coefficient has absolute value at most one and $X_{aa} = 0$. Hence $\|X\|_F \leq \|M - c\mathbf{I}\|_F$. Choosing $c = (g_{\min}^{-1} + g_{\max}^{-1})/2$ and using $\Xi = RXR^{\top}$, $\|R\|_2^2 = g_{\max}$ and orthogonality of $U$ gives

$$
\begin{aligned}
\|\Xi\|_F &\leq g_{\max}\|X\|_F \leq g_{\max}\|M - c\mathbf{I}\|_F \\
&= g_{\max}\left[\sum_i (g_i^{-1} - c)^2\right]^{1/2} \leq \frac{\sqrt{N}}{2}(\kappa_g - 1).
\end{aligned} \tag{S48}
$$

As $\kappa_g \to 1$, both the Jacobian asymmetry $J - J^{\top}$ and the correction $\Xi$ vanish, so $P$ approaches $(\sigma^2/2)E$.

##### 5.6 Heterogeneous noise can substantially alter the EC–covariance relation

For positive regional noise amplitudes $\sigma_i > 0$, the homogeneous-noise derivation extends to heterogeneous noise through the rescaling $\widehat{s} = D_\sigma^{-1}s$, which gives

$$
\begin{gathered}
\widehat{J} = D_\sigma^{-1} J D_\sigma, \qquad \widehat{P} = D_\sigma^{-1} P D_\sigma^{-1}, \\
\widehat{J}\widehat{P} + \widehat{P}\widehat{J}^{\top} + \mathbf{I} = 0.
\end{gathered} \tag{S49}
$$

The similarity transformation preserves the stability of $J$. Applying Eq. (S45) with unit noise gives

$$
\begin{aligned}
\widehat{E} &= -\widehat{J}^{-1} = D_\sigma^{-1} E D_\sigma, & \widehat{\Xi} &= \widehat{J}\widehat{P} - \widehat{P}\widehat{J}^{\top}, \\
\widehat{J}\widehat{\Xi} + \widehat{\Xi}\widehat{J}^{\top} &= -(\widehat{J} - \widehat{J}^{\top}), & \widehat{P} &= \frac{1}{2}\widehat{E}(\mathbf{I} - \widehat{\Xi}).
\end{aligned} \tag{S50}
$$

Transforming back yields the exact covariance:

$$
\boxed{P = \frac{1}{2} E D_\sigma (\mathbf{I} - \widehat{\Xi}) D_\sigma.} \tag{S51}
$$

For the MFM Jacobian, the transformed asymmetry explicitly depends on the regional noise amplitudes:

$$
(\widehat{J} - \widehat{J}^{\top})_{ij} = J_N C_{ij} \frac{g_i \sigma_j^2 - g_j \sigma_i^2}{\sigma_i \sigma_j}. \tag{S52}
$$

Thus at fixed $J$, changing a homogeneous noise amplitude only rescales $P$ (Eq. (S45)), whereas varying regional noise amplitudes directly reweights $P$ through the explicit $D_\sigma$ factors and modifies the correction $\widehat{\Xi}$ in Eq. (S51). Together, these effects allow heterogeneous noise to substantially reshape covariance and FC node-strength patterns while leaving $E$ unchanged.

#### 5.7 Regional noise can suppress or enhance FC node strength

Let $J$ and all other noise amplitudes be held fixed when one noise source is varied. Set $q_k = \sigma_k^2 > 0$ and define the covariance contribution from unit noise variance at node $k$:

$$A^{(k)} = \int_0^\infty K(t)\boldsymbol{e}_k\boldsymbol{e}_k^\top K(t)^\top \mathrm{d}t, \qquad P = \sum_k q_k A^{(k)}. \tag{S53}$$

Each $A^{(k)}$ depends only on $J$. Since $K(t)$ is entrywise nonnegative, $A_{ij}^{(k)} \geq 0$, so $\partial P_{ij}/\partial q_k = A_{ij}^{(k)} \geq 0$. The exact FC is

$$\mathrm{FC}_{ij} = \frac{P_{ij}}{\sqrt{P_{ii}P_{jj}}} = \frac{\sum_k q_k A_{ij}^{(k)}}{\sqrt{\left(\sum_k q_k A_{ii}^{(k)}\right)\left(\sum_k q_k A_{jj}^{(k)}\right)}} \tag{S54}$$

**Noise sensitivities and their signs** Differentiating the Pearson normalization gives

$$\frac{\partial \mathrm{FC}_{ij}}{\partial q_k} = \frac{A_{ij}^{(k)}}{\sqrt{P_{ii}P_{jj}}} - \frac{\mathrm{FC}_{ij}}{2}\left(\frac{A_{ii}^{(k)}}{P_{ii}} + \frac{A_{jj}^{(k)}}{P_{jj}}\right). \tag{S55}$$

The first term increases shared covariance, whereas the second accounts for the increase in the two variances used for normalization. Thus FC increases when relative covariance growth exceeds the mean relative variance growth and decreases when it falls below it.

For the effect of a node's own noise on its FC strength, summing the off-diagonal edge sensitivities yields

$$\boxed{\begin{aligned} \frac{\partial \mathrm{NS}_{\mathrm{FC}}(i)}{\partial q_i} &= \sum_{j\neq i}\left[\frac{A_{ij}^{(i)}}{\sqrt{P_{ii}P_{jj}}} - \frac{\mathrm{FC}_{ij}}{2}\left(\frac{A_{ii}^{(i)}}{P_{ii}} + \frac{A_{jj}^{(i)}}{P_{jj}}\right)\right], \\ \frac{\partial \mathrm{NS}_{\mathrm{FC}}(i)}{\partial \sigma_i} &= 2\sigma_i \frac{\partial \mathrm{NS}_{\mathrm{FC}}(i)}{\partial q_i}. \end{aligned}} \tag{S56}$$

The diagonal contributes no derivative because $\mathrm{FC}_{ii} = 1$. The node-strength derivative aggregates the competing contributions from all off-diagonal edges, so its sign depends on the FC-weighted balance between relative covariance growth and mean relative variance growth. Increasing a node's noise can therefore either enhance or suppress its FC node strength, depending on which contribution dominates overall. The following four-node example illustrates a case of FC node strength suppression with increased regional noise.

**Four-node toy example** We consider a four-node toy model with nodes labeled $1, 2, 3, 4$. The structural matrix is

$$\mathrm{SC} = \begin{pmatrix} 0 & 0.1 & 0.6 & 1.2 \\ 0.1 & 0 & 1 & 1 \\ 0.6 & 1 & 0 & 1 \\ 1.2 & 1 & 1 & 0 \end{pmatrix}, \tag{S57}$$

with the remaining parameters given by

$$\begin{gathered} (a, b, d) = (270, 108, 0.154), \qquad \tau_s = 0.1, \quad \gamma = 0.641, \quad J_N = 0.2609, \\ G = 8, \qquad w_i^{(0)} = -20, \qquad I_i^{(0)} = 0.4, \qquad \sigma_i^{(0)} = \sigma_0 = 0.05. \end{gathered} \tag{S58}$$

The resulting equilibrium is

$$\boldsymbol{S}^* \simeq (0.283,\ 0.305,\ 0.344,\ 0.391)^\mathsf{T}. \tag{S59}$$

**Stage (i): recurrence with input compensation.** All noise amplitudes remain $\sigma_0$. Increase $w_1$ from $-20$ to $-18$ while adjusting the background input according to $I_1(w_1) = 0.4 - J_N(w_1 + 20)S_1^*$ and keeping the other regional parameters fixed. This preserves the full equilibrium and increases the EC and FC strengths in this example. Stability holds throughout: $\mathcal{J}(w_1) \preceq \mathcal{J}(-18) \prec 0$ for $-20 \leq w_1 \leq -18$.

**Stage (ii): noise at fixed recurrence and input.** Keep $w_1 = -18$ and its compensated input $I_1$ fixed while increasing only $\sigma_1$ from 0.05 to 0.20.

Table S3 summarizes the two stages. In stage (i), increasing $w_1$ from $-20$ to $-18$ with input compensation raises $\mathrm{NS}_{\mathrm{EC}}(1)$ from 0.064 to 0.134 and $\mathrm{NS}_{\mathrm{FC}}(1)$ from 3.579 to 3.775. In stage (ii), raising $\sigma_1$ from 0.05 to 0.125 and 0.20 reduces $\mathrm{NS}_{\mathrm{FC}}(1)$ from 3.775 to 3.652 and 3.618, respectively, while $\mathrm{NS}_{\mathrm{EC}}(1)$ remains 0.134. These values establish that regional noise can suppress FC strength at fixed EC.

### Supplementary Tables

| Index | Abbreviation | Index | Abbreviation |
|---|---|---|---|
| 1 | ctx-lh-bankssts | 35 | ctx-rh-bankssts |
| 2 | ctx-lh-caudalanteriorcingulate | 36 | ctx-rh-caudalanteriorcingulate |
| 3 | ctx-lh-caudalmiddlefrontal | 37 | ctx-rh-caudalmiddlefrontal |
| 4 | ctx-lh-cuneus | 38 | ctx-rh-cuneus |
| 5 | ctx-lh-entorhinal | 39 | ctx-rh-entorhinal |
| 6 | ctx-lh-fusiform | 40 | ctx-rh-fusiform |
| 7 | ctx-lh-inferiorparietal | 41 | ctx-rh-inferiorparietal |
| 8 | ctx-lh-inferiortemporal | 42 | ctx-rh-inferiortemporal |
| 9 | ctx-lh-isthmuscingulate | 43 | ctx-rh-isthmuscingulate |
| 10 | ctx-lh-lateraloccipital | 44 | ctx-rh-lateraloccipital |
| 11 | ctx-lh-lateralorbitofrontal | 45 | ctx-rh-lateralorbitofrontal |
| 12 | ctx-lh-lingual | 46 | ctx-rh-lingual |
| 13 | ctx-lh-medialorbitofrontal | 47 | ctx-rh-medialorbitofrontal |
| 14 | ctx-lh-middletemporal | 48 | ctx-rh-middletemporal |
| 15 | ctx-lh-parahippocampal | 49 | ctx-rh-parahippocampal |
| 16 | ctx-lh-paracentral | 50 | ctx-rh-paracentral |
| 17 | ctx-lh-parsopercularis | 51 | ctx-rh-parsopercularis |
| 18 | ctx-lh-parsorbitalis | 52 | ctx-rh-parsorbitalis |
| 19 | ctx-lh-parstriangularis | 53 | ctx-rh-parstriangularis |
| 20 | ctx-lh-pericalcarine | 54 | ctx-rh-pericalcarine |
| 21 | ctx-lh-postcentral | 55 | ctx-rh-postcentral |
| 22 | ctx-lh-posteriorcingulate | 56 | ctx-rh-posteriorcingulate |
| 23 | ctx-lh-precentral | 57 | ctx-rh-precentral |
| 24 | ctx-lh-precuneus | 58 | ctx-rh-precuneus |
| 25 | ctx-lh-rostralanteriorcingulate | 59 | ctx-rh-rostralanteriorcingulate |
| 26 | ctx-lh-rostralmiddlefrontal | 60 | ctx-rh-rostralmiddlefrontal |
| 27 | ctx-lh-superiorfrontal | 61 | ctx-rh-superiorfrontal |
| 28 | ctx-lh-superiorparietal | 62 | ctx-rh-superiorparietal |
| 29 | ctx-lh-superiortemporal | 63 | ctx-rh-superiortemporal |
| 30 | ctx-lh-supramarginal | 64 | ctx-rh-supramarginal |
| 31 | ctx-lh-frontalpole | 65 | ctx-rh-frontalpole |
| 32 | ctx-lh-temporalpole | 66 | ctx-rh-temporalpole |
| 33 | ctx-lh-transversetemporal | 67 | ctx-rh-transversetemporal |
| 34 | ctx-lh-insula | 68 | ctx-rh-insula |

**Table S1:** Cortical regions of Desikan-Killiany parcellation. ctx = cortex, lh = left hemisphere, rh = right hemisphere.

| **Stage** | **Setting** | **Two-stage EC–FC training** | **FC-only training** | **Hybrid EC–FC training** |
|---|---|---|---|---|
| Init. | Initial seeds | $\{G_{\text{init}}=1\} \times \{w_{\text{init}}=-150{:}1{:}20\} \times \{I_{\text{init}}=0\} \times \{\sigma_{\text{init}}=10^{-5}\}$ | | |
| 1 | Loss | EC loss | FC loss | Hybrid EC–FC loss |
| | Parameters to train | $\{G, w_i, I_i\}$ | $\{G, w_i, I_i, \sigma_i\}$ | $\{G, w_i, I_i, \sigma_i\}$ |
| | Batch size | 68 | 68 | 68 |
| | Epochs | 10,000 | 10,000 | 10,000 |
| | Base learning rate (LR) | 0.05 | 0.05 | 0.05 |
| | LR multiplier for $w_i$ | ×10 | ×1 | ×1 |
| | Loss ratio $\alpha$ | – | – | $\{0.1, 0.25, 0.5, 0.75, 0.9\}$ |
| 2 | Loss | FC loss | – | – |
| | Parameters to train | $\{\sigma_i\}$ | – | – |
| | Batch size | 68 | – | – |
| | Epochs | 1,000 | – | – |
| | Base learning rate (LR) | 0.1 | – | – |
| | LR multiplier for $w_i$ | – | – | – |

**Table S2:** Settings for two-stage EC–FC, FC-only and hybrid EC–FC training. Initialization is homogeneous across cortical regions; −150:1:20 denotes initial recurrent strengths from −150 to 20 in unit increments. FC-only and hybrid training each use a single stage; only two-stage training includes Stage 2, in which $\sigma_i$ is optimized while the EC-trained $G$, $w_i$ and $I_i$ remain fixed. The learning-rate multiplier is the ratio of the learning rate for $w_i$ to the base learning rate; ×10 denotes a tenfold increase and ×1 denotes the base learning rate. In hybrid training, the loss ratio $\alpha$ sets the EC and FC loss coefficients to $\alpha$ and $1 - \alpha$, respectively. The set of $\alpha$ values indicates that hybrid training was performed separately for every listed value. Intrinsic noise is omitted when simulating EC. Dashes indicate settings that are not applicable.

| Stage | $w_1$ | $I_1$ | $\sigma_1$ | $NS_{EC}(1)$ | $NS_{FC}(1)$ |
|---|---|---|---|---|---|
| (i) | -20.0 | 0.400 | 0.050 | **0.064** | **3.579** |
| (i) | **-19.0** | **0.326** | 0.050 | **0.086** | **3.672** |
| (i) | **-18.0** | **0.252** | 0.050 | **0.134** | **3.775** |
| (ii) | -18.0 | 0.252 | **0.125** | - | **3.652** |
| (ii) | -18.0 | 0.252 | **0.200** | - | **3.618** |

**Table S3:** Increasing recurrence with input compensation at node 1 of a four-node example model raises EC and FC strengths in stage (i); increasing its noise amplitude then reduces FC strength at unchanged EC in stage (ii). Dashes denote the unchanged EC strength in stage (ii). Bold values highlight the parameters changed in each stage.

### Supplementary Figures

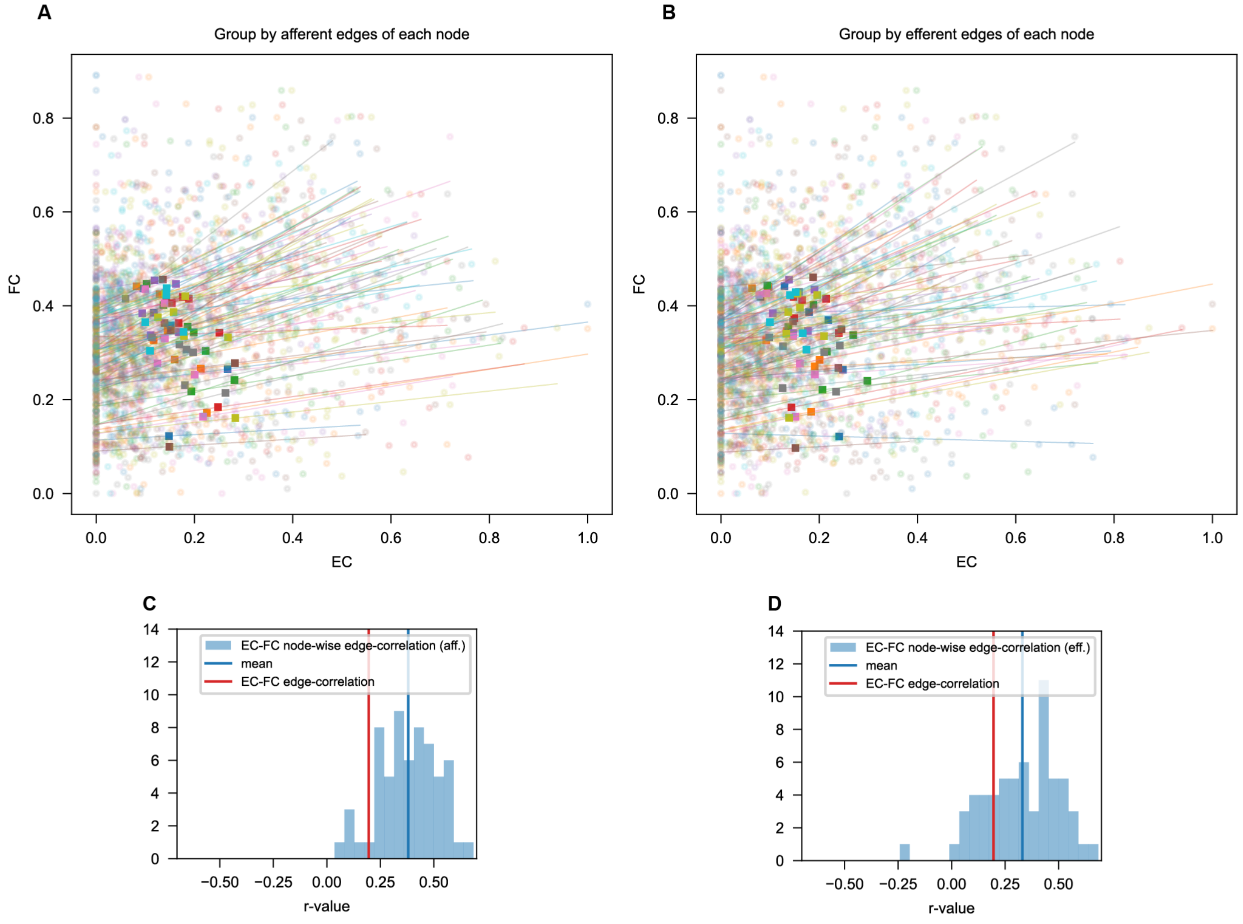

**Fig. S1: Node-wise edge correlations between EC and FC. A,B** FC versus EC for connections grouped by their target node (afferent edges; A) or source node (efferent edges; B). Colours identify nodes, lines show within-node linear fits, and squares indicate the group means, corresponding to node strengths. **C,D** Distributions of within-node Pearson correlations for afferent (panel **C**) and efferent (panel **D**) groups. The blue vertical line marks the mean within-node correlation, and the red line marks the correlation obtained by pooling all measured connections.

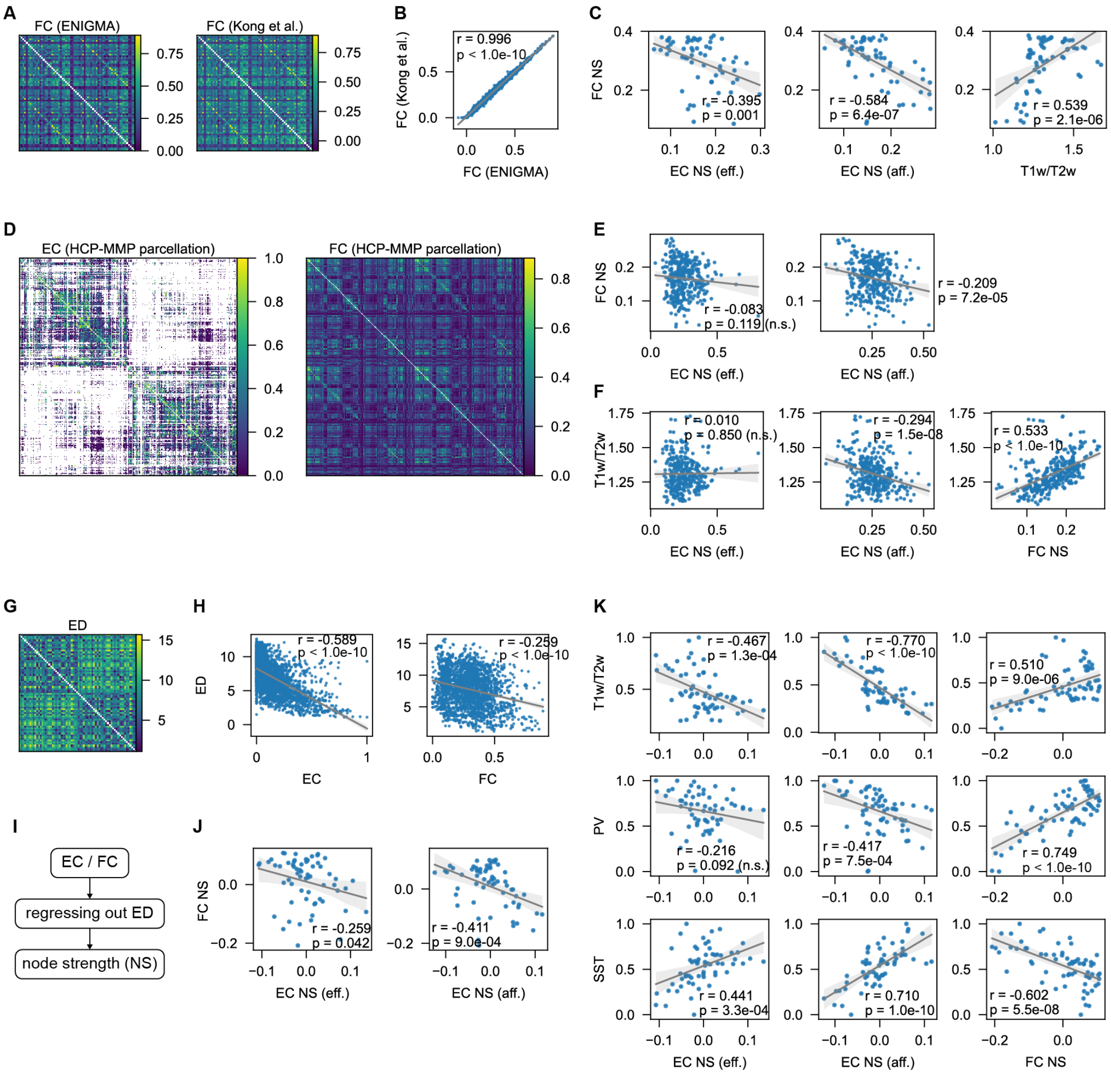

**Fig. S2: Validation of node strength findings across preprocessing pipelines, parcellations, and after regressing out Euclidean distance. A** Group-averaged HCP FC matrices in DK parcellation from the ENIGMA toolbox (left; used in the main text) and Kong et al. (*15*) (right), with independent preprocessing and subject selection. **B** Comparison of their upper-triangular entries. **C** Kong et al. FC node strength (NS) versus efferent EC NS, afferent EC NS and T1w/T2w, from left to right. **D** EC (left) and ENIGMA FC (right) in the 360-region HCP-MMP parcellation (*17*). **E** FC NS versus efferent (left) and afferent (right) EC NS. **F** T1w/T2w versus efferent EC NS, afferent EC NS and FC NS, from left to right. **G** Euclidean distance (ED) between DK parcel centroids (colour bar, cm). **H** ED versus EC (left) and upper-triangular FC entries (right). **I** ED is regressed from EC or

FC entries before recomputing NS from the residuals. **J** ED-regressed FC NS versus efferent (left) and afferent (right) EC NS. **K** T1w/T2w, PV and SST (rows) versus ED-regressed efferent EC NS, afferent EC NS and FC NS (columns). Grey lines show linear fits; Pearson's *r* and *p* are annotated. n.s., not significant; eff., efferent; aff., afferent.

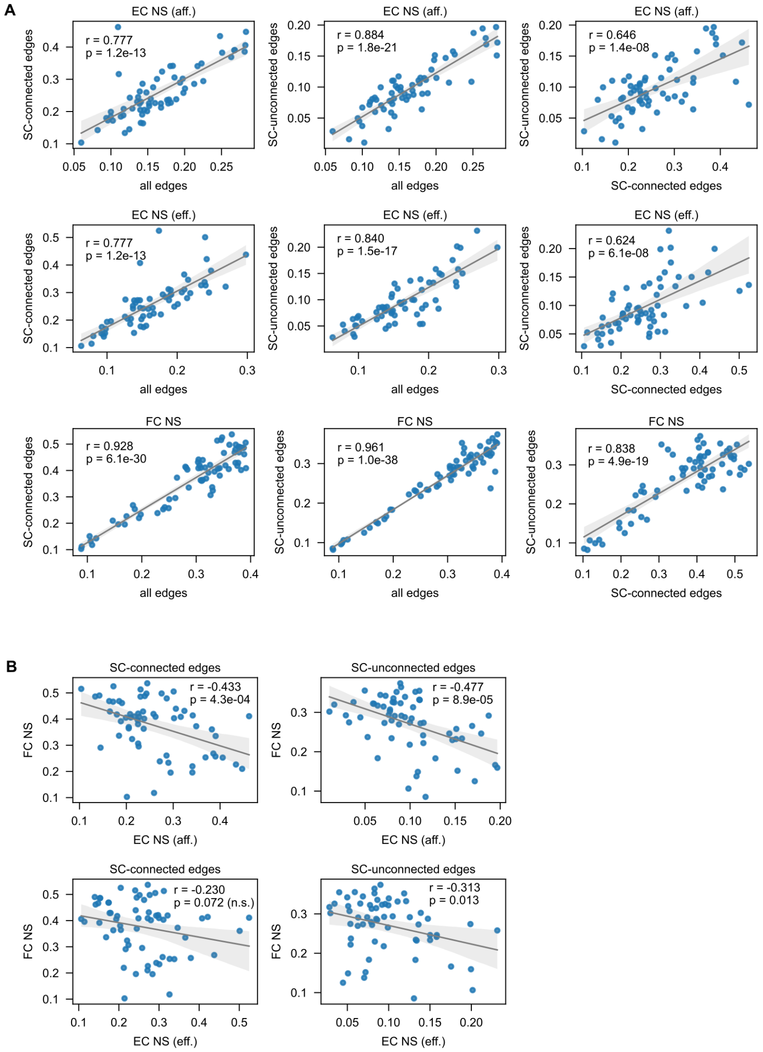

**Fig. S3: Node-strength relationships across structurally connected and unconnected edges. A** Pairwise comparisons of NS calculated from all available edges, SC-connected edges ($SC_{ij} > 0$) and SC-unconnected edges ($SC_{ij} = 0$). Rows show afferent EC, efferent EC and FC; columns compare all versus connected, all versus unconnected, and connected versus unconnected definitions. Self-connections and unmeasured EC entries are excluded. **B** FC NS versus afferent EC NS (top) and efferent EC NS (bottom), calculated using SC-connected (left) or SC-unconnected (right) edges. Afferent EC–FC relationships remain significantly negative in both subsets; the efferent relationship for SC-connected edges is not significant. Points represent regions and grey lines show linear fits. Pearson's $r$ and $p$ are annotated in each panel; n.s., not significant.

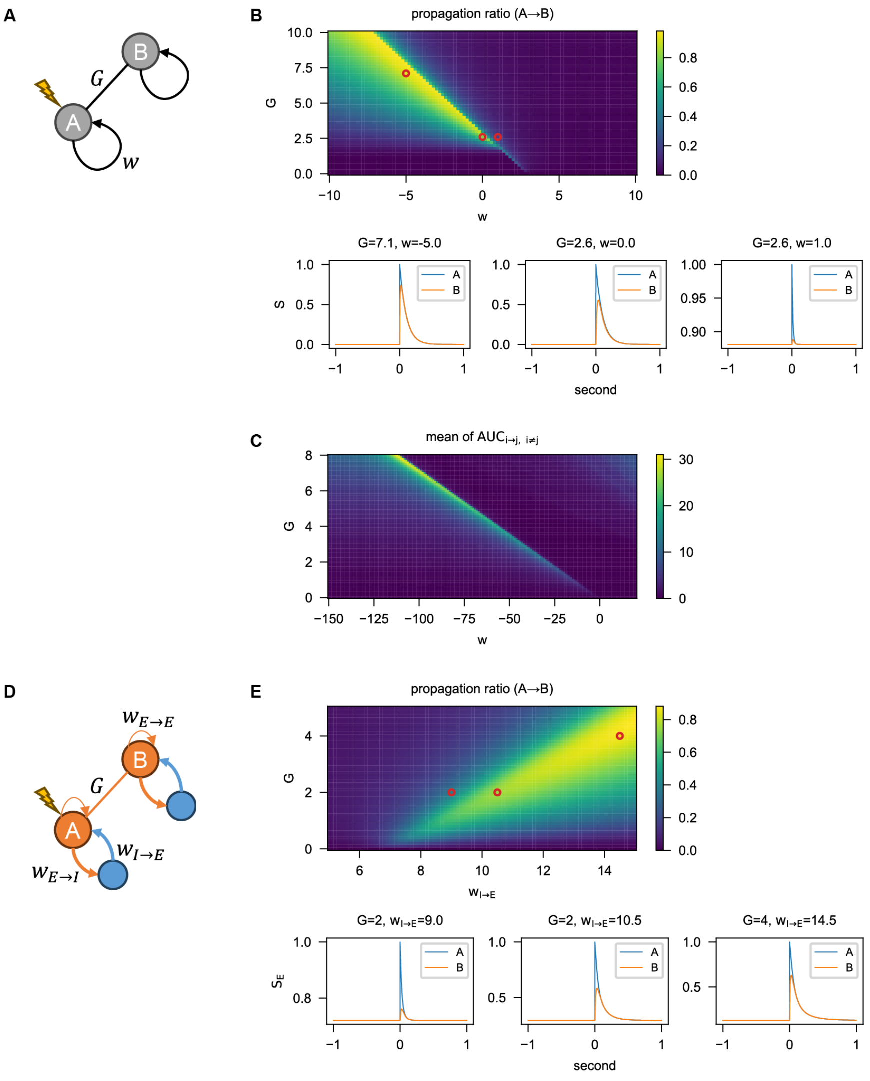

**Fig. S4: Signal propagation regime with negative recurrent strength. A** Model scheme of a two-node example using the mean-field model (MFM), where node A receives a single-pulse stimulus. **B** Heatmap illustrating the propagation ratio (node A⟶B) of the two-node MFM under varying conditions of $G$ and $w$ ($I$ = 0). Propagation ratio (node A⟶B) was obtained when single-pulse stimulating node A and recording node B, and defined as the $AUC_B$ / $AUC_A$. AUC, area under the curve of the evoked response after stimulus onset. Red circles denote three selected parameter sets, with their corresponding time series shown in the bottom panels. **C** Network propagation in the homogeneous model under varying conditions of $G$ and $w$ ($I$ = 0.5). Network propagation was measured when single-pulse stimulating a targeted node i and recording the remaining nodes j (j

$\neq$ i), and defined as the averaged value of $AUC_{i \to j}$ across all target-recording pairs of nodes. **D** Model scheme of a two-node example using the E-I MFM with excitatory (orange) and inhibitory (blue) populations. $w_{E \to E} = 5$, weight of local self-excitation. $w_{E \to I} = 5$, weight of local excitatory-to-inhibitory connection. $w_{I \to E} \in [5, 15]$, weight of local inhibitory-to-excitatory connection. $G \in [0, 5]$, global coupling strength for inter-regional excitatory connection. Other parameters follow the default settings of (*4*). **E** Heatmap illustrating the propagation ratio (node A⟶B) of the two-node E-I MFM under varying conditions of $G$ and $w_{I \to E}$. Red circles in **E** mark the three parameter settings whose response traces are shown below the heatmap.

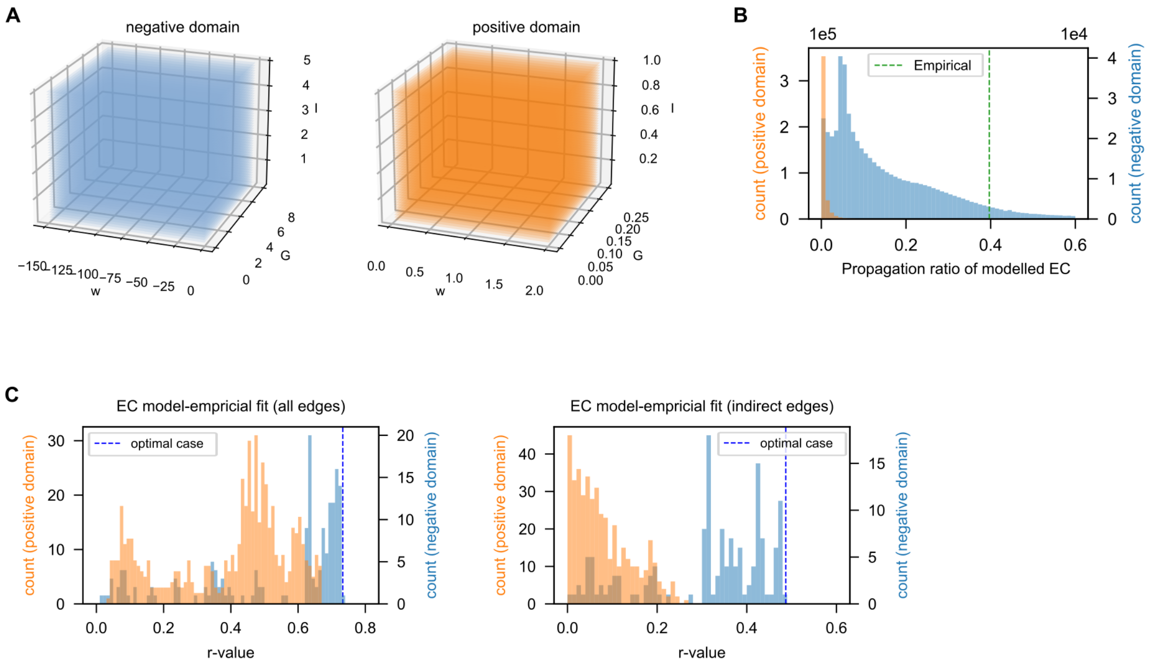

**Fig. S5: Validation of the extended parameter space of negative recurrent strength. A** Visualization of the grid-searched parameter seeds for regional homogeneous models in negative (left panel, blue) and positive (right panel, orange) domains. Negative domain samples $\{G = 0.1 : 0.1 : 8\} \times \{w = -150 : 1 : -1\} \times \{I = 0.1 : 0.1 : 5\}$ with 600,000 seeds in total. Positive domain samples $\{G = 0.005 : 0.005 : 0.25\} \times \{w = 0.01 : 0.01 : 2\} \times \{I = 0.02 : 0.02 : 1\}$ with 500,000 seeds in total. **B** Comparative distributions of the propagation ratio (PR) of modelled EC across sampled parameter seeds from positive and negative domains. PR is defined as the ratio of the sum of weights for indirect connections (SC-unconnected) to the sum of weights for direct connections (SC-connected): $\mathrm{PR} = \frac{\sum \mathrm{EC}_{i,j\in\{\mathrm{SC}_{ij}=0\}}}{\sum \mathrm{EC}_{i,j\in\{\mathrm{SC}_{ij}>0\}}}$ . The negative domain (blue), but not positive domain (orange), yields models capable of reaching the empirical EC's $\mathrm{PR} = 0.397$ (dashed green line). **C** Histograms of model-empirical fit ($r$-value) for all edges (left panel) and indirect edges (i.e., $\mathrm{SC}_{ij} = 0$, right panel), comparing regional heterogeneous models with parameters extended to the negative domain (blue; corresponding to the results in main text Fig. 4A,B) versus bounded in the positive domain (orange). For the positive-domain condition, parameters were enforced via $w_i \longrightarrow |w_i|$ and $I_i \longrightarrow |I_i|$, with training initialized from seeds $\{G = 0.05, 0.25\} \times \{w = 0.1 : 0.1 : 2.5\} \times \{I = 0\}$

and hyperparameters $\{lr = 0.0001, 0.0005, 0.001, 0.005, 0.01, 0.05\} \times \{lr_w = 1, 10\}$. Models with parameters extended into the negative domain (blue) consistently yield higher model-empirical fit (dashed blue, the optimal case), especially for indirect edges, where bounding parameters to the positive domain (orange) fails to achieve comparable performance. The two histogram colours use the corresponding left (orange) and right (blue) count axes.

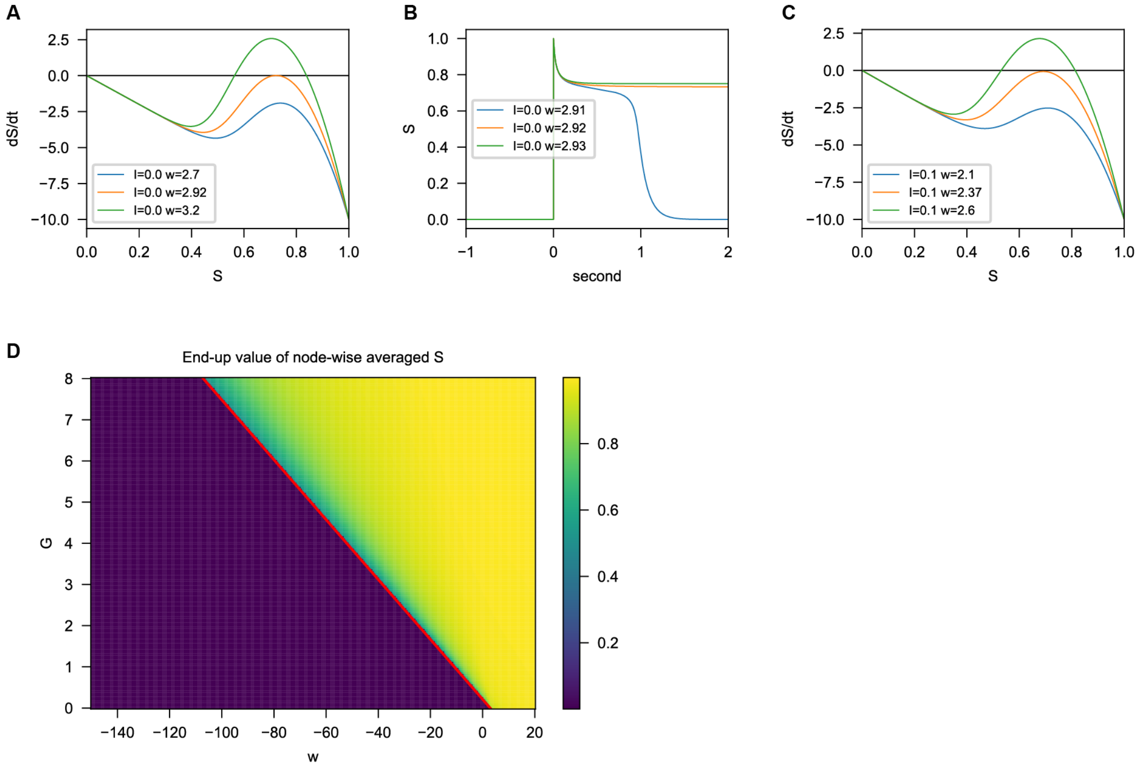

**Fig. S6: Bistability and the bifurcation line in the mean-field model. A** One-dimensional phase portraits ($dS/dt$ versus $S$) for single node, noise-free models with $I = 0$ and increasing recurrent strength ($w = 2.70, 2.92, 3.20$; blue, orange, green). **B** Timeseries of $S$ after a perturbation ($S \longrightarrow 1$ at $t = 0$) for single node, noise-free models with $I = 0$ and increasing recurrent strength ($w = 2.91, 2.92, 2.93$; blue, orange, green). The model with a value of $w$ smaller than the critical $w_0$ fails to end-up with a maintained high-state after perturbation. **C** Same as in **A**, but with $I = 0.1$ and $w = 2.10$, 2.37 and 2.60 (blue, orange and green). **D** Grid search of parameters $G$ and $w$ of homogeneous, noise-free models with $I = 0$. The heatmap shows the end-up values after the perturbation same as in **B** of node-wise averaged $S$ under every parameter combination. The bifurcation line (red; $w + \lambda G = w_0$, with $w_0 = 2.9185$ and SC spectral radius $\lambda = 13.7563$) partitions low- and high-state regimes, serving as a necessary condition for bistability.

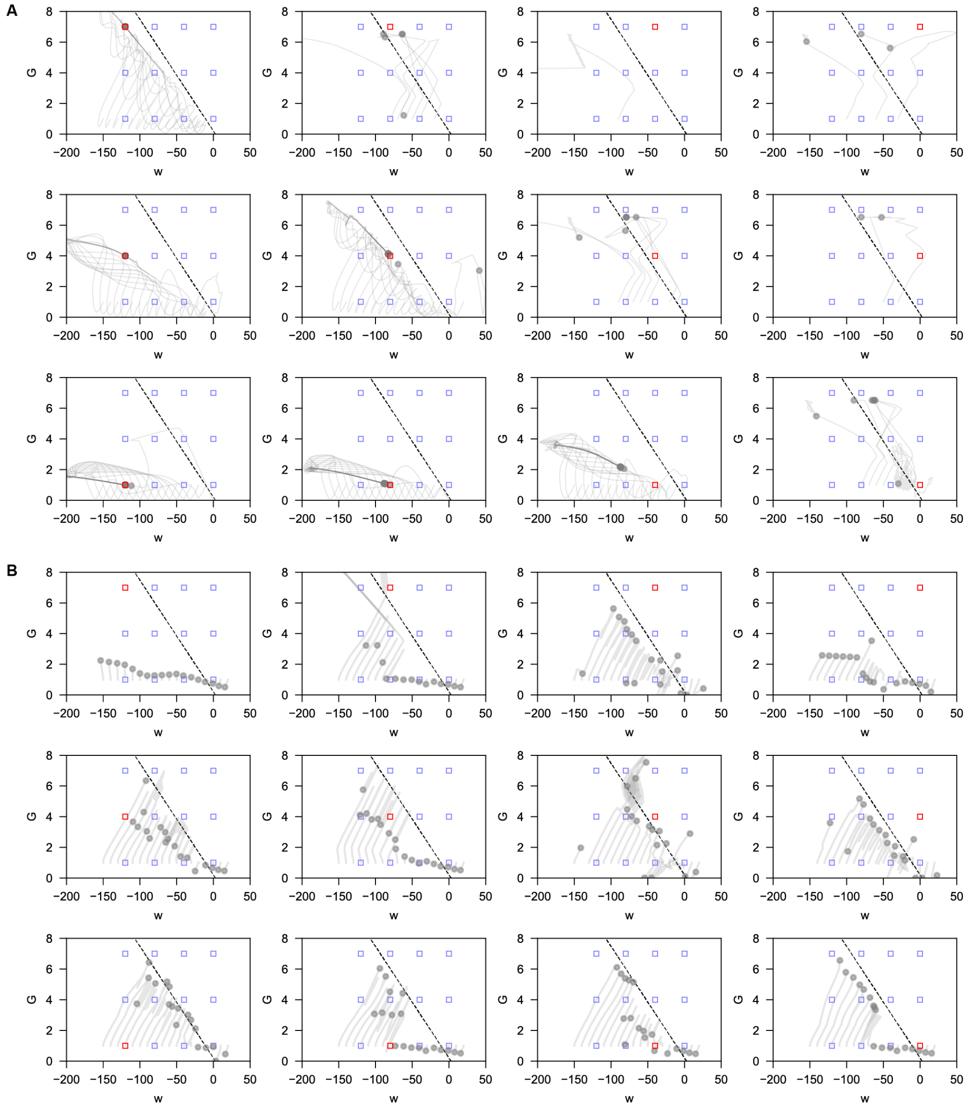

**Fig. S7: Parameter recovery from synthetic connectomes generated by homogeneous models.** **A** EC fitting of homogeneous models (i.e., scalar $w$ and $I$) on synthetic EC matrices derived from predefined ground-truth parameters. In each panel, the ground-truth parameters (red square) were drawn from a grid of $\{G = 1, 4, 7\} \times \{w = \text{-}120, \text{-}80, \text{-}40, 0\} \times \{I = 1\}$ (blue squares). A synthetic EC matrix was generated with a homogeneous model using the ground-truth parameters, and served as the training objective. Training was applied to optimize homogeneous models as initialized at seeds of $\{G = 1\} \times \{w = \text{-}150{:}10{:}20\} \times \{I = 0\}$, with hyperparameters lr = 0.5, $\mathrm{lr}_w \times 10$ and epoch size = 10,000. The course of parameter recovery during training is illustrated by the trajectories in the ($G$, $w$) plane (gray lines). Grey dots mark the end of training. Dashed lines indicate the homogeneous-model bifurcation reference. **B** Same as in **A** for FC fitting of homogeneous models (i.e., scalar $w$, $I$ and $\sigma$) on synthetic FC matrices derived from predefined ground-truth parameters, which were drawn from a grid of $\{G = 1, 4, 7\} \times \{w = -120, -80, -40, 0\} \times \{I = 1\} \times \{\sigma = 0.1\}$ (blue squares). Training was initialized at seeds of $\{G = 1\} \times \{w = -150{:}10{:}20\} \times \{I = 0\} \times \{\sigma = 10^{-5}\}$, with hyperparameters lr = 0.05, $\mathrm{lr}_w \times 10$ and epoch size = 10,000.

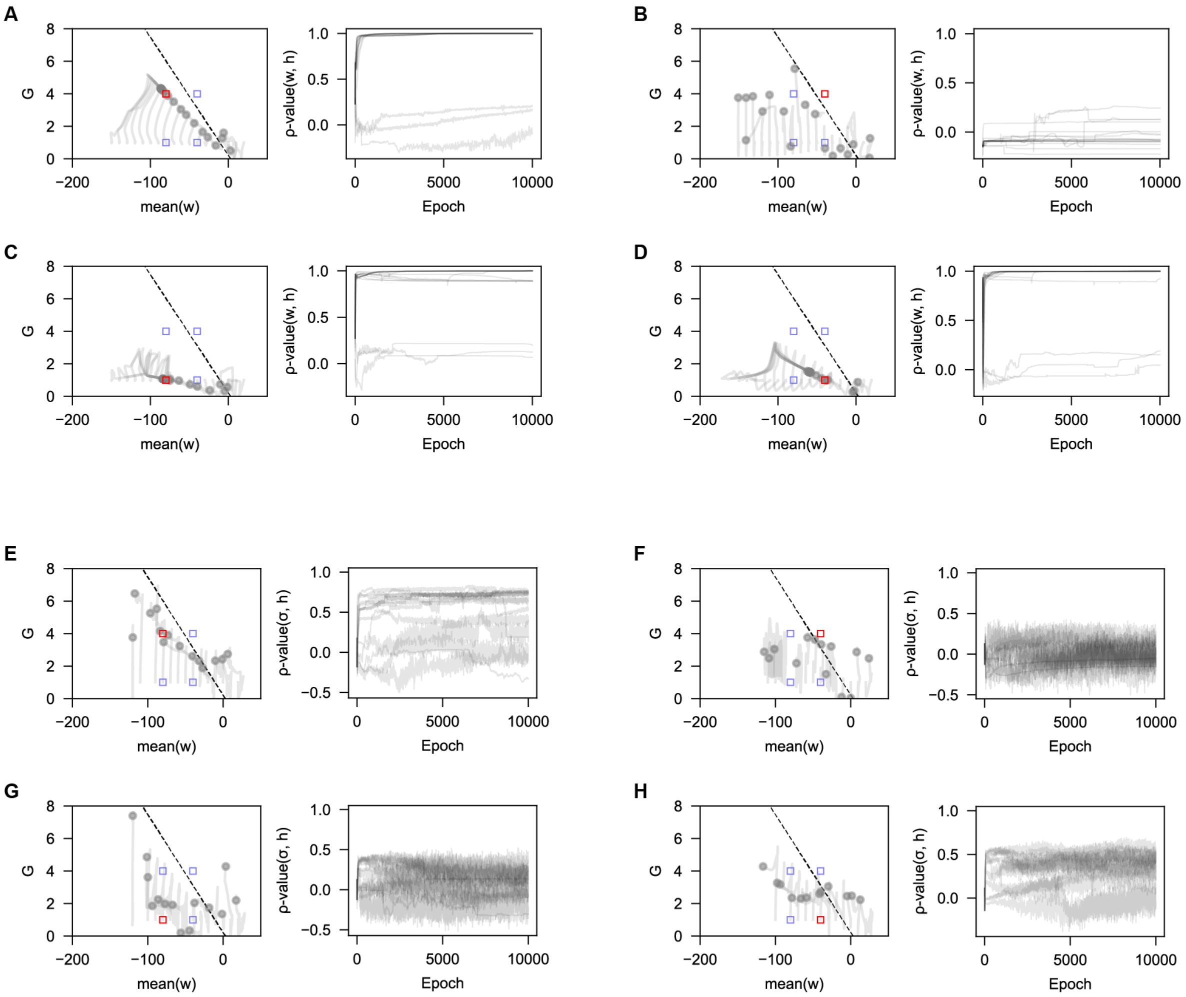

**Fig. S8: Parameter and heterogeneity recovery from synthetic connectomes. A–D** EC fitting of heterogeneous models (i.e., vector $\vec{w}$ and $\vec{I}$) on synthetic EC matrices derived from predefined ground-truth parameters. A vector $\vec{h}$ was generated with its elements sampled from Gaussian distribution, and normalized to zero mean and unit standard deviation, serving as a synthetic heterogeneity. The ground-truth parameters (red square) were sampled from a grid of $\{G = 1, 4\} \times \{w =$ -80, -40$\} \times \{I = 1\}$ (blue squares), with regional heterogeneity introduced into recurrent strength by the operation $\vec{w} \leftarrow w + 8 \times \vec{h}$. A synthetic EC matrix was generated using the ground-truth parameters as the training objective. Training was applied to optimize heterogeneous models as initialized homogeneously at seeds of $\{G = 1\} \times \{w = \text{-}150{:}10{:}20\} \times \{I = 0\}$, with hyperparameters lr = 0.05, $\mathrm{lr}_w \times 10$ and epoch size = 10,000. Left panel: the course of parameter recovery during training is illustrated by the trajectories in the ($G$, mean of $\vec{w}$) plane (gray lines). Grey dots mark the end of training. Dashed lines indicate the homogeneous-model bifurcation reference. Right panel: Spearman's $\rho$-value between vector $\vec{w}$ and $\vec{h}$ along the epochs during training. **E–H** Same as in **A–D** for FC fitting of heterogeneous models (i.e., vector $\vec{w}$, $\vec{I}$ and $\vec{\sigma}$) on synthetic FC matrices derived from predefined ground-truth parameters, which were sampled from a grid of $\{G = 1, 4\} \times \{w = \text{-}80, \text{-}40\} \times \{I = 1\} \times \{\sigma = 0.1\}$ (blue squares) with regional heterogeneity introduced into intrinsic noise by the operation $\vec{\sigma} \leftarrow \sigma + 0.1 \times \vec{h}$. Training was initialized at seeds of $\{G = 1\} \times \{w = \text{-}150{:}10{:}20\} \times \{I = 0\} \times \{\sigma = 10^{-5}\}$, with hyperparameters lr = 0.1, $\text{lr}_w \times 1$ and epoch size = 10,000.

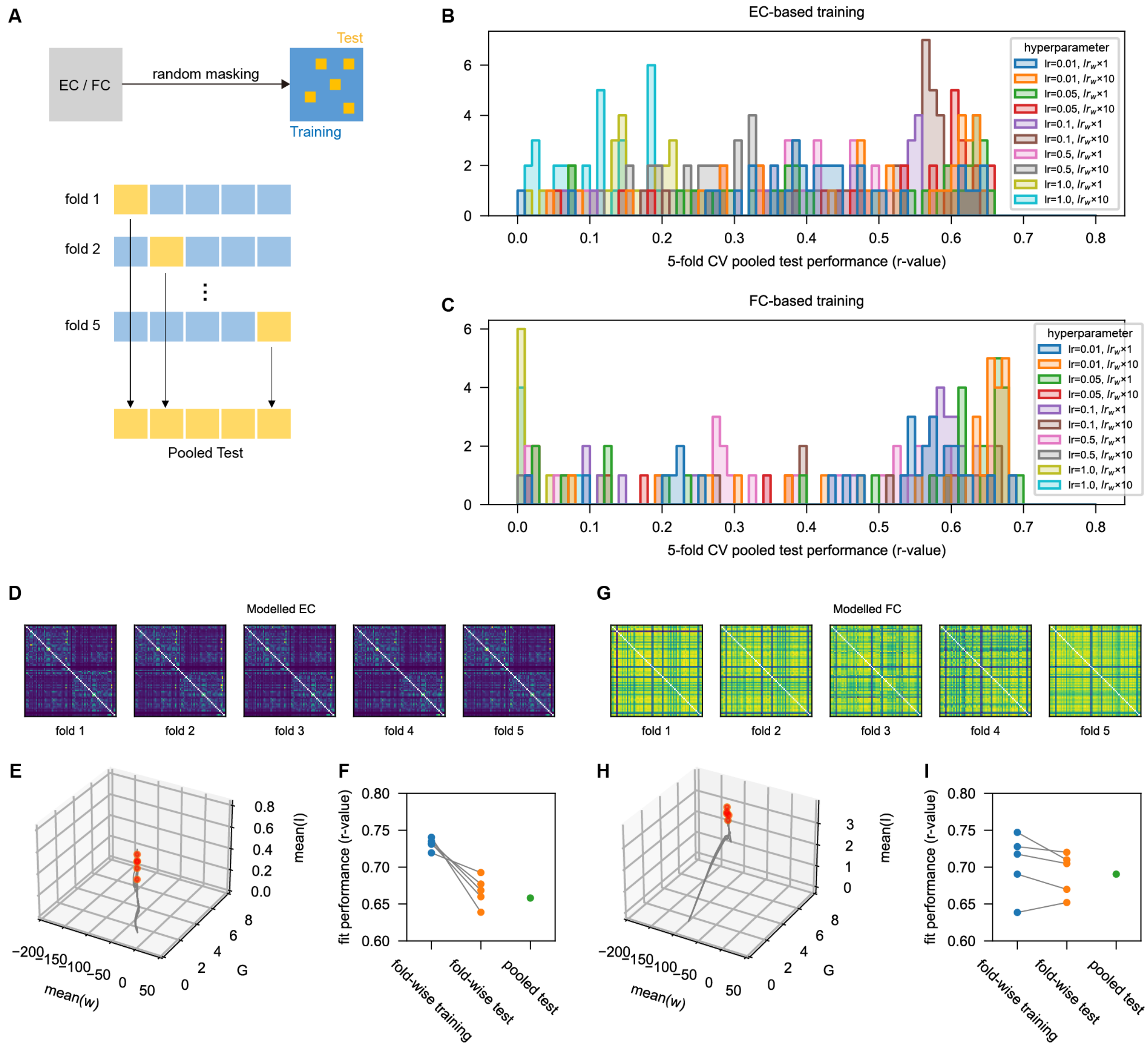

**Fig. S9: Five-fold cross-validation of EC fitting and FC fitting.** **A** Top panel: schematic of the random masking procedure for training and test set generation in a 5-fold cross validation (CV) design. Yellow squares represent masked entries forming the test set in each fold, and blue squares represent entries retained for training. Bottom panel: schematic illustrating the pooled test procedure, in which predictions of masked test entries across all folds are concatenated for an overall evaluation of fit performance, quantified by Pearson's $r$-value between concatenated predictions and empirical entries of the whole connectome. **B–C** Histograms displaying the distribution of pooled test performance across different hyperparameters for EC fitting (panel **B**) and FC fitting (panel **C**). Colors denote distinct hyperparameter configurations (lr, learning rate). **D–F** Example of the optimal case in 5-fold CV of EC fitting. The modelled EC matrices after training **D** and trajectories of parameters ($G$, mean of $w$, mean of $I$) during training **E** are illustrated for each fold. Fit performances (Pearson's $r$-value) are illustrated for training entries per fold, test entries per fold and the pooled test **F**. **G–I** Same as **D–F**, but for the optimal case in 5-fold CV of FC fitting.

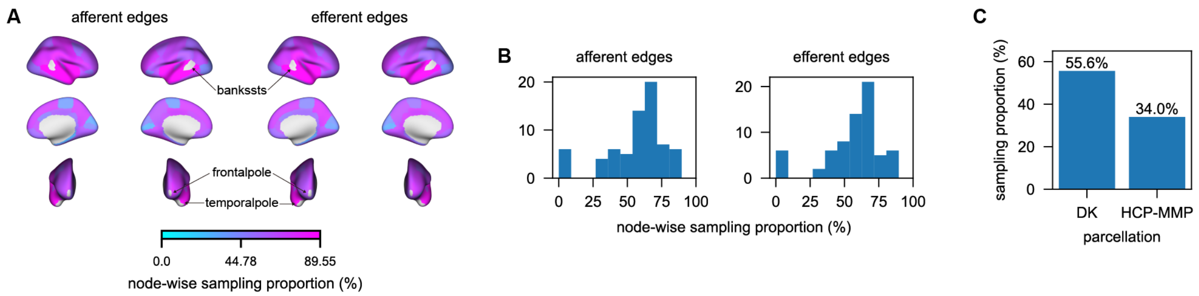

**Fig. S10: Sampling coverage of the empirical effective connectome. A** Cortical maps (DK parcellation) of node-wise sampling proportion, calculated as the ratio of measured edge number to total edge number for each region (node), either using afferent (i.e., along row axis of EC matrix; left panel) or efferent (i.e., along column axis of EC matrix; right panel) edges. 6 cortical regions (left and right bankssts, frontalpole and temporalpole) with sampling proportion = 0 are indicated, and excluded from node strength analysis involving EC. **B** Histograms showing the distribution of node-wise sampling proportions of afferent (left panel) and efferent (right panel) edges across all regions. **C** Overall sampling proportion of EC in DK and HCP-MMP parcellations, calculated as the ratio of the total number of measured edges to the total possible edges in the entire connectome. Overall coverage is 55.6% for DK and 34.0% for HCP-MMP.

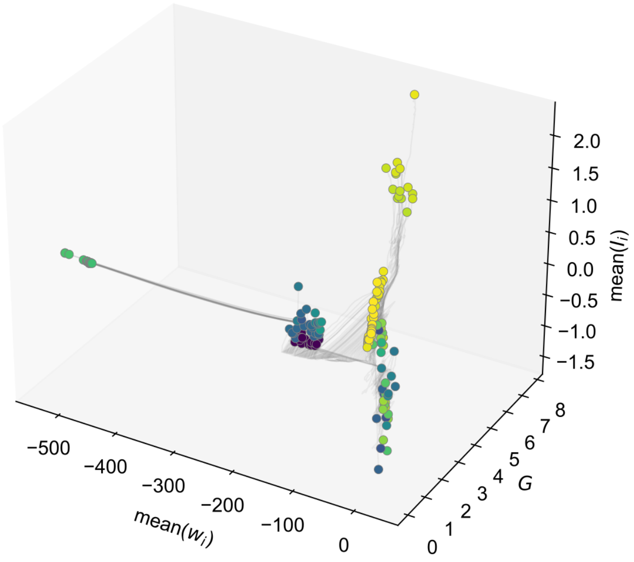

**Fig. S11: Three-dimensional parameter trajectories during EC fitting.** Trajectories are projected onto global coupling $G$, mean regional recurrent strength mean$(w_i)$ and mean background input mean$(I_i)$. Grey curves show individual training runs, and coloured circles mark their endpoints. Endpoint colours represent the final EC-fitting loss, using the convention of Fig. 4A. This full three-dimensional view complements the two-dimensional projection in Fig. 4A and retains trajectories beyond its displayed recurrent-strength range.

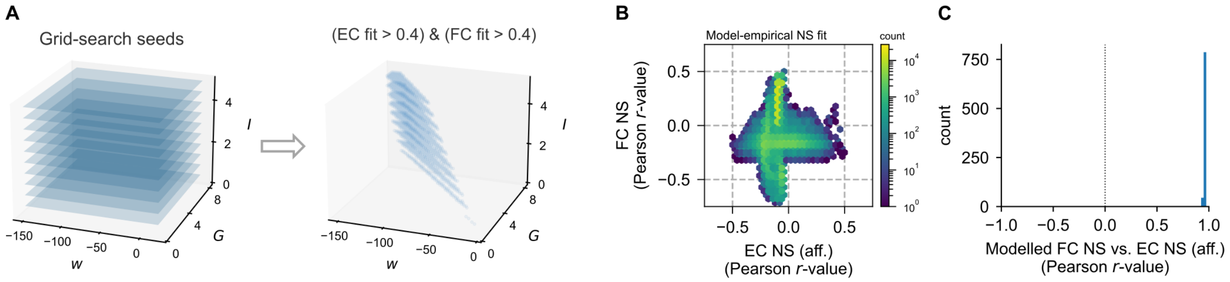

**Fig. S12: Homogeneous-model control for opposing EC and FC node-strength patterns. A** Four-dimensional grid search over homogeneous parameters $G \in \{0.2 : 0.2 : 8\}$, $w \in \{-150 : 2 : 20\}$, $I \in \{0.5 : 0.5 : 5\}$ and $\sigma \in \{0.5 : 0.5 : 5\}$ (344,000 combinations). Left: sampled parameter combinations projected onto the $(w, G, I)$ axes. Right: retained models with Pearson model-empirical fits greater than 0.4 for both EC and FC. **B** Joint distribution of model-empirical afferent EC NS and FC NS fits in all grid-searched homogeneous models. Colour indicates the number of models per bin on a logarithmic scale. **C** Distribution of Pearson correlations between modelled FC NS and afferent EC NS in retained homogeneous models. The dotted line marks zero. These models yield predominantly positive EC–FC NS correlations.

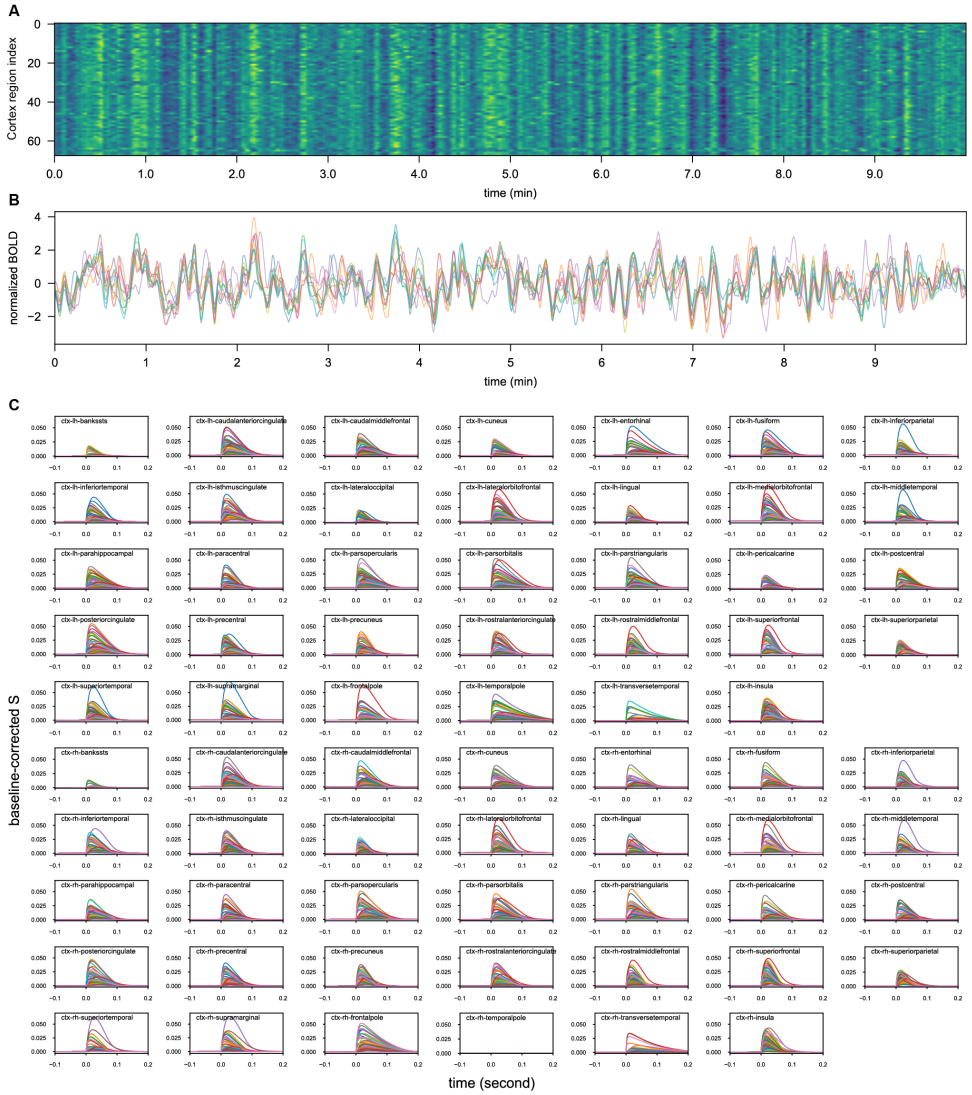

**Fig. S13: Resting-state and stimulus-evoked time courses of the optimal model. A** Heatmap of modelled BOLD signals across cortical regions over a 10-minute virtual resting-state simulation. BOLD signals are normalized to zero mean and unit standard deviation. **B** Representative time courses of normalized BOLD signals from 10 selected cortical regions, extracted from **A**. **C** modelled time courses of baseline-corrected responses (i.e., $S$ – baseline of $S$) under virtual single-pulse stimulation. The stimulated target region is indicated in each panel.

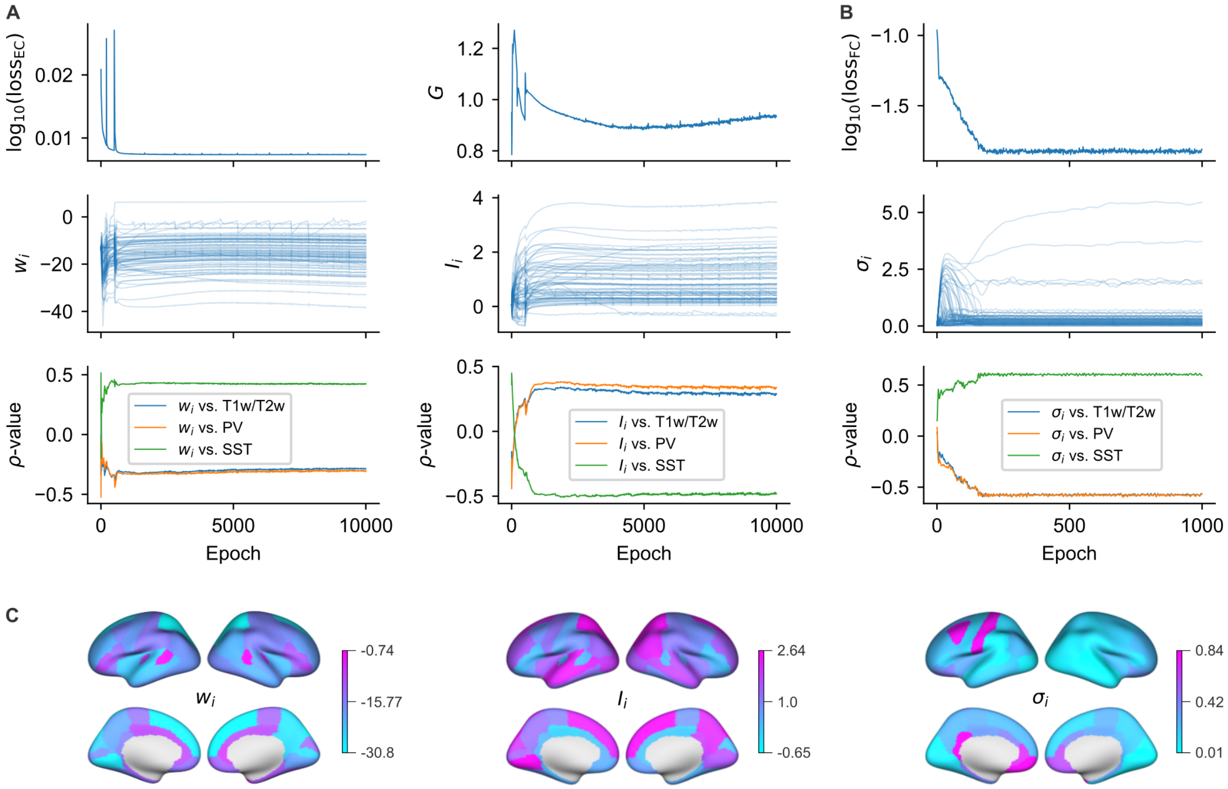

**Fig. S14: Emergence of regional parameter heterogeneity during training of the optimal model.** **A** Evolution during the 10,000-epoch EC-fitting stage. Upper: EC-fitting objective (left) and global coupling $G$ (right). Middle: regional recurrent strengths $w_i$ (left) and background inputs $I_i$ (right). Lower: Spearman correlations of $w_i$ (left) and $I_i$ (right) with T1w/T2w (blue), PV (orange) and SST (green). **B** Evolution during the subsequent 1,000-epoch FC-fitting stage, in which only $\sigma_i$ is optimized. Upper: FC-fitting objective. Middle: regional noise amplitudes $\sigma_i$. Lower: Spearman correlations of $\sigma_i$ with the same cortical gradients. Thin parameter traces in **A,B** correspond to individual regions. **C** Cortical maps of the final $w_i$, $I_i$ and $\sigma_i$ of the optimal model. Each map has its own parameter-value colour scale.

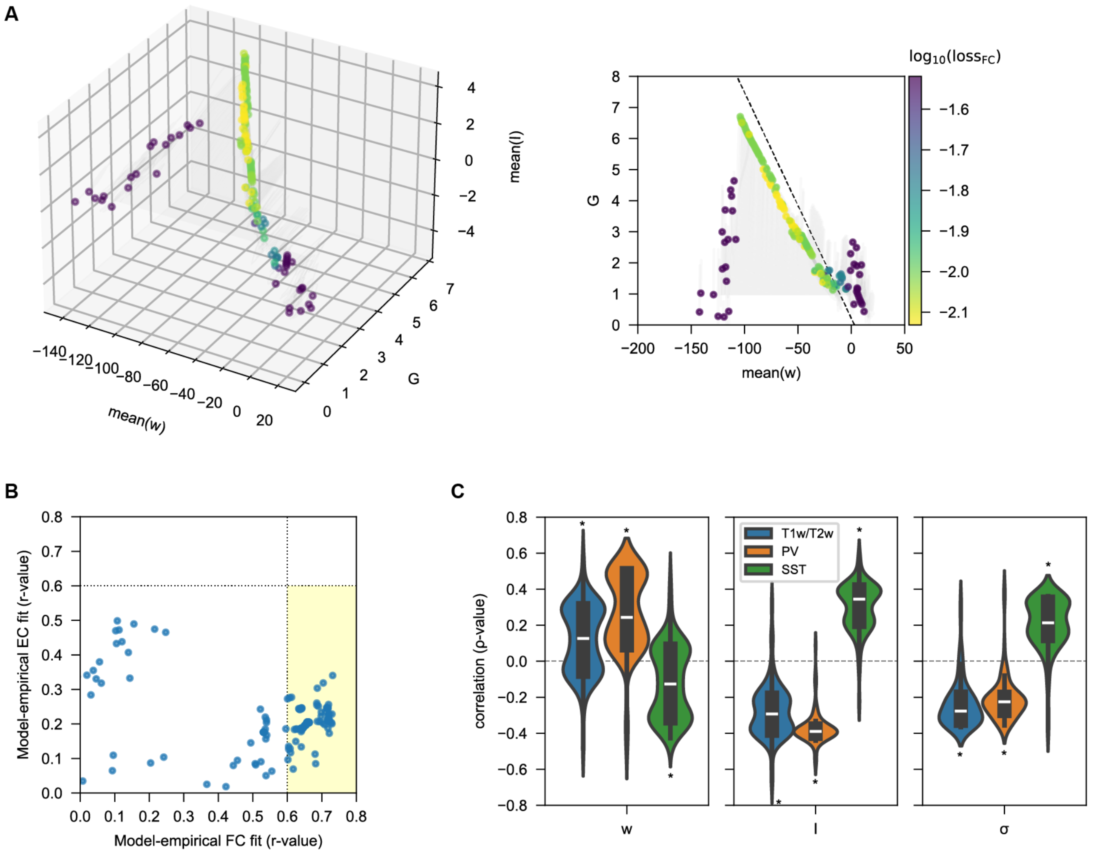

**Fig. S15: Outcomes of single-stage FC-only training.** **A** Left panel: 3D plot of trajectories of parameters during FC fitting, with axes representing $G$, the mean of $\vec{w}$ and the mean of $\vec{I}$. Right panel: 2D projection of trajectories to the plane of $G$ and the mean of $\vec{w}$. The dashed line indicates the homogeneous-model bifurcation reference. End of training is indicated by the scatter plot as color-coded by $\log_{10}(\text{loss}_{\text{FC}})$. **B** Scatter plot illustrates the fitting performances after training according to the Pearson's $r$-value between empirical and modelled connectomes. Shaded area highlights the selected models with Pearson's $r$-value > 0.6 for FC fitting. **C** Violin plots depicting the distribution of Spearman's $\rho$-value between $\vec{w}$, $\vec{I}$ and $\vec{\sigma}$ of the selected models in **B** and T1w/T2w ratio, PV and SST (*, FWE-corrected p < 0.05, one sample t-test).

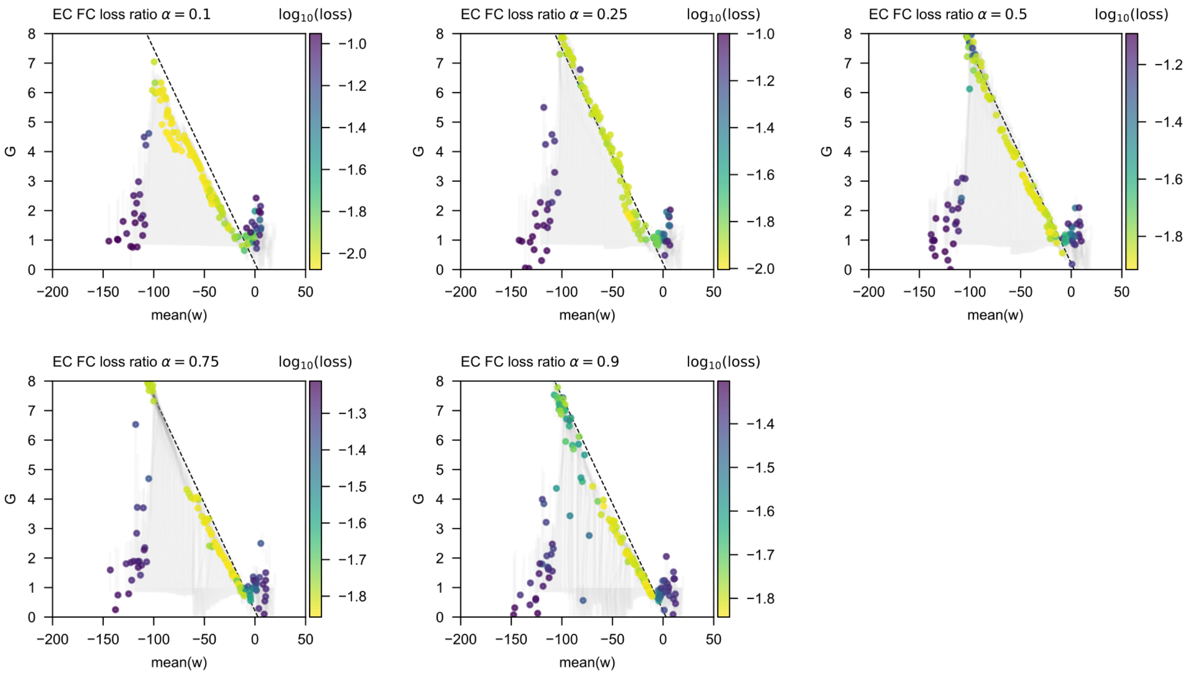

**Fig. S16: Training trajectories under a hybrid EC–FC loss.** Training minimizes $\mathcal{L}_{\text{hybrid}} = \alpha\mathcal{L}_{\text{EC}} + (1-\alpha)\mathcal{L}_{\text{FC}}$, with loss ratio $\alpha = 0.1, 0.25, 0.5$ (top row, left to right), 0.75 and 0.9 (bottom row). Each condition uses Adam with learning rate 0.05, batch size 68 and 10,000 epochs. Grey curves show trajectories projected onto $G$ and mean regional recurrent strength. Coloured circles indicate endpoints and encode $\log_{10}(\mathcal{L}_{\text{hybrid}})$ using the colour scale in each panel. Dashed lines indicate the homogeneous-model bifurcation reference. The corresponding fit performances and parameter–hierarchy correlations are shown in main-text Fig. 6C.

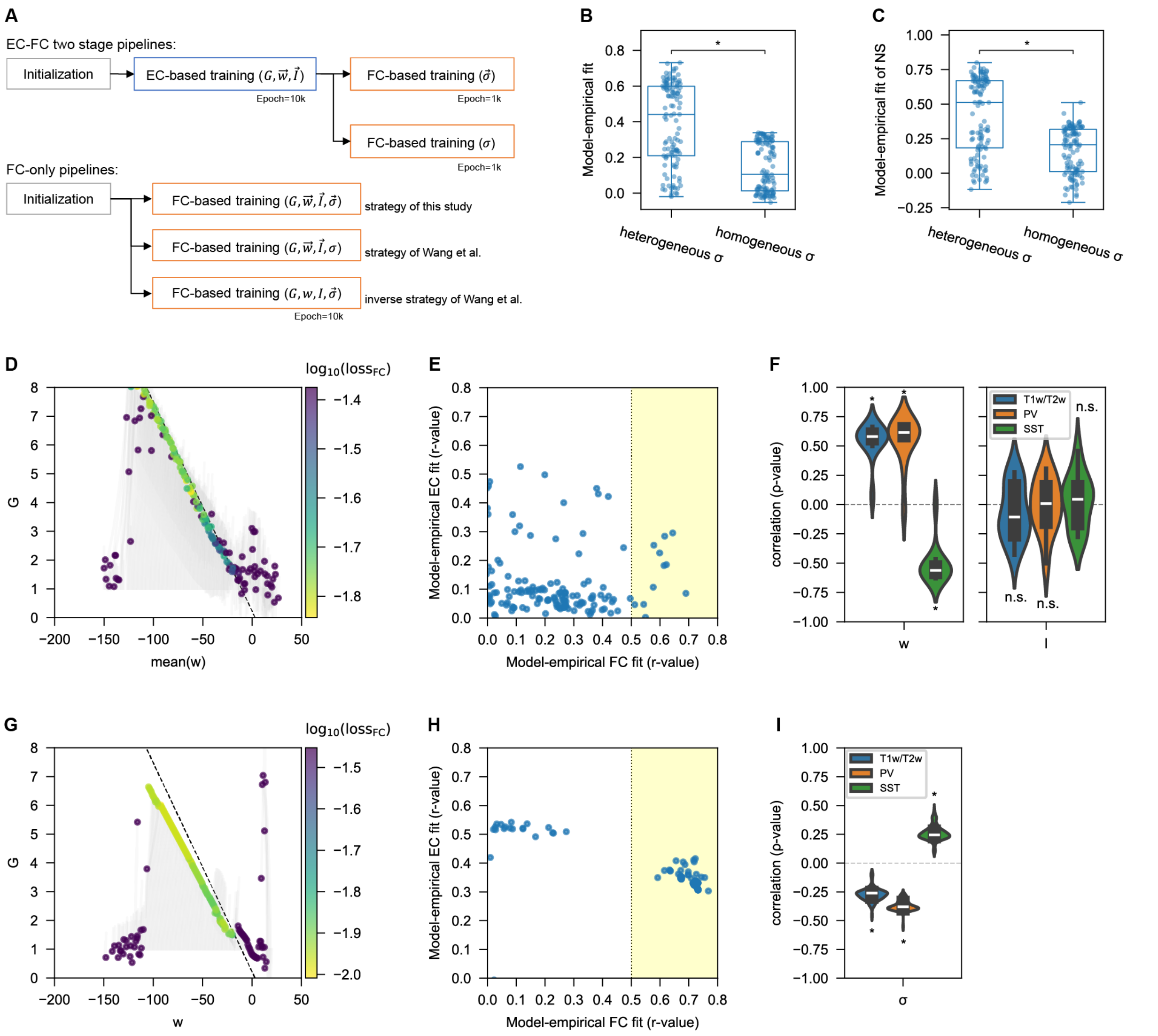

**Fig. S17: Training with partially homogeneous and partially heterogeneous parameters. A** Schematic overview of two-stage EC–FC training and FC-only training with mixed homogeneous/heterogeneous parameterization. Vector symbols denote heterogeneous parameters trained independently across cortical regions, whereas scalar symbols indicate homogeneous parameters constrained to be identical across regions. Top panel: two-stage training begins with EC fitting to optimize $G$, $\vec{w}$ and $\vec{I}$, followed by FC fitting of heterogeneous $\vec{\sigma}$. A control condition constrains $\sigma$ to be homogeneous. Bottom panel: FC-only training strategy of this study with heterogeneous $\vec{w}$, $\vec{I}$ and $\vec{\sigma}$; the strategy of Wang et al. (*14*) with heterogeneous $\vec{w}$ and $\vec{I}$ but homogeneous $\sigma$; an inverse strategy of Wang et al. with heterogeneous $\vec{\sigma}$ but homogeneous $w$ and $I$. **B–C** Boxplots comparing model–empirical fit performance between heterogeneous $\vec{\sigma}$ and homogeneous $\sigma$ conditions under two-stage training (*p $< 10^{-5}$, two-sample t-test). Model-empirical FC fit at the edge level (panel **B**) or node-strength level (NS; panel **C**) is quantified using Pearson's $r$-value. **D–F** FC-only training following the strategy of Wang et al. with heterogeneous $\vec{w}$ and $\vec{I}$ but homogeneous $\sigma$. **D** Training trajectories projected onto the plane of $G$ and mean of $\vec{w}$. **E** Scatter plot of model-empirical EC/FC fit after training. Well-trained models are thresholded in the shaded region (r > 0.5 for FC fit). **F** For these well-trained models, violin plots show the distribution of Spearman's $\rho$-value between heterogeneous parameters ($\vec{w}$, $\vec{I}$) and cortical gradients. Significance of deviation from zero was assessed using one-sample t-test (*, FWE-corrected p < 0.05; n.s., not significant). **G–I** Same as **D–F**, but using the inverse strategy of Wang et al., where **G** shows training trajectories projected onto the plane of $G$ and homogeneous $w$.

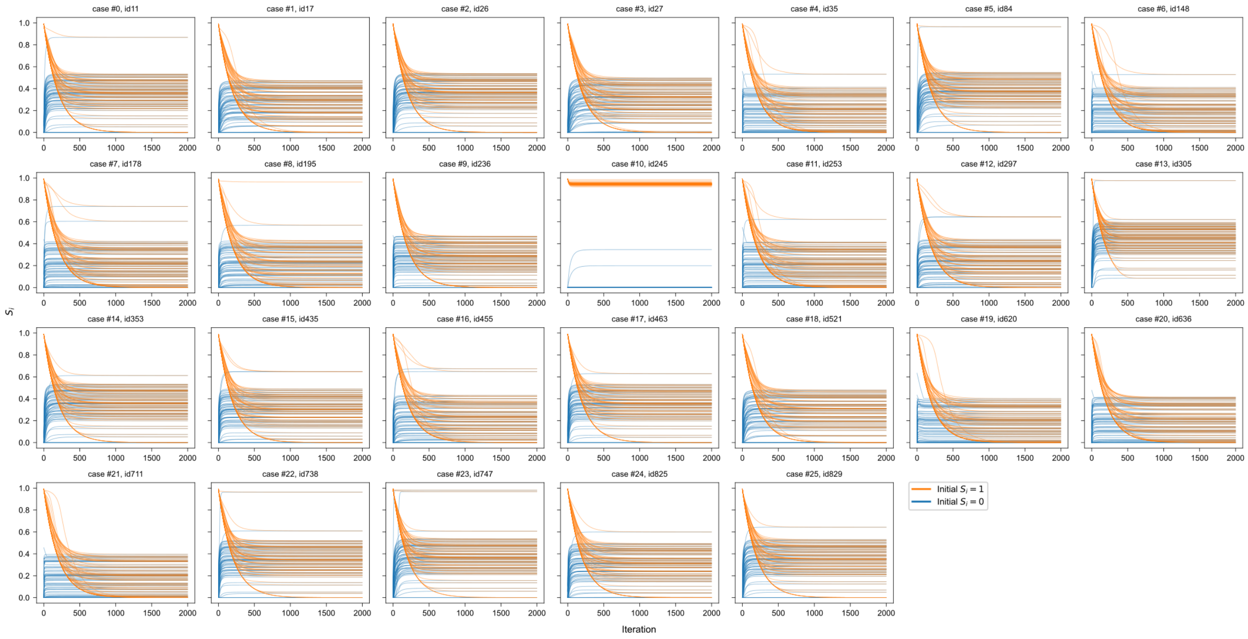

**Fig. S18: Noise-free trajectories toward equilibrium in the 26 retained models.** Each panel shows the synaptic-gating variables $S_i$ of all 68 cortical regions in one retained model, using its final trained parameters. Blue and orange lines show trajectories initialized with $S_i = 0$ and $S_i = 1$ in every region, respectively; each line represents one region. Stochastic input and external perturbations are absent. The horizontal axis shows 2,000 integration steps at $\Delta t = 0.5$ ms, corresponding to 1 s; the vertical axis shows $S_i$. Except for case #10 which exhibits bistability, trajectories from the two initial conditions converge to the same stable equilibrium in all other models. The optimally trained model is case #13.

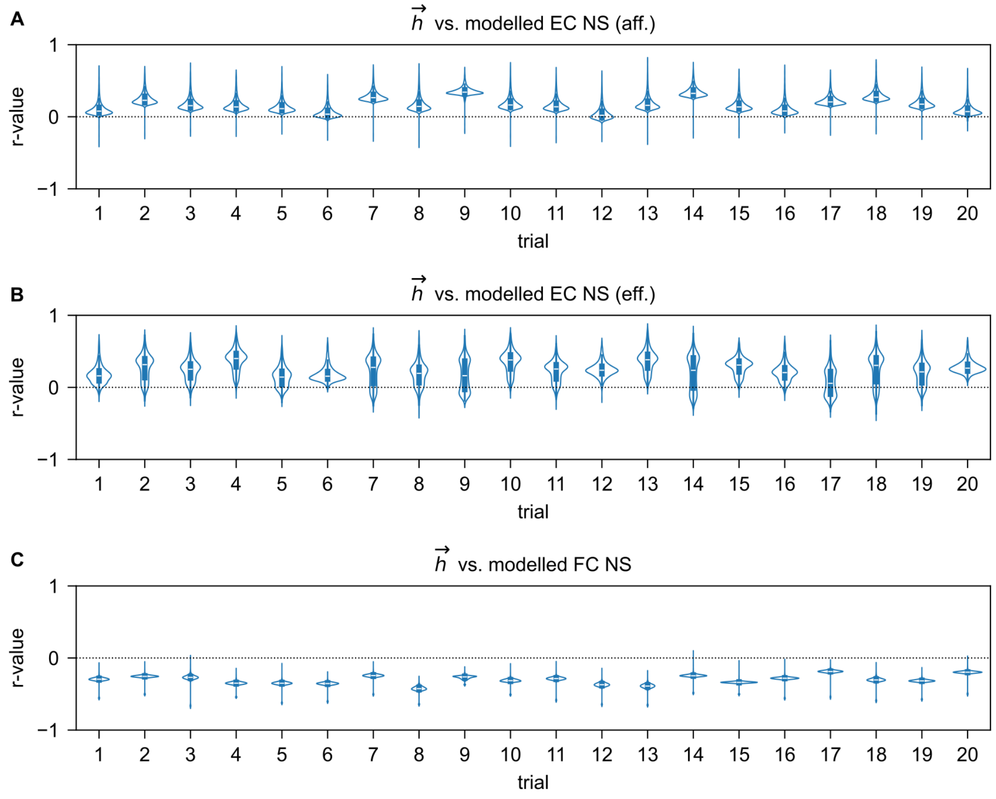

**Fig. S19: Robustness of heterogeneity-induced node-strength changes across random gradients. A–C** Violin plots show the distribution of Pearson's correlation coefficients ($r$-values) between a randomly generated heterogeneity vector $\vec{h}$ and modelled node strengths (NS) at the final heterogeneity-introduction step across 20 independent trials, with each trial following the same procedure in Fig. 8A,B. Trial number is shown on the x-axis, with NS calculated using afferent (aff.) and efferent (eff.) EC and FC in panel **A**, **B** and **C**, respectively.

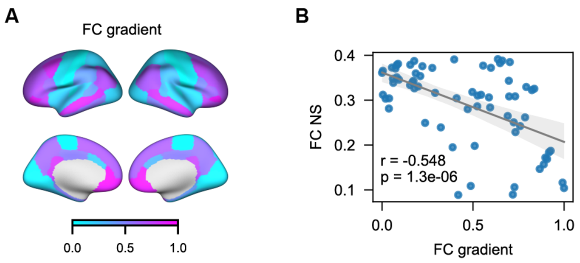

**Fig. S20: Relationship between the principal FC gradient and FC node strength. A** Cortical map showing the regional distribution of the FC principal gradient from (*15*). **B** Scatter plot comparing the FC principal gradient and FC node strength (NS). The gray line denotes the linear regression fit, with the corresponding Pearson's correlation coefficient and *p*-value shown. The FC principal gradient, derived using nonlinear dimensionality reduction via diffusion embedding, captures a dominant axis of variation in the FC matrix and is an established marker of cortical hierarchy (*18*). FC node strength is significantly negatively correlated with this gradient, suggesting that node strength reflects the low-rank structure of FC and is negatively aligned with cortical hierarchy.

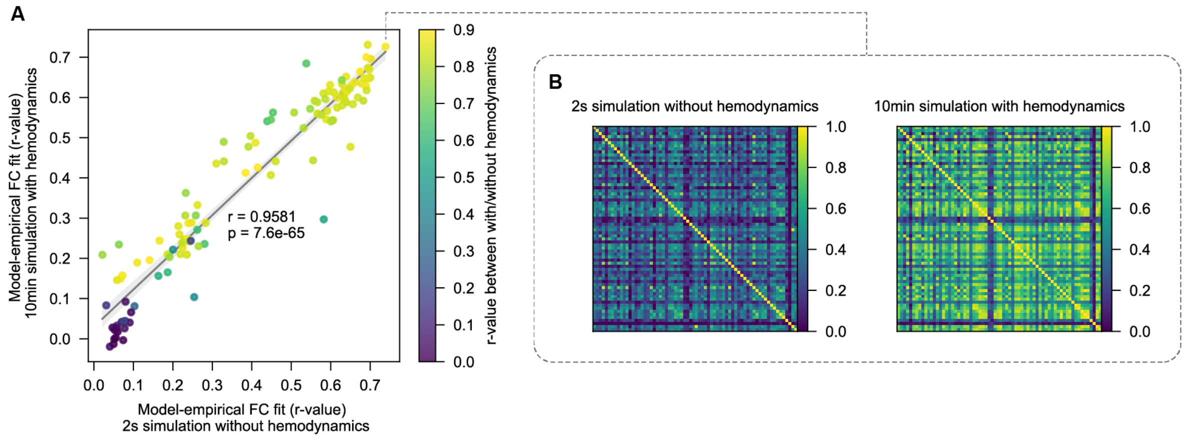

**Fig. S21: Modelled FC with or without the Balloon-Windkessel hemodynamic model. A** Scatter plot comparing model-empirical FC fits, i.e., $r$-value of upper triangular entries between modelled and empirical FC matrices, between 2s simulation without hemodynamics (x-axis) and 10 min simulation with hemodynamics (y-axis). The gray line denotes linear regression fit with the corresponding Pearson's correlation coefficient and statistical significance. The colormap indicates the $r$-value of upper triangular entries between modelled FC matrices with and without hemodynamics. The MFMs use the parameters obtained through two-stage training. **B** Illustration of modelled FC matrices for a selected case from **A**, derived from a 2s simulation without hemodynamics (left panel) and a 10 min simulation with hemodynamics (right panel).

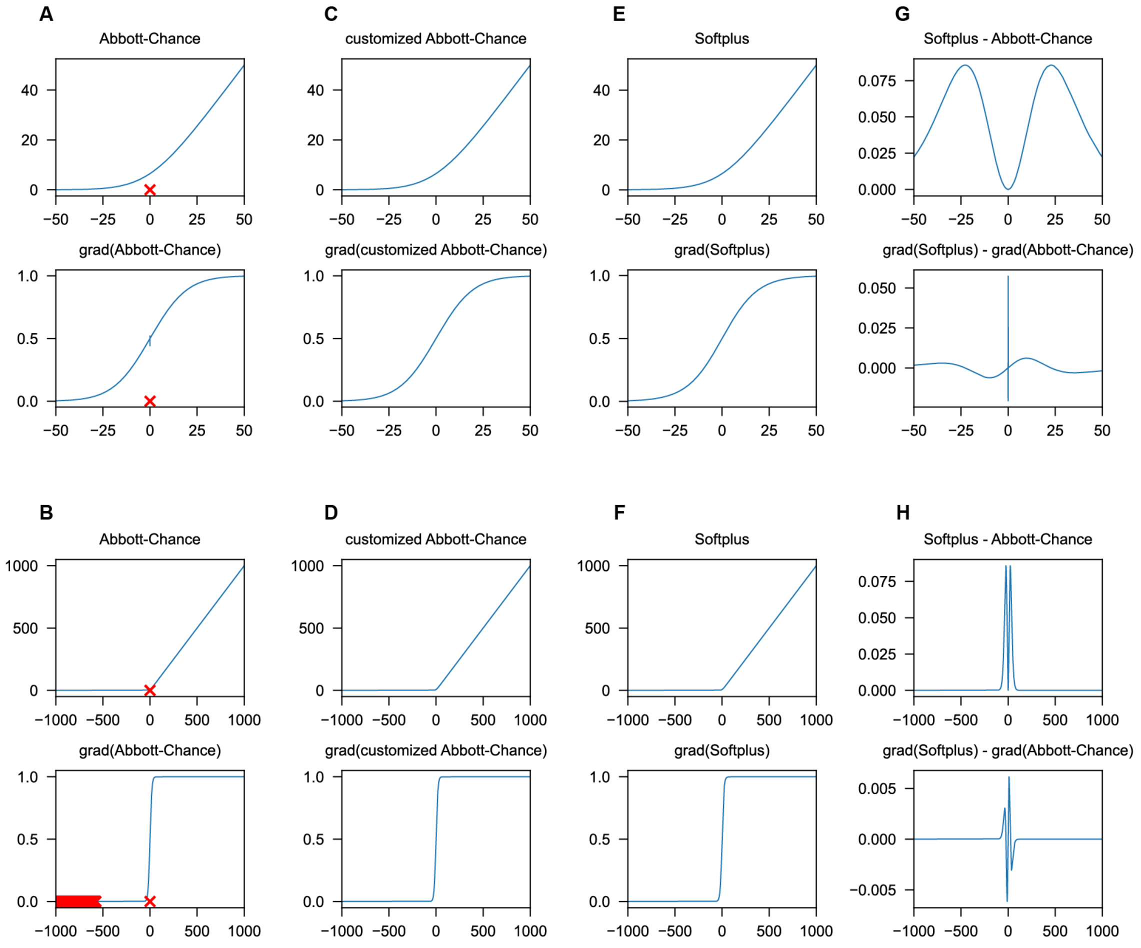

**Fig. S22: Input-output curves for activation functions. A,B** Abbott-Chance function (top panel) and its gradient (bottom panel) over narrow (±50; A) and wide (±1000; B) input ranges. **C,D** Same plots as **A,B** for the customized Abbott-Chance function. **E,F** Same plots as **A,B** for Softplus function. **G,H** Point-wise differences, Softplus minus Abbott-Chance (top panel), and the corresponding gradient differences (bottom panel) over narrow (±50; G) and wide (±1000; H) input ranges. The parameter settings: $d = 0.154$ for Abbott-Chance and $\beta = \ln(2) \times d$ for Softplus. Red crosses indicate the points with NaN return. All gradients were computed using automatic differentiation in BrainPy.
